# Comparative gene expression in cells competent in or lacking DNA-PKcs kinase activity following etoposide exposure reveal differences in gene expression associated with histone modifications, inflammation, cell cycle regulation, Wnt signaling, and differentiation

**DOI:** 10.1101/2020.09.17.300129

**Authors:** Sk Imran Ali, Mohammad J. Najaf-Panah, Johnny Sena, Faye D. Schilkey, Amanda K. Ashley

## Abstract

Maintenance of the genome is essential for cell survival, and impairment of the DNA damage response is associated with multiple pathologies including cancer and neurological abnormalities. DNA-PKcs is a DNA repair protein and a core component of the classical non-homologous end-joining pathway, but it may have roles in modulating gene expression and thus, the overall cellular response to DNA damage. Using cells producing either wild-type (WT) or kinase-inactive (KR) DNA-PKcs, we assessed global alterations in gene expression in the absence or presence of DNA damage. We evaluated differential gene expression in untreated cells and observed differences in genes associated with cellular adhesion, cell cycle regulation, and inflammation-related pathways. Following exposure to etoposide, we compared how KR versus WT cells responded transcriptionally to DNA damage. Downregulation of pathways involved in biosynthesis were observed in both genotypes, but upregulated biological pathways were divergent, again with KR cells manifesting a more robust inflammatory response compared to WT cells. To determine what major transcriptional regulators are controlling the differences in gene expression noted, we used pathway analysis and found that many master regulators of histone modifications, proinflammatory pathways, cell cycle regulation, Wnt/β-catenin signaling, and cellular development and differentiation were impacted by DNA-PKcs status. Overall, our results indicate that DNA-PKcs, in a kinase-dependent fashion, decreases proinflammatory signaling following genotoxic insult. As multiple DNA-PK kinase inhibitors are in clinical trial as cancer therapeutics utilized in combination with DNA damaging agents, understanding the transcriptional response when DNA-PKcs cannot phosphorylate downstream targets will inform the overall patient response to combined treatment.

## Introduction

Eukaryotic cells frequently experience genotoxic stress from various exogenous and endogenous sources, including exposure to UV radiation, reactive oxygen/nitrogen species formed during metabolism, genotoxic chemicals, and others, inducing myriad types of DNA damage (1). DNA double strand breaks (DSB) are highly deleterious, which if unrepaired or misrepaired can cause genomic instability associated with multiple diseases, including cancer. In response to DSB, cells initiate a complex, coordinated set of pathways, collectively known as DNA damage response (DDR) involving damage sensing, transferring signals via signal transducers and many effector proteins to respond to damage. Ideally, the DDR serves to repair DNA damage when possible through activation of repair proteins and upregulating transcription of additional genes needed for a full response. However, if repair is not feasible, the DDR initiates apoptotic death. In eukaryotes the majority of DSB are repaired by the non-homologous end-joining (NHEJ) and homologous recombination (HR) pathways. The DNA dependent protein kinase catalytic subunit (DNA-PKcs), a member of the phosphatidyl inositol 3-kinase related kinase (PIKK) family of proteins including ATM and ATR, is directly involved in NHEJ-mediated repair (2). DNA-PKcs is recruited to DSB by the Ku 70/80 heterodimer, forming the catalytically active DNA-PK holoenzyme, which in turn results in DNA-PKcs autophosphorylation and phosphorylation of multiple other proteins (Ku, H2AX, RPA32). Additionally, DNA-PKcs serves to tether the two broken DNA ends (3) and activates other NHEJ proteins (XRCC4, XLF, Lig4 or PAXX) to facilitate DNA end processing and ultimately ligation (4, 5). DNA-PKcs regulates proteins involved in HR, including ATM, WRN, RPA32, cAbl and SMC1, facilitating crosstalk between the DDR proteins and two dominant DSB repair pathways (2, 6–11). DNA-PKcs phosphorylates KAP-1 immediately after DSB and promotes chromatin decondensation to facilitate recruiting DDR proteins (12). Clearly, DNA-PKcs has multifaceted roles in regulating the cellular response to DSB.

DNA-PKcs is dually involved in regulating proteins involved in cellular transcription. DNA-PKcs phosphorylates Snail1, a zinc-finger transcription factor involved in the epithelial-to-mesenchymal transition, which promotes its stability and function potentiation (8, 13). DNA-PKcs is also involved in the activation of several metabolic genes. DNA-PKcs activation in hypoxic condition protects HIF1α from degradation which promotes one of its target gene’s, GluT1, expression (14). DNA-PKcs promotes fatty acid biosynthesis in an insulin-dependent manner by activating transcription factor USF-1 (15). During aging, DNA-PKcs phosphorylation of heat-shock protein HSP90α decreases the mitochondrial function in skeletal muscles, thus decreasing overall metabolism and fitness (16). DNA-PKcs was originally isolated as a component of SP1 transcriptional complex (17) as well as co-eluted with the largest subunit of RNA polymerase II (RNAPII) (18). DNA-PKcs phosphorylates RNAPII (19) as well as many other transcription factors TFIIB, SP1, Oct1, c-Myc, c-Jun and TATA binding protein (TBP) (20). The availability of DNA-PKcs at the active transcription sites modulates the function of many transcription factors like, autoimmune regulator (AIRE), p53, erythroblast transformation-specific related gene (ERG) by direct or indirect interactions (13). DNA-PKcs may mediate transcription initiation by interacting with TopoIIβ and PARP1 in the presence of DSB (21, 22). The effects of DNA-PKcs on global gene expression are yet unknown. In this study, we analyzed global gene expression with and without DNA damage in cell lines expressing wild-type or kinase-inactive DNA-PKcs, to elucidate the roles DNA-PKcs has in regulating gene expression in kinase-dependent and -independent manners alone or following damage.

## Materials and Methods

### Cells culture

V3-derived Chinese Hamster Ovary (CHO) cell lines were kindly provided by Dr. Katherine Meek, complemented with either human wild-type (WT) or kinase inactive (K3753R) DNA-PKcs (KR) (23). Cells were cultured in complete media: α-MEM (LifeTechnologies, Waltham, MA) supplemented with 10% FBS (MilliporeSigma, St. Louis, MO), 1% penicillin/streptomycin (LifeTechnologies), 200 μg/mL G418 (LifeTechnologies) and 10 μg/ml puromycin (Santa Cruz Biotechnology, Santa Cruz, CA) at 37°C with 5% CO_2_ and 100% humidity. All chemicals were purchased from MilliporeSigma unless otherwise indicated.

### Immunoblotting

Cells cultured to 80-90% confluence and whole cell lysate were prepared using RIPA buffer (50mM Tris (pH 7.4), 2mM EDTA, 150mM NaCl, 0.1% SDS, 1.0% Triton X-100) supplemented with Halt phosphatase and protease inhibitor cocktails (Fisher Scientific, Waltham, MA). Protein was quantified using the Pierce BCA Protein Assay (Fisher Scientific) per manufacturer’s instructions. Immunoblotting was used to assess the production of DNA-PKcs in all cell lines. Briefly, 25μg of protein was subjected to SDS-PAGE, transferred to PVDF membrane, and blocked with 5% nonfat dried milk in 1X TBS-T (Tris buffer saline with 0.1% Tween-20) for 1 h at room temperature (RT), then primary antibody recognizing DNA-PKcs (Abcam, catalog # 70250) was added overnight at 4°C in blocking buffer. Membranes were washed, then a HRP-conjugated secondary antibody (Jackson Immuno Research, catalog # 111035144, West Grove, Pennsylvania) in blocking buffer was added for 1h at RT. DNA-PKcs protein was assessed using Clarity Western ECL Substrate (BioRad, Hercules, CA) and captured using a Chemi Doc MP Imaging System (BioRad). Total vinculin levels (Santa Cruz, catalog # 73614) were assessed after DNA-PKcs analysis following the same procedure.

### Clonogenic Survival Assay

We assessed clonogenic survival as previously described (24). Cells were plated onto 6 wells plates at 150 cells per well and allowed to attach for 4 hours and then treated with etoposide (0.01 to 100 μM) or DMSO. After 24 hours incubation at 37°C the treatment containing media was removed and the cells were re-incubated for 5 to 7 days with fresh media to allow colony formation. Surviving colonies were fixed using 4% paraformaldehyde and stained with 0.5% crystal violet in 20% methanol, washed with water and air dried. Colonies with ≥ 50 cells were counted. Plating efficiency (PE) and surviving fraction (SF) were calculated: PE = (number of colonies formed divided by the number of cells seeded) ×100; SF = PE of treated cells divided by PE of control cells. Data is the average of multiple assays which each contained three experimental replicates.

### Cell Viability

Cells were cultured in complete media in white-walled 96-well plates (CoStar, Corning, NY) for 24 h and then treated with increasing concentrations of etoposide or DMSO and incubated for 72h. Viability was quantified using the CellTiter-Glo Assay (Promega, Madison, WI) per manufacturer’s instructions.

### RNA Isolation

Cells were cultured on 100 mm dishes overnight in complete medium and then treated with DMSO or 20 μM etoposide for 24 h. Total RNA was isolated using Trizol reagent (MilliporeSigma) isolation was according to manufacturer’s instructions. RNA was dissolved in nuclease-free water followed by spectrophotometric analysis of the RNA quantity; RNA was subjected to electrophoresis on an agarose gel to assess RNA quality and purity.

### Transcriptome Library Preparation and Sequencing

RNA libraries were constructed strictly according to [kit or protocol credit needed] user’s guide. The oligo (dT) beads were used to purify poly-A mRNAs then fragmented in fragmentation buffer. The first-strand complementary DNA was synthesized using random-hexamer primers. The second-strand cDNA was polymerized using dNTPs, RNase H, DNA polymerase I, and buffer. After DNA library quantification and validation steps, they amplified by PCR. The sequencing of amplified library was conducted through Illumina paired-end runs (100 bp long reads) on HiSeq™ 2000 technology (Illumina Inc, USA).

### Differentially expressed gene analysis

The low quality raw reads (fastq format) filtered based on Q30 and GC content, then the Illumina adaptors were trimmed of reads for minimum read length of 36 bases using *Trimmomatic* v0.34 (25). The index of the *Chinese hamster* reference genome (CHOK1GS_HDv1) was built using HISAT2 v2.1.0.(26). Hisat2 v2.2.1 tool was utilized to align the reads to the genome sequences in FASTA format and output aligned reads in binary alignment map (BAM) format were translated into the transcriptomes of each sample using stringtie v2.0 tool which uses a novel network flow algorithm as well as an optional de novo assembly step to assemble and quantitate full-length transcripts representing multiple splice variants for each gene locus (27). The stringtie outputs (GTF files) were merged to create a single master transcriptome GTF with exact same naming and numbering scheme across all transcripts. The *feature Counts* tool implemented under *subread v2.0* was utilized to quantify transcripts assembled by stringtie mapped to each gene (28). Eventually, the differentially expressed (DE) gene profiles were statistically analyzed through *edgeR* and *limma* R libraries (29, 30).

### Biological Pathway Analyses

The ClusterProfiler R tool was applied on the to demonstrate the functional pathways and gene network enrichment analysis (31). This analysis provides information related to the biological pathways significantly enriched in up- or down-regulated genes through mediating the Kyoto Encyclopedia of Genes and Genomes (KEGG) database and gene ontology (GO) terms (using a standard false discovery rate (FDR) < 0.05 and p-value cutoff 0.05).

### Regulatory Network Analyses

The upstream regulatory network analysis was performed through Ingenuity Pathway Analysis (IPA, QIAGEN) platform. The IPA casual network approach was applied on contrast matrices, by having gene symbol, log fold change, and p-value<0.05 of each gene, to characterize the upstream regulators as well as master regulator (the root of network) those responsible for driving a set of target genes (32).

### Statistical analysis

Cell viability was analyzed using the non-linear regression function in GraphPad Prism (version 8) comparing viability as a function of the log etoposide dose. The area under the curve (AUC) for viability and colony forming assays was assessed using GraphPad Prism (version 7; San Diego, CA), and the resultant total peak AUC and standard error per sample were compared using a one-way ANOVA with an *ad hoc* Tukey’s multiple comparisons test.

## Results

### DNA-PKcs promotes survival following exposure to etoposide

We confirmed that DNA-PKcs is produced in our WT and KR cells (Figure 1A). We assessed the effect of DNA-PKcs on cell survivability after exposure to the topoisomerase II inhibitor, etoposide. Consistent with previous reports, cells lacking DNA-PKcs kinase activity are more sensitive to etoposide (Figure 1B – E).

**Figure 1:**
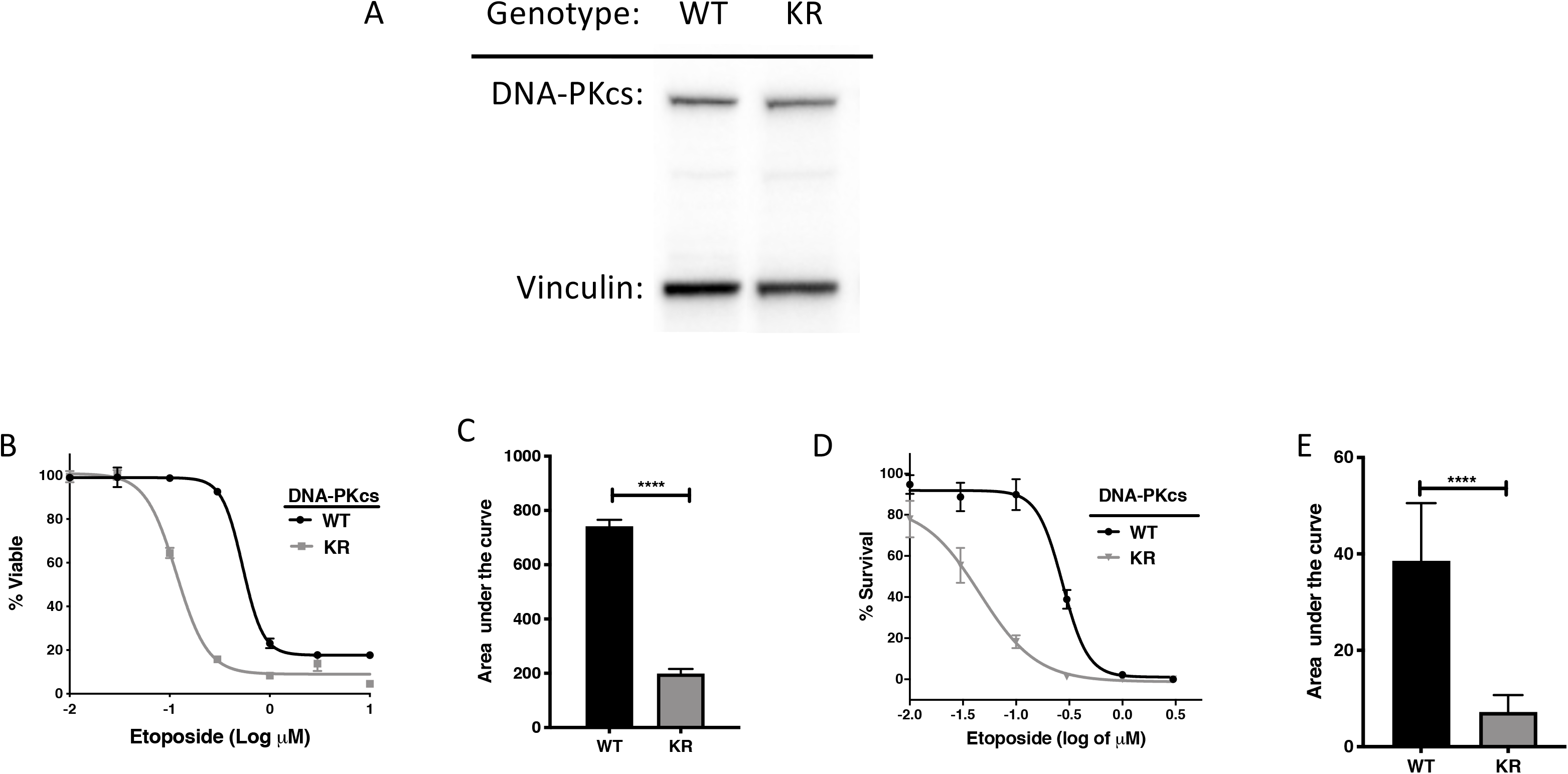
DNA-PK protects cells from etoposide toxicity via its kinase domain. (A) Evaluation of DNA-PK proteins in CHO variants by western blot. (B, C) Cells survival profile of CHO cells assessed using the CellTiter Glo (CTG) assay with increasing doses (0.003 – 1 μM) of etoposide at 72 h post-exposure with associated area under the curve analysis. (D, E) Clonogenic survival of DNA-PKcs WT, Null, or KR cells following exposure to etoposide (0.003 – 1 μM) with associated area under the curve analysis.

### Differential expression mediated by DNA-PKcs

A total of 12,759 genes were of sufficient quality to allow assessment of potential differential regulation in our samples (Table 1). In each of our comparisons, approximately 7,000 genes were differentially expressed, either up or down regulated. We assessed genotype specific differentially expressed genes (DEG) by comparing DMSO-treated WT versus KR gene expression, etoposide-induced alterations within each genotype compared to DMSO alone, and finally, differential expression due to drug treatment in WT compared to KR cells. Each red dot is one single gene that has a significant (p < 0.05) positive log-fold change and blue dot denotes gene with a significant negative log-fold change (Figure 2A – D). Each black dot represents a gene with a p-value that was not significantly altered (p ≥0.05).

**Table 1.** Summary of differentiallyexpressed genes in cells expressing wild-type DNA-PKcs compared to two mutant cell lines with and without exposure to etoposide.

| Comparisons |  | # Down | # Up | Sum changed | # Unchanged | Sum All |
| --- | --- | --- | --- | --- | --- | --- |
| **Genotype-specific alterations (DMSO only)** |  |  |  |  |  |  |
|  | WT vs KR | 3818 | 3798 | 7616 | 5143 | 12759 |
| **Etoposide-induced alterations by genotype** |  |  |  |  |  |  |
|  | WT | 3426 | 3082 | 6508 | 6251 | 12759 |
|  | KR | 3378 | 3762 | 7140 | 5619 | 12759 |
| **Genotype-specific alterations (Etoposide treatment)** |  |  |  |  |  |  |
|  | WT vs KR | 3285 | 3604 | 6889 | 5870 | 12759 |

**Figure 2:**
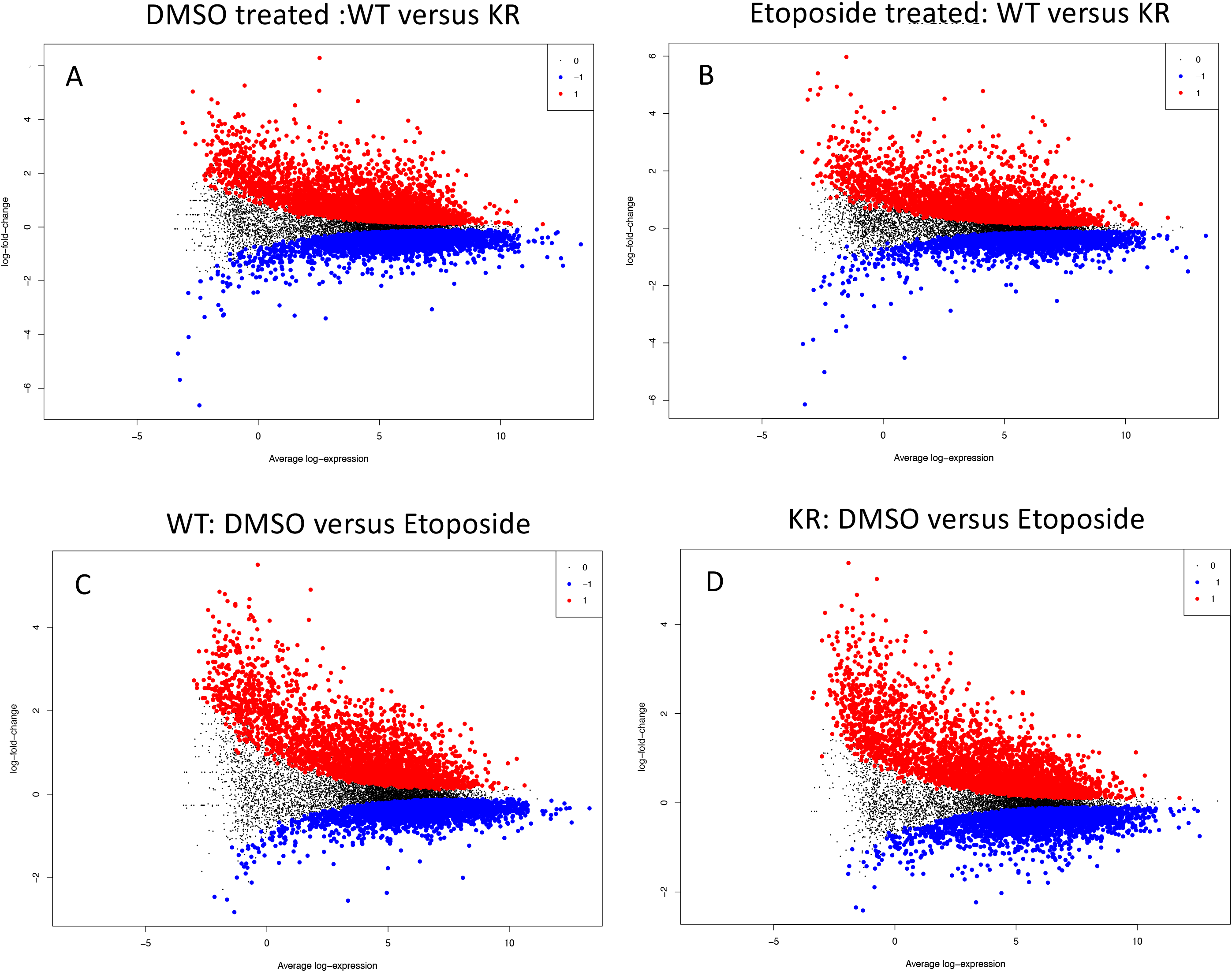
Differentially expressed genes in DNA-PK wild-type (WT) versus kinase-inactive (KR) cells. Each red dot is one single gene that has a significant (p < 0.05) positive log-fold change and blue dot denotes gene with a significant negative log-fold change. Each black dot defines a gene with a p-value that was not significantly altered (p > 0.05). A – C, represent the genes differentially expressed between DNA-PK variant genotypes. D – F, represent the ifferentially expressed genes following etoposide treatment in different genotypes. G – I, represent the differentially expressed genes within etoposide treated genotypes.

Upon assessing DEG in KR following etoposide to those in WT, we found most genes, 2511 of 4333, are induced independently of DNA-PKcs kinase activity, as these were upregulated in both KR and (Figure 3A), where 571 were discretly induced in WT and 1251 were observed only in KR cells. In comparing diminished expression, most, 2338 of 4466 genes were downregulated independently of DNA-PKcs kinase in both KR and WT in response to etoposide (Figure 3B). The number of privately downregulated DEGs are similar in WT (1088 DEGs) and KR (1040 DEGs) cells (Figure 3B).

**Figure 3:**
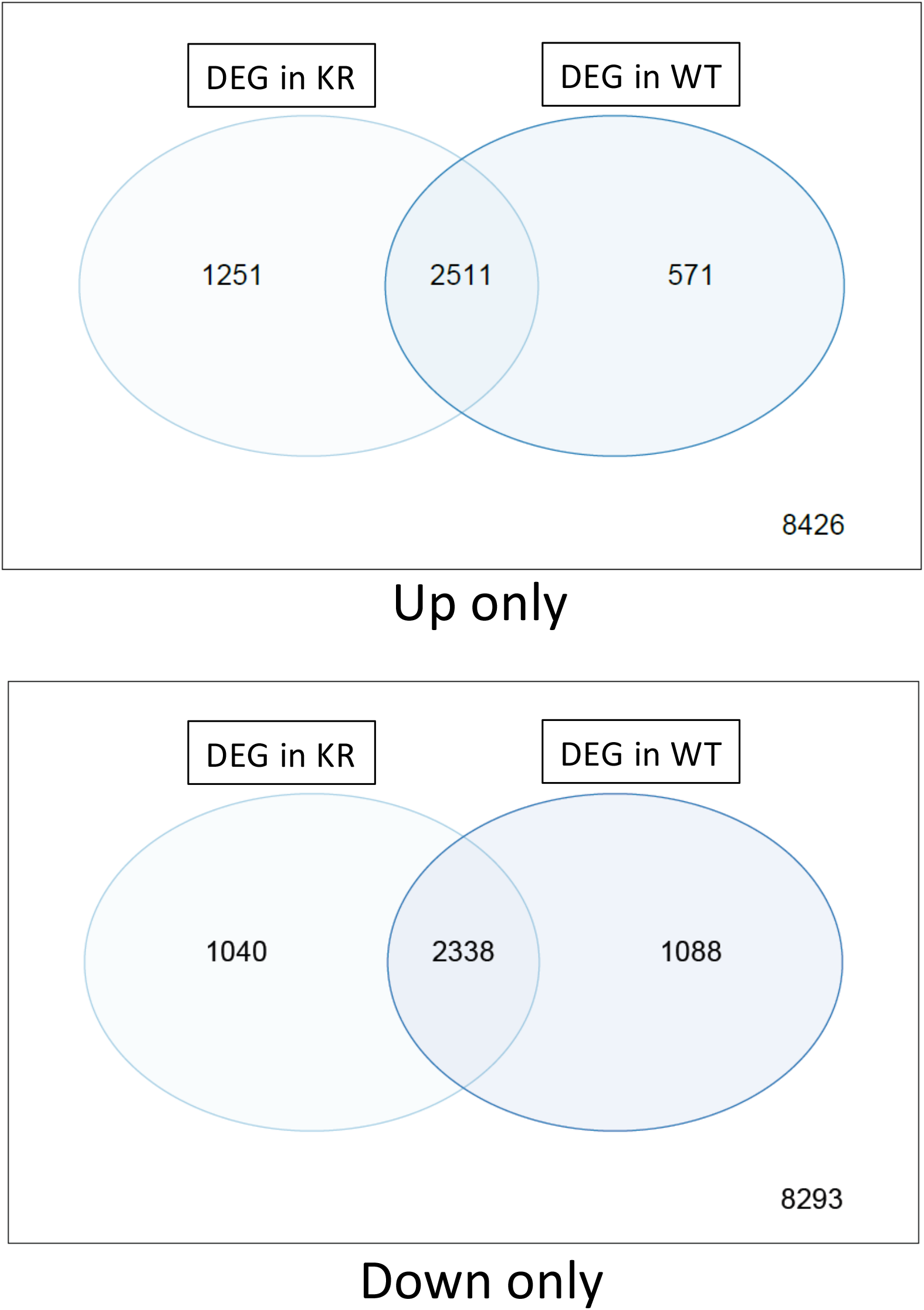
Comparison of shared versus discreet differentially regulated genes in DNA-PK wild-type (WT) versus kinase-inactive (KR) cells. A, Assessment of differentially regulated genes between genotype. B, Assessment of differentially regulated genes following etoposide treatment. Comparisons are made to DMSO-treated controls within genotype. C, Assessment of genotype-specific differentially expressed genes following etoposide treatment.

### DNA-PKcs alters the transcription of multiple genes involved in crucial cellular processes

DEG were analyzed with KEGG pathway data base and were involved in diverse cellular processes (supplemental figures S 1A – F). Upregulated genes are represented with a blue to red dots and downregulated genes are represented in blue to green dots. Corresponding pathways regulated by DEG are represented with a solid tangerine circle. The size of the tangerine circle is correlated with the number of genes. We analyzed these cellular pathways regulated by DEG to ascertain significantly altered biological processes within WT and KR cells in the context of normal homeostasis (DMSO) or DNA damage (etoposide).

### Genotype specific alterations in biological processes

We compared the biological pathways populated by the DEG in WT and KR cells without etoposide treatment. A total of 14 biological pathways were enriched in upregulated DEG (Figure 4A) and 7 pathways were associated with downregulation (Figure 4B) in KR cells compared to WT.

**Figure 4:**
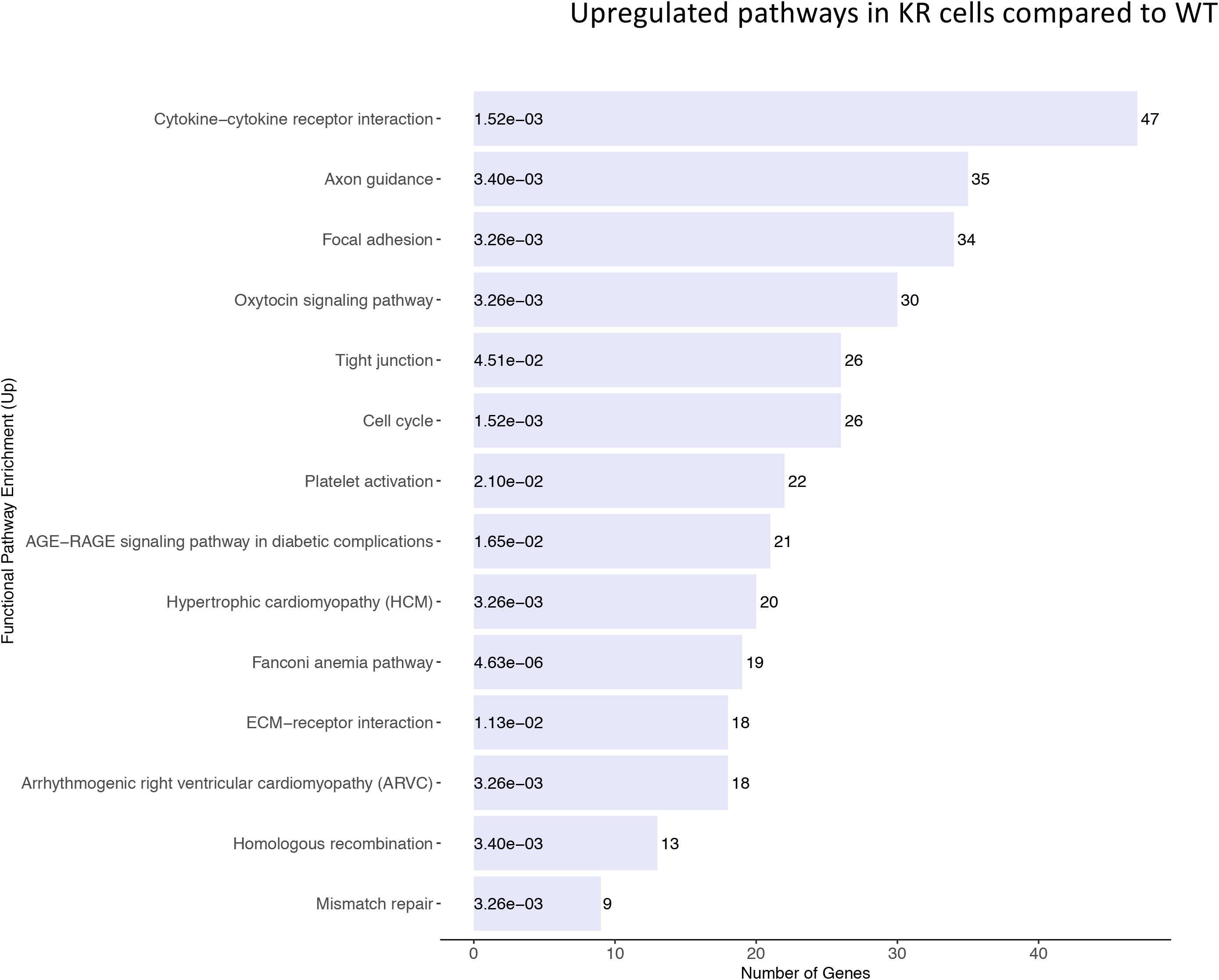
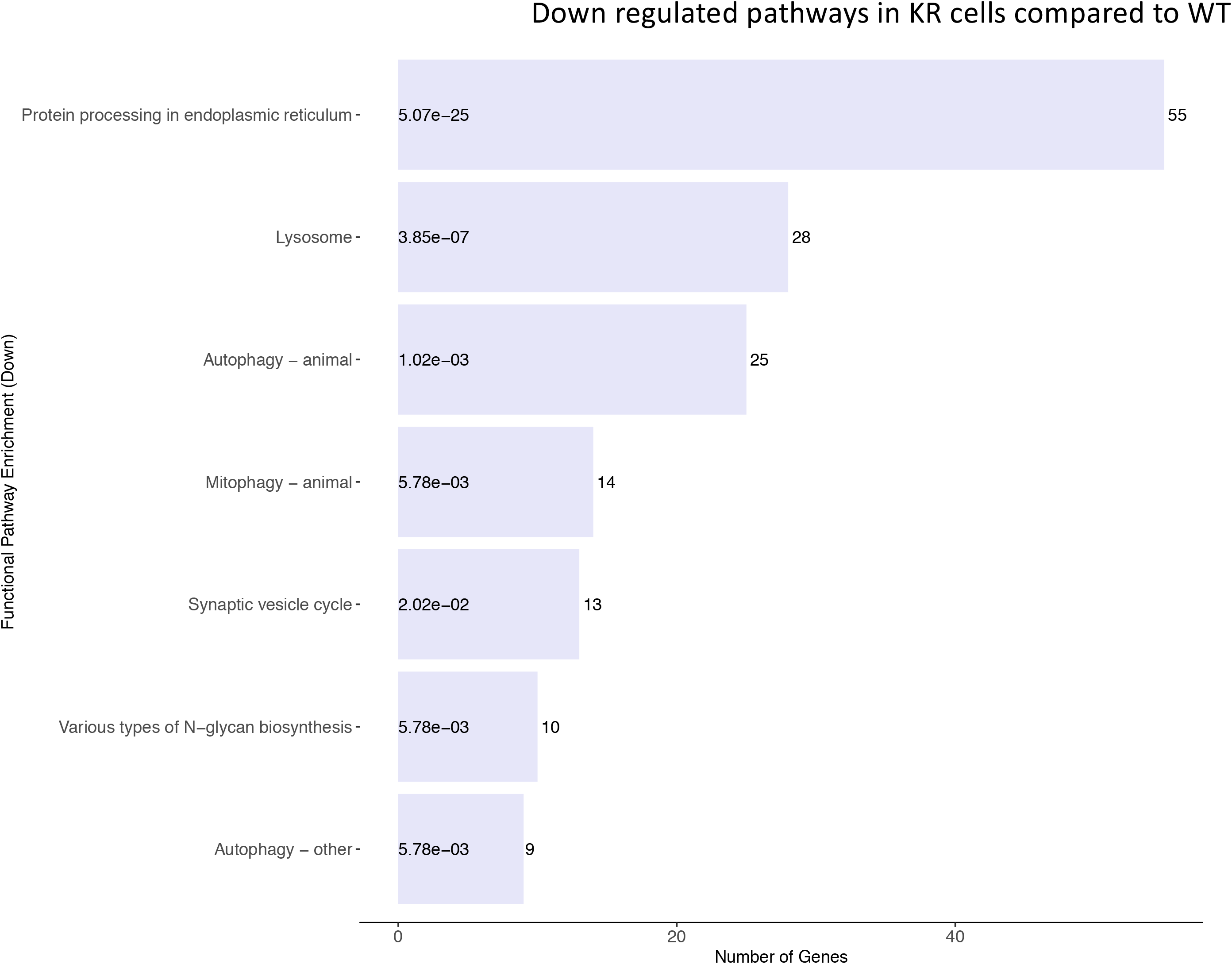
Genotype-specific biological pathways altered by differential gene regulation in DNA-PKcs wild-type (WT) versus kinase-inactive (KR) without drug exposure. A, Genotypic specific upregulated biological processes in KR. B, Genotypic specific downregulated biological processes in KR.

We observed several genes involved in DNA damage repair pathways, including homologous recombination, mismatch repair and Fanconi anemia pathways, were upregulated in KR cells (Figure 4A and S1A). Genes upregulated in KR cells compared to WT were predominantly involved in the cell cycle, cytokine signaling and associated anomalies, and cytoskeletal regulation (Figure 4A). Genes upregulated in KR compared to WT were involved ARVC, HCM, focal adhesion, axon guidance and cytokine signaling pathways which indicates DNA-PKcs kinase dependent regulation of these pathways. Either these are suppressed by DNA-PKcs endogenously or they are upregulated when DNA-PKcs cannot function as a kinase. In contrast, genes involved in protein processing and degradation, autophagy, protein and sugar metabolism were less abundant in KR cells compared to WT (Figure 4B and Figure S1B).

### Etoposide-induced differentially regulated biological pathways within WT and KR cells

We analyzed the role of DNA damage in differentially regulated biological pathways between cell lines. A total of 34 pathways were upregulated following etoposide treatment in WT cells (Figure 5A), whereas only two pathways were suppressed in WT cells (Figure 5B). In KR cells, 37 pathways were upregulated (Figure 5C) and 3 pathways were downregulated (Figure 5D) after etoposide treatment. Comparing KR to WT following drug treatment, five pathways were uniquely up regulated in WT, and 8 were observed only in KR following drug treatment. Most, 29 pathways of 37 were shared between genotypes (Figure 5A, 5C and 6A) indicating genes represented here are upregulated independent of DNA-PKcs kinase activity. Few pathways, 3, decreased in response to etoposide, and 2 pathways were observed in both cell lines, again, indicating they are regulated in a DNA-PKcs kinase independent manner (Figure 5B, 5D, and 6B).

**Figure 5:**
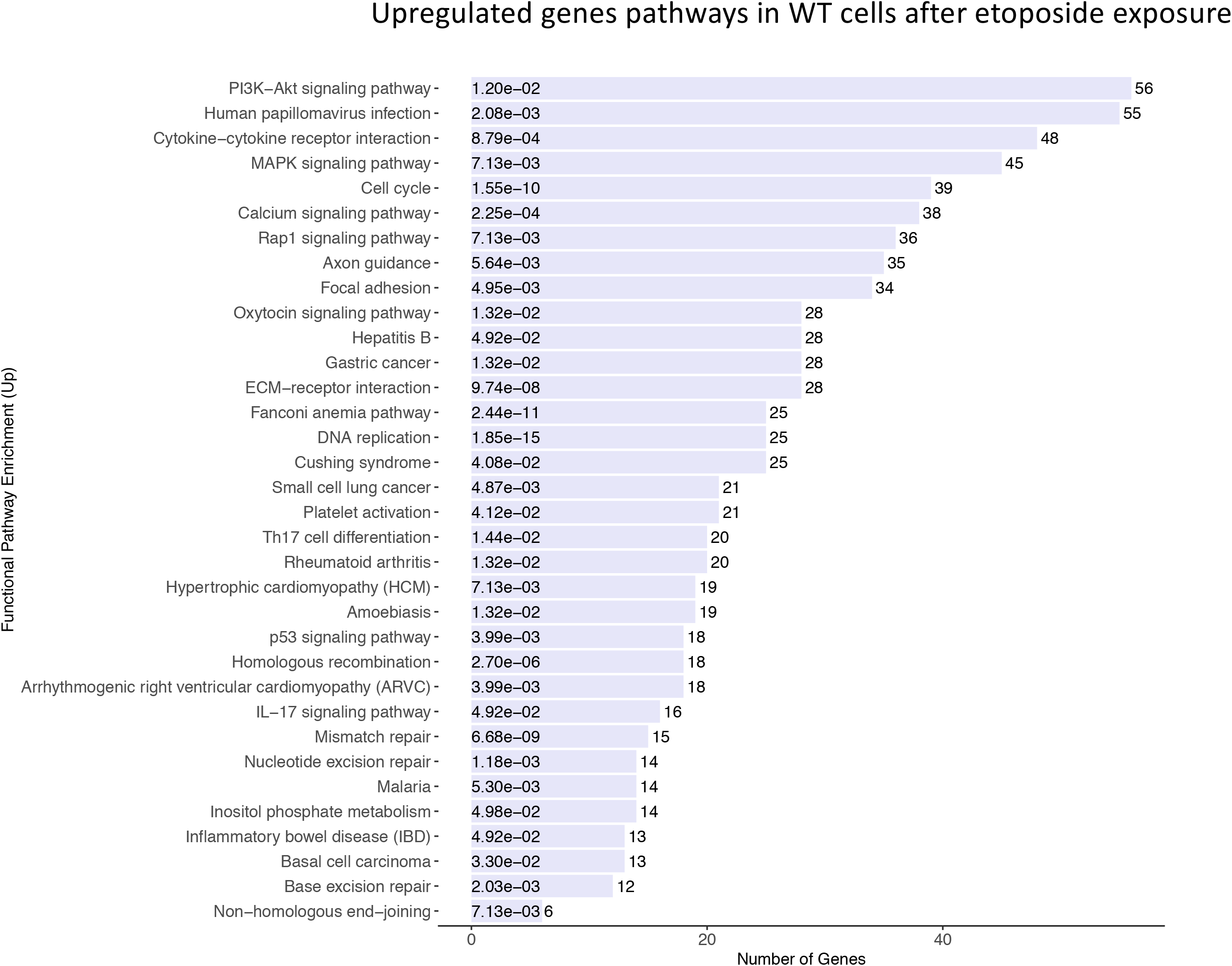
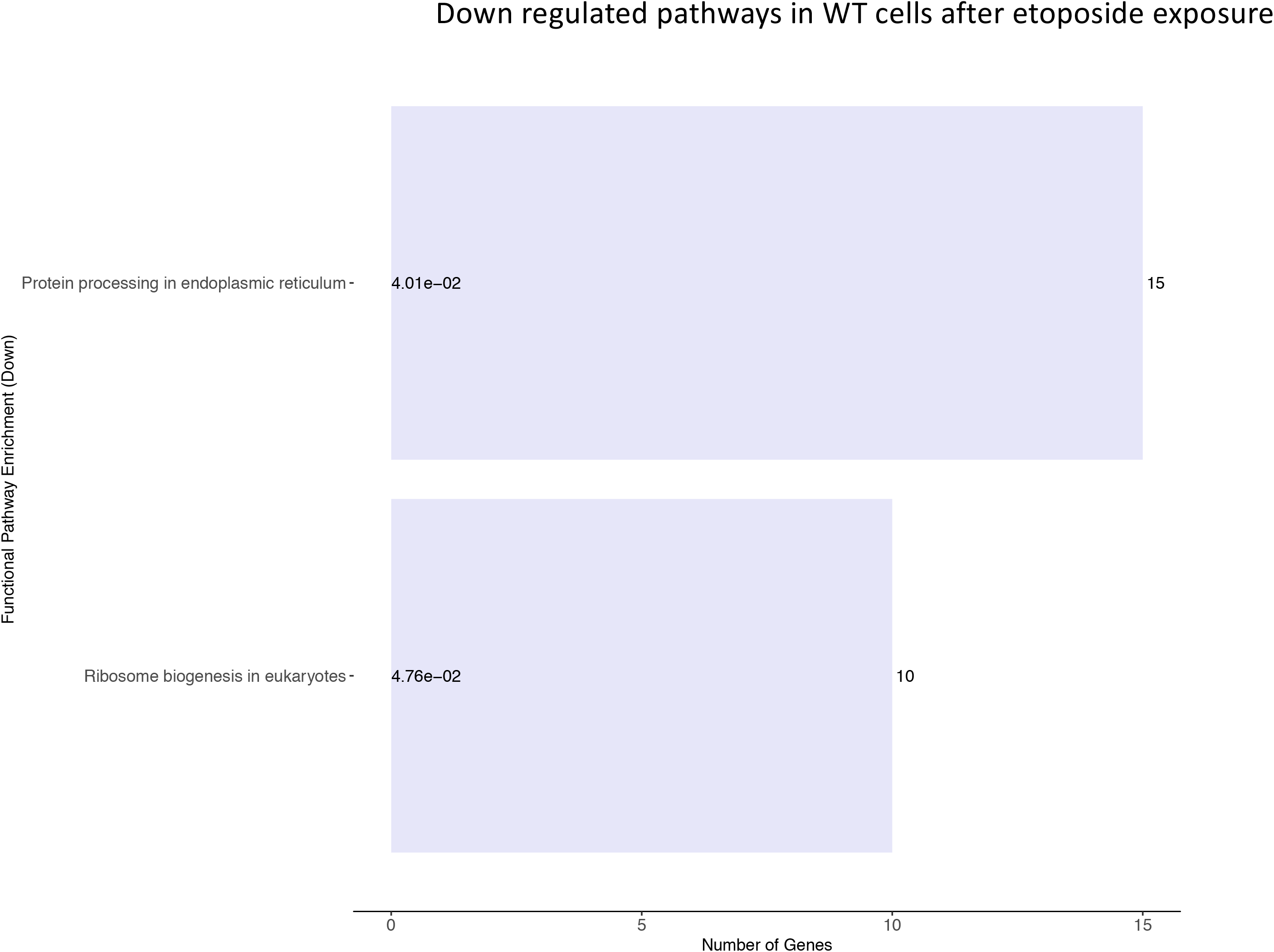
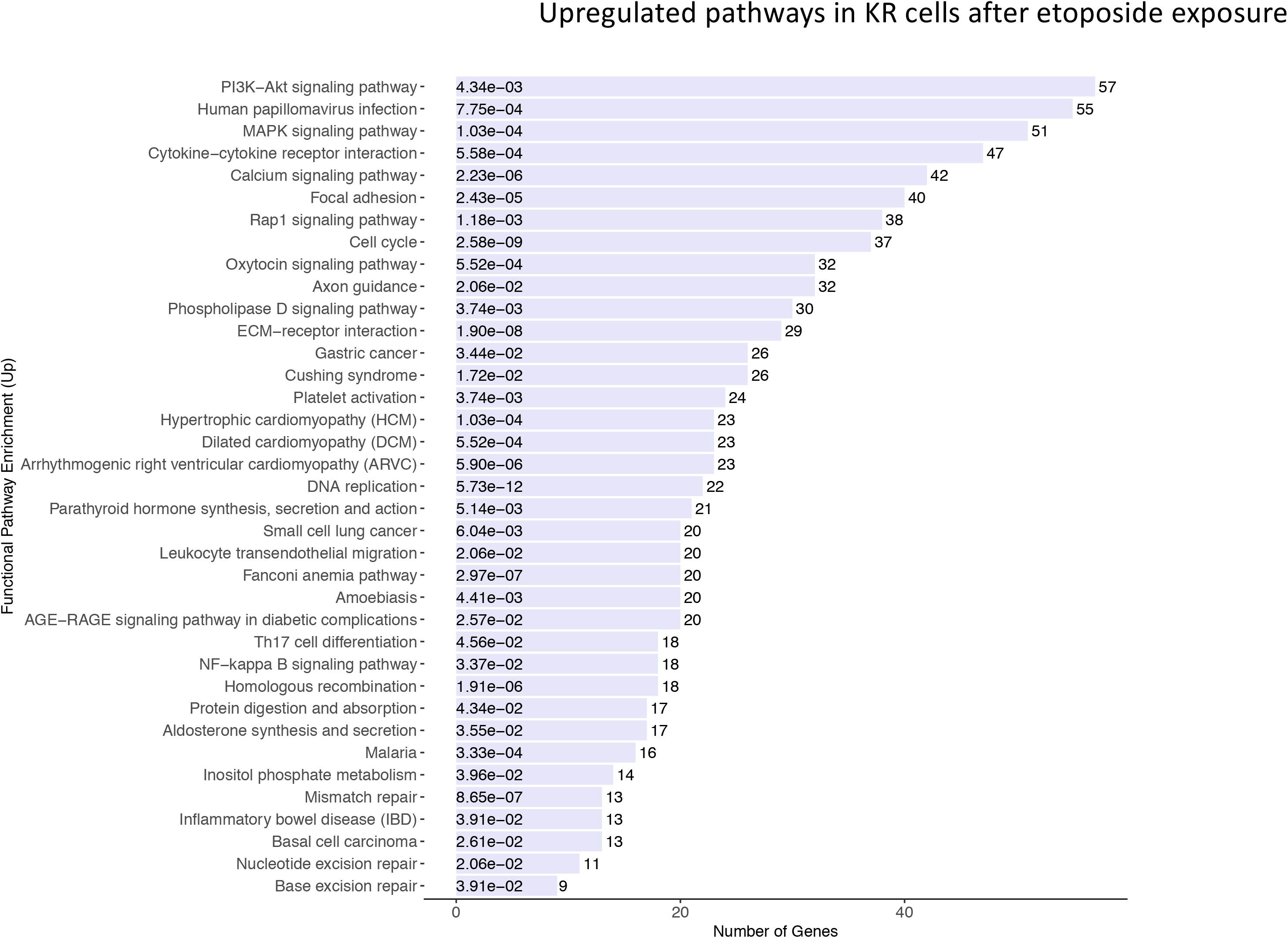
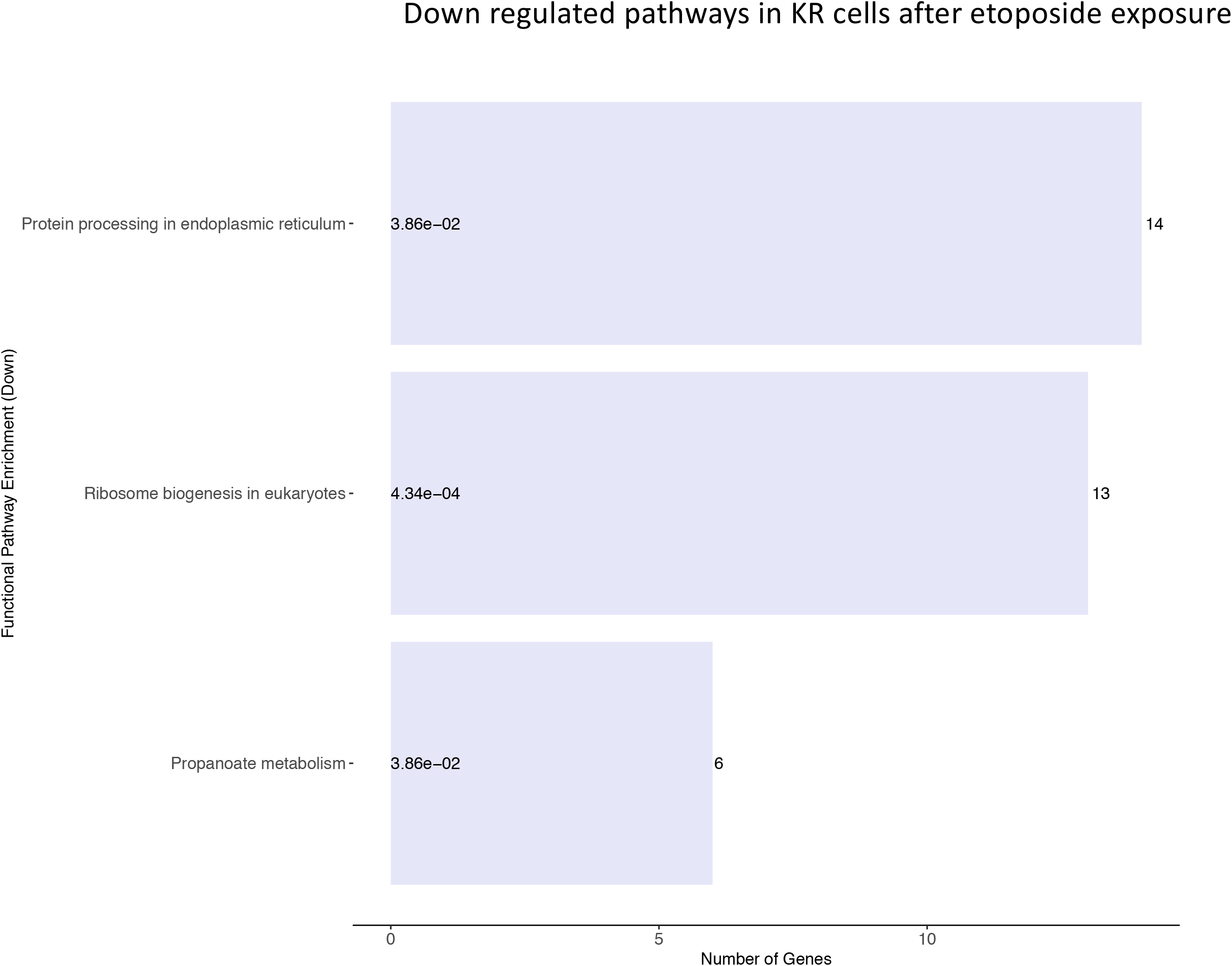
Etoposide-induced biological pathways altered by differential gene regulation in DNA-PKcs wild-type (WT) and kinase-inactive (KR) cells. A, Etoposide induced upregulated biological processes within WT cells. B, Etoposide induced downregulated biological processes in WT cells. C, Etoposide induced upregulated biological processes within KR cells. D, Etoposide induced downregulated biological processes in KR cells.

**Figure 6:**
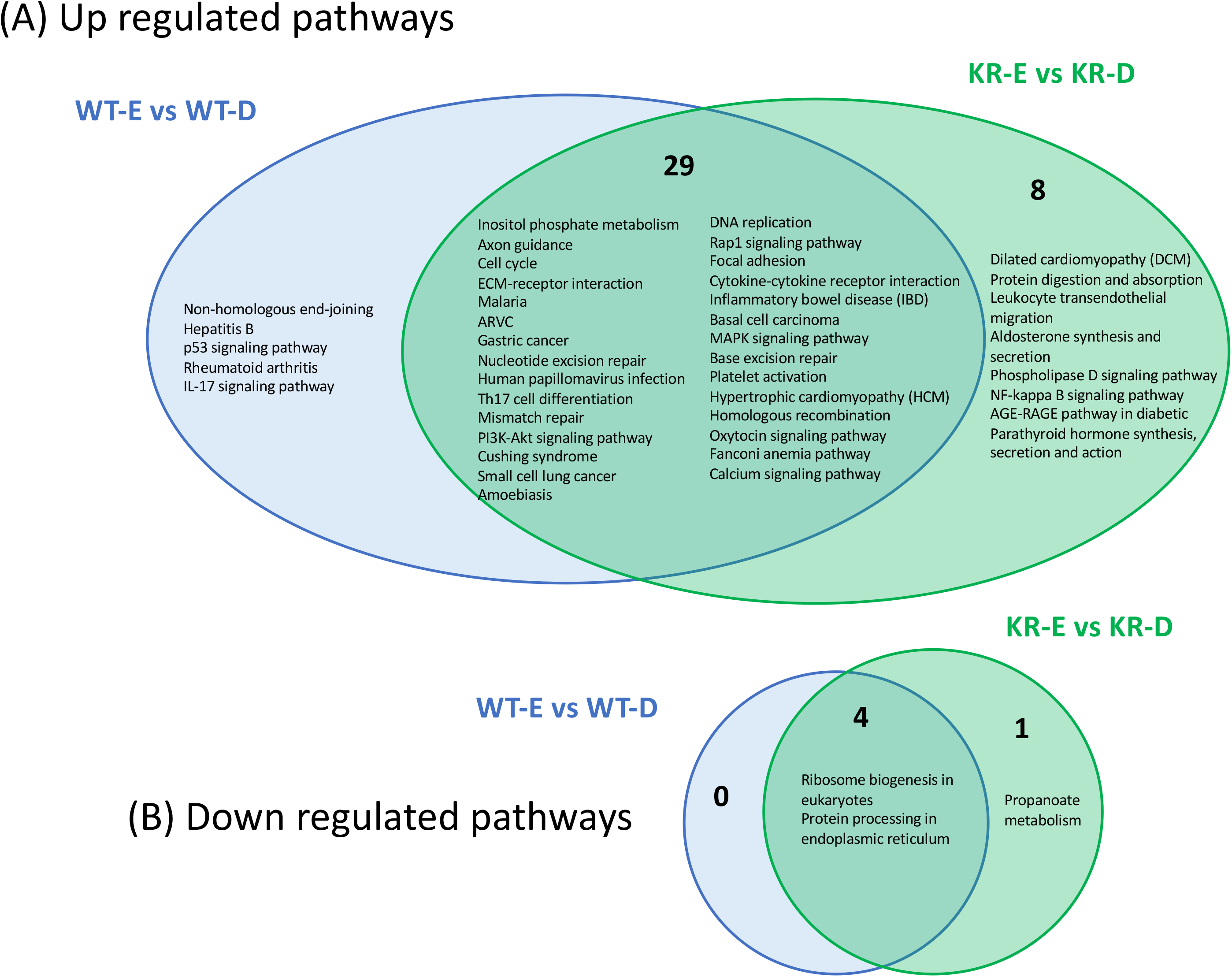
Comparison of shared versus divergent etoposide-induced biological pathways altered by differential gene regulation in DNA-PKcs wild-type (WT) compared to kinase-inactive (KR) cells. A, Etoposide induced upregulated biological processes within WT and KR cells. B, Etoposide induced downregulated biological processes in WT and KR cells

Multiple DNA damage repair processes were commonly upregulated in both cell types after etoposide treatment, whereas NHEJ was exclusively induced in WT cells (Figure 6A). Other commonly upregulated pathways include cell cycle, DNA replication, cytokine signaling, calcium signaling, and extracellular matrix (ECM)-receptor interactions are observed in both WT and KR cells. The two common pathways that were downregulated in both WT and KR cells were related to ribosome biogenesis and protein processing (Figure 6B).

### Differentially regulated biological processes between etoposide exposed WT and KR cells

We assessed the biological pathways impacted by differential gene regulation between each genotype after exposure to etoposide to discern how DNA-PKcs regulates expression changes following topoisomerase II inhibition. A total of 25 biological pathways were upregulated in etoposide treated KR cells compared to WT cells, whereas only three pathways were diminished (Figure 7A and B). When comparing the effects of etoposide on gene expression between genotypes, we found many genes upregulated in KR cells compared to WT are involved in multiple kinase-driven signaling pathways, including many involved in cancer biology, TNF signaling, cell-cell and cell-matrix interactions, and DNA damage response pathways. Interestingly genes involved in pathways related human papilloma virus infection were also upregulated in etoposide treated KR cells compared to etoposide exposed WT cells (Figure S1G). Downregulated genes in KR cells compared to WT after etoposide exposure are involved in mainly protein processing and carbon metabolism (Figure S1H).

**Figure 7:**
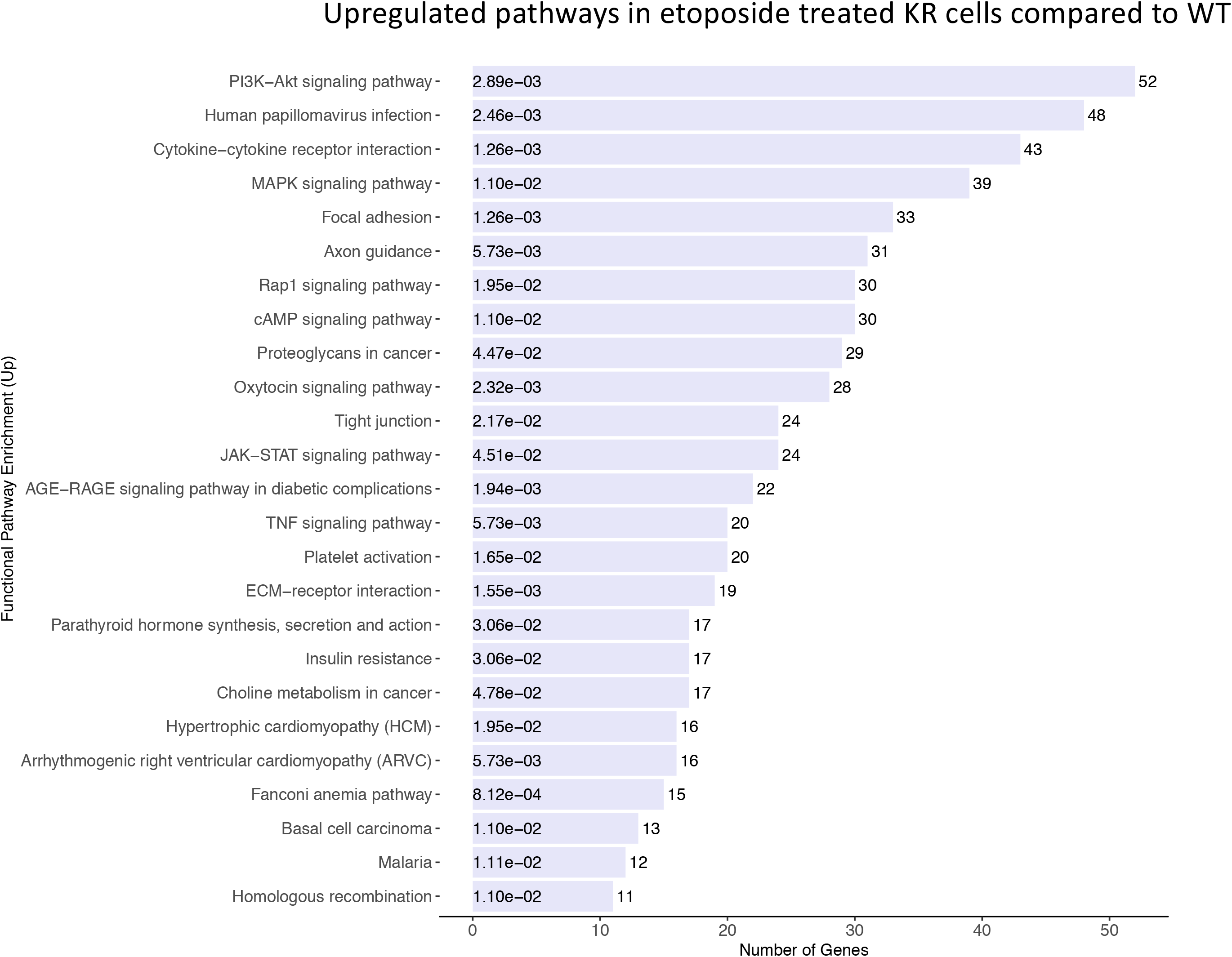
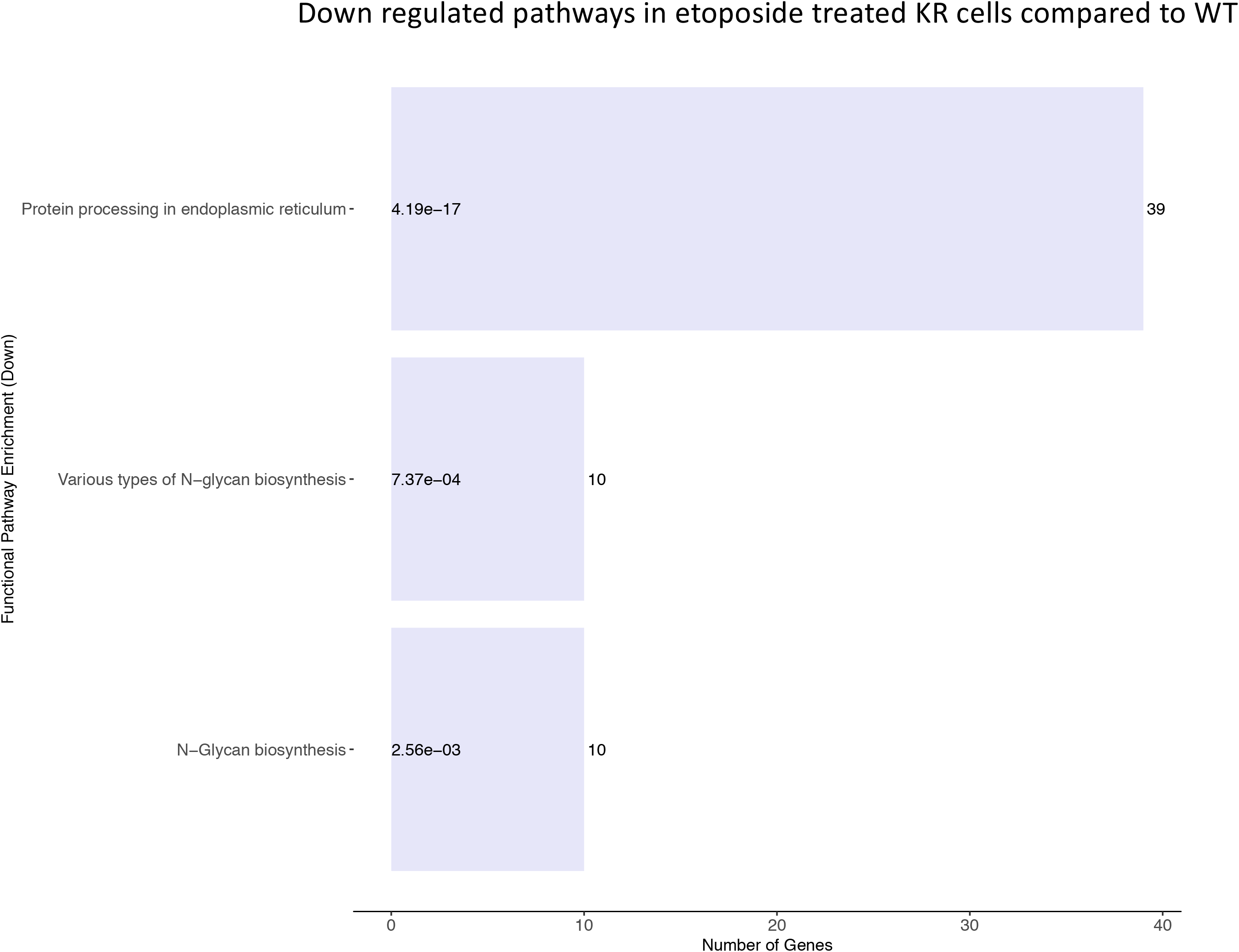
A comparative assessment of etoposide-induced biological pathways altered by differential gene regulation in DNA-PKcs wild-type (WT) and kinase-inactive (KR) cells. A, Upregulated biological processes in etoposide treated KR cells compared to WT Genotypic specific upregulated biological processes. B, Downregulated biological processes in etoposide treated KR cells compared to WT.

### Analysis of global transcriptional regulators

We evaluated our DEGs using Ingenuity Pathway Analysis (IPA, © 2000-2020 QIAGEN) to determine principal transcriptional regulators that drive gene expression changes observed in our samples (Table 2, Supplementary Tables 1-4). Master regulators controlling genes induced in KR without treatment include steroid-responsive proteins such as the estrogen receptor, growth hormone, IGFBP2, corticotropin releasing hormone receptor and others. We also note an enrichment in proteins involved in proliferation, including some listed above, as well as, NOTCH1, and oncogenes including Raf, KDR, MAP3K1FGR and RET. In addition, genes associated with the inflammatory response and angiogenesis, including VEGF and its receptors were observed.

**Table 2.**
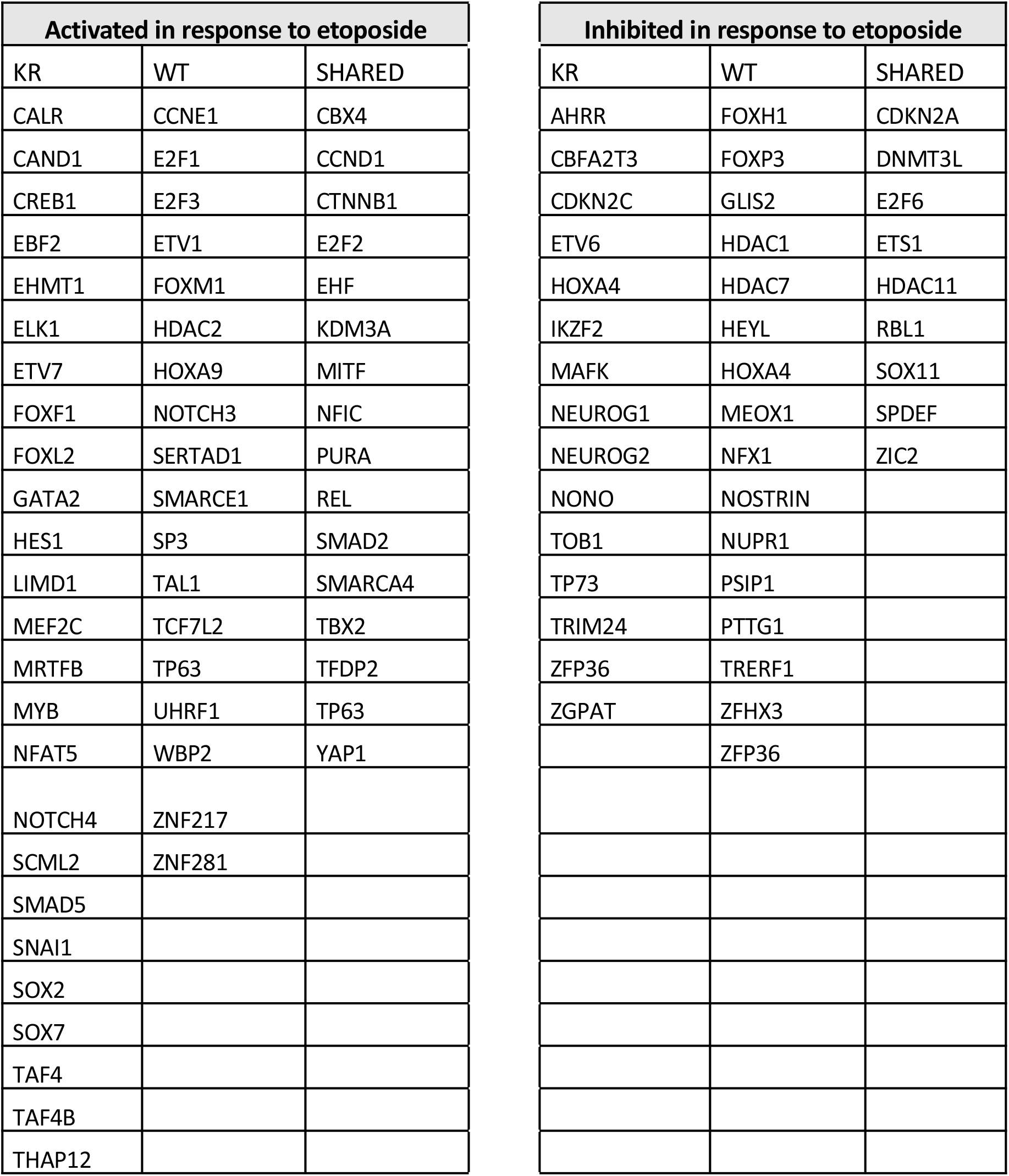
Summary of transcriptional regulators dictating differential gene expression in cells expressing wild-type (WT) or kinase inactive (KR) DNA-PKcs with and without exposure to etoposide.

Master regulators controlling down regulation include ZFHX3, ETV6, SPARC, SAFB and NEUROG2. Of note, many of these are potentially tumor suppressors and known modulators of gross or intracellular morphology. IPA analysis reveals that drugs inhibiting VEGF and various pro-growth pathways, including those induced by oncogenes such as Raf, KIT, ALK and ROS1 are diminished, consistent with what we observed based on upregulation. Overall, this indicates that the cellular milieu induced by expressing DNA-PKcs without functional kinase activity varies substantially compared to WT.

We also analyzed etoposide induced DEGs in all three DNA-PKcs variant cell types for detecting the master transcriptional regulators that modify DNA damage related gene expression. A total of 35 global regulator have been identified those were involved in the upregulation of DEGs in WT after etoposide exposure (Table 2). Some of these activators are HDAC2 (2.41E-15), TP63 (5.54E-15), CTNNB1 (2.23E-14), MITF (3.11E-14) andSMAD2 (6.76E-13). In comparison, 30 global regulators are involved in downregulation of the DEG in etoposide treated WT cells. Some of the transcriptional regulators are as follows: ETS1 (1.27E-23), MEOX1 (1.94E-15), CDKN2A (3.06E-15), HDAC7 (3.07E-15), ZFHX3 (3.08E-15) and SOX11 (4.21E-15). In KR cells, after etoposide exposure, a total of 46 global regulators, including ETV7 (3.02E-19), SMAD5 (3.47E-19), CTNNB1 (3.5E-16), MEF2C (1.26E-15) and SMAD2 (6.06E-15), were driving the upregulation of the genes. In contrast, 25 master regulators, including TOB1 (3.33E-19), ZGPAT (1.54E-17), MAFK (1.74E-17) and SOX11 (6.88E-17), were maintaining the downregulation of the genes in etoposide treated KR cells.

Ten (CBX4, SMARCA4, EHF, MITF, TFDP2, SMAD2, NFIC, KDM3A, PURA and E2F2) and five (RBL1, ETS1, CDKN2A, SOX11 and E2F6) master regulators were detected commonly in etoposide treated WT and KR cells, up and down-regulating genes, respectively, indicating these are regulated independently of DNA-PKcs. We analyzed the upstream master regulators between etoposide treated WT and KR cells. Comparing the upregulated genes in etoposide treated KR cells after with etoposide treated WT cells, we found14 global regulators, including MRTFB (1.82E-09) and TGIF1 (3.04E-09) were involved in upregulation of genes. In contrast, only 4 upstream global regulators, includes HOXA4 (1.43E-09) and NAB2 (1.14E-06), were driving the repression of down regulated genes in the KR cells after etoposide treatment.

A complete summary of IPA-designated master regulators is presented in Supplementary Tables 1 – 4; we opted to assess the transcription regulators modified in WT versus KR cells with and without drug treatment. A total of five (CCNE1, HDAC2, SMARCE1, UHRF1 and ZNF281) and four (FOXP3, HDAC1, HDAC7 and NUPR1) master regulators involving in, respectively, upregulation and down regulation of histone modifying genes were found in WT cells (Table 2). Two (SCML2 and SNAI1) or four master transcriptional regulators (AHRR, CBFA2T3, CDKN2C and TRIM24) were involved in altering transcription of histone modifying genes in KR cells. Three (CBX4, KDM3A and SMARCA4) master regulators inducing histone modifying genes were detected in both WT and KR cells while two regulators (DNMT3L and HDAC11) in both WT and KR cells are associated with down regulation of genes controlling histone modifications. We detected comparatively higher numbers of master regulators involve in development or cells differentiation related DEG in WT as well as KR cells. In WT, seven regulators (E2F1, ETV1, HDAC2, HOXA9, NOTCH3, TAL1 and ZNF281) induced genes involve in cells differential, whereas, eight (FOXP3, GLIS2, HDAC7, HOXA4, MEOX1, NUPR1, PSIP1 and ZFHX3) mediate the downregulation of DEG of similar function. Fourteen regulators (ETV7, FOXF1, FOXL2, GATA2, LIMD1, MEF2C, SCML2, SNAI1, SOX2, SOX7, MRTFB, MYB, NFAT5 and NOTCH4) were found in KR cells maintains the upregulation of genes involved in cell differentiation. While eight master regulators (AHRR, ETV6, HOXA4, IKZF2, MAFK, NEUROG1, NEUROG2 and TP73) were detected in downregulating similarly functioning in KR cells. Nine (CBX4, CTNNB1, EHF, MITF, NFIC, PURA, REL, TBX2 and YAP1) and five (DNMT3L, ETS1, HDAC11, SOX11 and ZFP36) master regulators were respectively mediate up and down regulation of cell differentiation genes commonly in both WT and KR cells. We found 11 master regulators (CCNE1, E2F1, EE2F3, FOXM1, HDAC2, NOTCH3, SERTAD1, SP3, TAL1, UHRF1 and ZNF217) in WT cells were maintain the upregulation of genes involved in cell cycle or proliferation, while only four master regulators were detected which to downregulates genes from the similar function. On the other hand, nine (CARL, EHMT1, ELK1, ETV7, FOXL2, LIMD1, MYB, NOTCH4 and THAP12) and seven (CBFA2T3, CDKN2C, ETV6, TOB1, TP73, TRIM24 and ZGPAT) master regulators were involved in respectively upregulating and down regulating cell cycle and proliferation mediated genes in KR cells. We also found nine master regulators (CCND1, E2F2, EHF, PURA, REL, SMAD2, SMARCA4, TFDP2 and YAP1) involved in upregulating cell cycle related genes commonly in both genotypes and eight (CDKN2A, E2F6, HDAC11, RBL1, SOX11, SPDEF and ZFP36) were involved in down regulating. Interestingly, eight master regulators (FOXP3, GLIS2, HDAC7, HOXA4, MEOX1, NUPR1, PSIP1 and ZFHX3) involved in down regulating WNT signaling pathways genes were found only in WT cells, whereas only one regulator, TCF7L2, were found in upregulation of genes in the same signaling pathway. In KR cells two regulators (EBF2 and TAF4) were involved in upregulated genes expression and only one master regulator, NEUROG1, is found controlling the down regulated genes in KR cells.

## Discussion

DNA-PKcs is a key regulator in two major DNA double strand break repair pathways, HR and NHEJ (2). In parallel to its role in DNA damage response, DNA-PKcs is also involved in phosphorylation of many other cellular proteins including zinc finger transcription factor (Snail1), heat-shock proteins (HSP90α), RNA polymerase II, and other transcription factors including SP1, Oct1, c-Myc, c-Jun (11, 33, 34). In this study, we use high throughput RNA sequence technology to analyze the total mRNA transcripts isolated from DNA-PKcs WT and kinase inactivated KR CHO cells after exposure to DNA damaging agent etoposide. We applied bioinformatic tools to analyze our RNA sequence data to analyze the differentially regulated biological pathways depend on DNA-PKcs kinase activity with or without DNA damage response.

We found a high number of DEG in all samples compare to each other. Previously, using subtractive hybridization technique of cDNA, a study demonstrated a limited role of DNA-PKcs in transcription after DNA damage (35). Similarly, low differential expression of other DNA repair genes was observed in DNA-PKcs deficient human glioblastoma cell line M059J cells (36). Microarray based gene expression analysis identified upregulation of 15 genes in DNA-PKcs depleted HeLa cells using targeted siRNA, most of the genes were associated with interferon-mediated signal transduction and cell proliferation (37). They also found eight downregulated genes, involved as nuclear factors of activated T lymphocytes. Recently a microarray based analysis of DEG showed that DNA-PKcs depletion induced upregulation of 418 and downregulated 1316 genes in human prostate cancer cell line, whereas Inhibition of DNA-PKcs by chemical inhibitor upregulated 1571 genes and downregulated 3270 genes (33). We conducted DEG analysis on DNA-PKcs wildtype and stable kinase inactivated CHO cell lines using sophisticated high through-put RNAseq technology. We detected more than 7000 DEG in KR cells compared to WT, while, etoposide treatment alone changed the expression of more than 6500 and 7100 genes in WT and KR respectively.

Differentially expressed genes were subjected to enrichment analysis using KEGG pathway database. We prepared barplots demonstrating the over-expressed or repressed biological processes related to the upregulated or downregulated genes in WT or KR cells with or without etoposide treatment. Upregulated genes in DMSO treated KR cells compared to WT were enriched three DNA damage repair pathways including HR. DNA-PKcs is a core component of NHEJ mediated DSB repair, and as KR cells express a catalytically inactive DNA-PKcs, these cells rely more heavily on HR to repair endogenously formed DSB. We also observed upregulation of several cell genes involved in cell cycle regulation and proliferation, in agreement with our previous work indicating DNA-PKcs regulates replication in a kinase-dependent fashion (6, 8). Genes from two DNA repair clusters, HR and Fanconi anemia predictably connected to the cell cycle via ATR (Figure S1).

Other genes upregulated in KR cells compared to WT were predominantly enriched cell cycle, cytokine signaling and associated anomalies, including arrhythmogenic right ventricular cardio myopathy (ARVC) and hypertrophic cardiomyopathy (HCM), and cytoskeletal regulation (Figure and S1). The upregulation of these genes in KR compared to WT without etoposide exposure indicates DNA-PKcs kinase dependent regulation of these pathways in unstressed conditions. Therefore, fully functional endogenously expressed DNA-PKcs would normally suppress these pathways via its kinase activity. Induction of apoptosis in cardiomyocytes is associated with ARVC. N226S mutation in human *DSG2* gene, which is important in structural integrity of cardiac discs by Ca^2+^ binding in intracellular cadherins molecule for cell-cell interaction, has been associated with ARVC (44). High inflammatory in filters and coagulative necrosis as well as apoptotic cardiomyocytes are observed in cardiac restricted over-expression of N271S mutation (mice homolog of human N22S mutation) containing transgenic mice (45). Mitochondrial dysfunction related oxidative stress in also associated with ARVC (reviewed in (46)). More in-depth investigation of the role of DNA-PKcs on ARVC is required to understand its progression.

Downregulated genes in KR compared to WT cells can be clustered into two larger groups of biological pathways, protein metabolism (includes protein processing, degradation, autophagy and lysosome) and sugar metabolism (amino and nucleotide sugar metabolism) (Figure 4B and Figure S1B). Interestingly, Goodwin et. al. previously reported that genes involve in HR, mismatch repair, cell cycle and DNA replication pathways were repressed in DNA-PKcs inhibited prostate cancer cell lines (35). The same study showed over-expression of various metabolic pathways in DNA-PKcs depleted cancer cells (35). Our observation did not follow the previously reported result where multiple genes associated with metabolic pathways were repressed because of the downregulation of related genes in absence functional DNA-PKcs in KR cells. These discrepancies may be attributable to our assessment in kinase inactivated CHO cells compared using DNA-PKcs inhibitor or siRNA mediated DNA-PKcs depletion.

We observed that 29 biological pathways were commonly upregulated in both WT and KR cell (Figure 6A). The genes involved in these pathways are upregulated in a DNA-PKcs kinase independent manner, which may indicate ATM-mediated regulation. ATM plays major roles in the DDR, namely assisting in pathway choice between NHEJ and HR. In addition, ATM and DNA-PKcs share many protein targets, often phosphorylating the same proteins. Previous studies consistently indicate altering DNA-PKcs reduces the level of ATM in only in absence of the DNA-PKcs protein, whereas the ATM levels remain same in WT in kinase inactivated KR cells (37–39). Similar ATM expression in WT and KR cells after etoposide exposure coupled with a lack of DNA-PKcs kinase activity may modify gene expression differences noted between WT and KR cells (40). Several genes involved in HR, hepatitis B signaling, rheumatoid arthritis, p53 signaling pathway and IL-17 signaling pathway were upregulated following etoposide treated KR cells alone, indicating their transcription may regulated by DNA-PKcs kinase and other DNA damage response proteins, including ATM. HR differences are noted in the WT versus KR cells following exposure to other genotoxic agents, as we demonstrated the KR cells promote higher levels of recombination compared to WT cells or cells lacking DNA-PKcs (8). Future studies will be aimed at determining whether ATM may also be promoting alterations in gene expression in the context of DNA-PK inhibition.

To assess the influence DNA-PKcs exerts the cellular transcriptome following DNA damage, we treated cells with etoposide (topoisomerase II inhibitor, a widely used DNA damaging agent) and analyzed the transcriptomes. Genes involved in NHEJ are discretely upregulated in WT cells in presence of fully functional DNA-PKcs. Etoposide exposure modified the global transcription of several genes in all three genotypes. In both genotypes, transcripts involved in ECM – receptor interaction, HR, Fanconi Anemia and cell cycle regulation are upregulated following etoposide exposure, and genes involved to ribosome biogenesis are downregulated independently of DNA-PKcs. Previous report suggested several DNA repair proteins have defined role in ribosome biogenesis and vice versa (41). In WT cells, genes involved in calcium signaling, mismatch repair and DNA replication were upregulated in WT and KR cells in response to etoposide only in presence of DNA-PKcs protein. Upregulation of these genes in both WT and KR cells indicate their transcriptional regulation might be independent to DNA-PKcs kinase activity in response to etoposide exposure. Genes associated with various DNA repair pathways including HR, mismatch repair, nucleotide excision repair (NER), base excision repair (BER) and FA along with DNA replication were upregulated in WT and KR cells after treatment with etoposide. Genes involved to HIF-1 signaling pathways were repressed in WT after etoposide exposure which support the fact that DNA-PKcs positively regulates transcription factor HIF1 (14). In response to etoposide, genes modulating NFκB signaling were upregulated KR cells; previous studies suggested ATM may contribute to secretion of signaling molecules like NFκB (42) and inflammatory cytokines like interleukin-6 and interleukin-8 following DNA damage (43). Consistent with our data, adipocyte degeneration due to over activation of transcription of chronic pro-inflammatory factors IL-6, TNF and KC (murine homolog to IL-8) were correlated with persistent DNA damage in mice (44). DNA damage induces secretion of NFκB, which can activate interferon (IFN) signaling and induce the expression of IFN-α and IFN-λ genes (45). Etoposide induced genes associated with cardiovascular disease in KR cells. Induction of apoptosis in cardiomyocytes is associated with ARVC. N226S mutation in human *DSG2* gene, which is important in structural integrity of cardiac discs by Ca^2+^ binding in intracellular cadherins molecule for cell-cell interaction, has been associated with ARVC (46). High inflammatory infilters and coagulative necrosis as well as apoptotic cardiomyocytes are observed in cardiac restricted over-expression of N271S mutation (mice homolog of human N22S mutation) containing transgenic mice (47). Mitochondrial dysfunction related oxidative stress in also associated with ARVC (reviewed in(48)). Many genes associated with amino acid biosynthesis and protein metabolism were repressed in KR cells after etoposide exposure suggesting their expression controlled either by DNA-PKcs kinase activity.

We analyzed the upstream regulators those are involved in globally regulating the transcription of the genes with or without etoposide treatment. Among genotypes the changes in gene expression were dominantly regulated by Akt signaling pathway. A large percentage of the master regulators are fall within the category of transcription regulators, chemical drugs, enzymes and other biological drugs. Among others, upstream regulators included cytokines, growth factors, (e.g. VEGF), G-protein coupled receptors, various kinase and transporters. Some peptidase and phosphatases are also found as master regulator in etoposide treated WT and KR cells. Notably, alterations in histone modifiers, cancer signaling/proliferation/apoptosis, Wnt/β-catenin signaling, and regulators of development and differentiation were altered in our samples (Table 2). Wnt signaling was more frequently observed in WT cells, indicating DNA-PKcs may regulate this pathway, as we have previously reported (49).The abundance of regulators associated with development, differentiation, and steroid signaling are likely due to the origin of the cells as oocytes. However, we have previously indicated a role for DNA-PK in differentiation and stemness (49), so these observations warrant additional investigation. Master regulators involved in histone modification were found in response to etoposide mediated DNA damage. DNA-PKcs phosphorylates histone variant H2AX and promotes chromatin decondensation immediately after DSB to facilitate recruitment of DDR proteins (12). The presence of histone modifier in both WT and KR cells indicates DNA-PKcs has a role in dictating chromatin modification in DNA damage response in both kinase dependent and independent manners. Though common regulators were observed, so were discreet regulators altered in either the WT or KR cells, suggesting DNA-PKcs, in a kinase-dependent fashion, modifies a subset of transcriptional regulators following DNA damage. The KR cells produce a more robust inflammatory response to DNA damage, which may induce further secondary tumor formation if a patient is treated with a DNA-PK inhibitor in combination with chemo- or radiotherapy. This would a significant modification of the tumor microenvironment and may lead to secondary tumor formation. Further investigations into the clinical ramifications of this data will clarify if DNA-PKcs mitigates the inflammatory response following DNA damage.

Overall this global transcriptome study will help us to identify role of DNA-PKcs in transcriptional regulation with or without DNA damage response. We anticipate our results will provide additional knowledge about how DNA-PKcs might modify the transcriptional response to DNA damage, leading to alterations of many essential cellular pathways, which ultimately dictate genomic stability. Multiple DNA-PKcs inhibitors are in clinical trials as potential therapeutic interventions in multiple cancer subtypes coupled with traditional chemotherapy and/or radiotherapy; however, we lack a full appreciation for how inhibition of DNA-PKcs kinase will alter the cellular transcriptional response to DNA damage. Our results indicate impairing DNA-PKcs kinase activity may profoundly impact the transcriptional response to DNA damage, modulating the proclivity for secondary tumor formation both via impairing traditional DNA damage signaling and repair, as well as impairing normal transcriptional changes aimed at maintaining genomic stability.

## Supporting information

https://ashleylab.nmsu.edu/supplementary-data/

---

**Supplementary Table 1. Genotypic master regulators controlling the gene expression differences observed in cells producing wild-type (WT) or kinase inactive (KR) DNA-PKcs.**

**Supplementary Table 2. All master regulators controlling the gene expression differences observed in cells producing wild-type (WT) DNA-PKcs following etoposide treatment.**

**Supplementary Table 3. Master regulators controlling the gene expression differences observed in cells producing kinase-inactive (KR) DNA-PKcs following etoposide treatment.**

**Supplementary Table 4. Genotypic master regulators controlling the gene expression differences observed in cells producing wild-type (WT) or kinase inactive (KR) DNA-PKcs after etoposide exposure.**

**Supplemental Figure 1 (A – Q)**: Bar-plots represented differentially regulated biological pathways in DNA-PKcs variants genotypes followed by etoposide exposure.

**upplemental Figure 2**: Nearest neighbor plots represents major differentially regulated biological pathways in DNA-PKcs variants genotypes followed by etoposide exposure.

**Supplemental Figure 3**: Differentially regulated genes with major biological pathways in DNA-PKcs variants genotypes followed by etoposide exposure.

## Acknowledgements

The authors would like to acknowledge Dr. Kathy Meek from Michigan State University for the DNA-PK cell lines. The authors would also like to thank Dr. Ryan Ashley for valuable input on this project. Research reported in this publication was supported by Cowboys for Cancer Research Pilot Grant, NM-INBRE (Institutional Development Award (IDeA) of the National Institutes of Health (NIH) & National Institute of General Medical Sciences (NIGMS) Grant # P20GM103451), and the Partnership for the Advancement of Cancer Research: NMSU/FHCRC, supported in part by NCI grants U54 CA132383 (NMSU) and U54 CA132381 (FHCRC).

## References

1. Lindahl T & Barnes, DE (2000) Repair of endogenous DNA damage. Cold Spring Harb Symp Quant Biol 65:127–133.

2. Blackford AN & Jackson, SP (2017) ATM, ATR, and DNA-PK: The Trinity at the Heart of the DNA Damage Response. Mol Cell 66(6):801–817.

3. Yoo S & Dynan, WS (1999) Geometry of a complex formed by double strand break repair proteins at a single DNA end: recruitment of DNA-PKcs induces inward translocation of Ku protein. Nucleic Acids Res 27(24):4679–4686.

4. Neal JA & Meek, K (2011) Choosing the right path: does DNA-PK help make the decision? Mutat Res 711(1-2):73–86.

5. Xing M & Oksenych, V (2019) Genetic interaction between DNA repair factors PAXX, XLF, XRCC4 and DNA-PKcs in human cells. FEBS Open Bio 9(7):1315–1326.

6. Liu S, et al. (2012) Distinct roles for DNA-PK, ATM and ATR in RPA phosphorylation and checkpoint activation in response to replication stress. Nucleic Acids Res 40(21):10780–10794.

7. Karmakar P, et al. (2002) Werner protein is a target of DNA-dependent protein kinase in vivo and in vitro, and its catalytic activities are regulated by phosphorylation. J Biol Chem 277(21):18291–18302.

8. Ashley AK, et al. (2014) DNA-PK phosphorylation of RPA32 Ser4/Ser8 regulates replication stress checkpoint activation, fork restart, homologous recombination and mitotic catastrophe. DNA Repair (Amst) 21:131–139.

9. Shrivastav M, De Haro LP, & Nickoloff JA (2008) Regulation of DNA double-strand break repair pathway choice. Cell Res 18(1):134–147.

10. Ashley AK & Kemp, CJ (2017) DNA-PK, ATM, and ATR: PIKKing on p53. Cell Cycle:1–5.

11. Collis SJ, DeWeese TL, Jeggo PA, & Parker AR (2005) The life and death of DNA-PK. Oncogene 24(6):949–961.

12. Lu H, Saha J, Beckmann PJ, Hendrickson EA, & Davis AJ (2019) DNA-PKcs promotes chromatin decondensation to facilitate initiation of the DNA damage response. Nucleic Acids Res.

13. Goodwin JF & Knudsen, KE (2014) Beyond DNA repair: DNA-PK function in cancer. Cancer discovery 4(10):1126–1139.

14. Bouquet F, et al. (2011) A DNA-dependent stress response involving DNA-PK occurs in hypoxic cells and contributes to cellular adaptation to hypoxia. J Cell Sci 124(Pt 11):1943–1951.

15. Wong RH, et al. (2009) A role of DNA-PK for the metabolic gene regulation in response to insulin. Cell 136(6):1056–1072.

16. Park SJ, et al. (2017) DNA-PK Promotes the Mitochondrial, Metabolic, and Physical Decline that Occurs During Aging. Cell Metab 26(2):447.

17. Jackson SP, MacDonald JJ, Lees-Miller S, & Tjian R (1990) GC box binding induces phosphorylation of Sp1 by a DNA-dependent protein kinase. Cell 63(1):155–165.

18. Maldonado E, et al. (1996) A human RNA polymerase II complex associated with SRB and DNA-repair proteins. Nature 381(6577):86–89.

19. Dvir A, Peterson SR, Knuth MW, Lu H, & Dynan WS (1992) Ku autoantigen is the regulatory component of a template-associated protein kinase that phosphorylates RNA polymerase II. Proc Natl Acad Sci U S A 89(24):11920–11924.

20. Lees-Miller SP (1996) The DNA-dependent protein kinase, DNA-PK: 10 years and no ends in sight. Biochem Cell Biol 74(4):503–512.

21. Ju BG & Rosenfeld, MG (2006) A breaking strategy for topoisomerase IIbeta/PARP-1-dependent regulated transcription. Cell Cycle 5(22):2557–2560.

22. Ju BG, et al. (2006) A topoisomerase IIbeta-mediated dsDNA break required for regulated transcription. Science 312(5781):1798–1802.

23. Neal JA, et al. (2014) Unraveling the complexities of DNA-dependent protein kinase autophosphorylation. Mol Cell Biol 34(12):2162–2175.

24. Joshi RR, Ali SI, & Ashley AK (2019) DNA Ligase IV Prevents Replication Fork Stalling and Promotes Cellular Proliferation in Triple Negative Breast Cancer. J Nucleic Acids 2019:9170341.

25. Bolger AM, Lohse M, & Usadel B (2014) Trimmomatic: a flexible trimmer for Illumina sequence data. Bioinformatics 30(15):2114–2120.

26. Kim D, Langmead B, & Salzberg SL (2015) HISAT: a fast spliced aligner with low memory requirements. Nat Methods 12(4):357–360.

27. Pertea M, et al. (2015) StringTie enables improved reconstruction of a transcriptome from RNA-seq reads. Nature biotechnology 33(3):290–295.

28. Liao Y, Smyth GK, & Shi W (2014) featureCounts: an efficient general purpose program for assigning sequence reads to genomic features. Bioinformatics 30(7):923–930.

29. Robinson MD, McCarthy DJ, & Smyth GK (2010) edgeR: a Bioconductor package for differential expression analysis of digital gene expression data. Bioinformatics 26(1):139–140.

30. Ritchie ME, et al. (2015) limma powers differential expression analyses for RNA-sequencing and microarray studies. Nucleic Acids Res 43(7):e47.

31. Yu G, Wang LG, Han Y, & He QY (2012) clusterProfiler: an R package for comparing biological themes among gene clusters. OMICS 16(5):284–287.

32. Kramer A, Green J, Pollard J, Jr., & Tugendreich S (2014) Causal analysis approaches in Ingenuity Pathway Analysis. Bioinformatics 30(4):523–530.

33. Goodwin JF, et al. (2015) DNA-PKcs-Mediated Transcriptional Regulation Drives Prostate Cancer Progression and Metastasis. Cancer Cell 28(1):97–113.

34. Park SJ, et al. (2017) DNA-PK Promotes the Mitochondrial, Metabolic, and Physical Decline that Occurs During Aging. Cell Metab 25(5):1135–1146 e1137.

35. Bryntesson F, Regan JC, Jeggo PA, Taccioli GE, & Hubank M (2001) Analysis of gene transcription in cells lacking DNA-PK activity. Radiat Res 156(2):167–176.

36. Galloway AM & Allalunis-Turner J (2000) cDNA expression array analysis of DNA repair genes in human glioma cells that lack or express DNA-PK. Radiat Res 154(6):609–615.

37. An J, Xu QZ, Sui JL, Bai B, & Zhou PK (2005) Silencing of DNA-PKcs alters the transcriptional profile of certain signal transduction genes related to proliferation and differentiation in HeLa cells. Int J Mol Med 16(3):455–462.

38. Ng WL, Yan D, Zhang X, Mo YY, & Wang Y (2010) Over-expression of miR-100 is responsible for the low-expression of ATM in the human glioma cell line: M059J. DNA Repair (Amst) 9(11):1170–1175.

39. Peng Y, et al. (2005) Deficiency in the catalytic subunit of DNA-dependent protein kinase causes down-regulation of ATM. Cancer Res 65(5):1670–1677.

40. Shrivastav M, et al. (2009) DNA-PKcs and ATM co-regulate DNA double-strand break repair. DNA Repair (Amst) 8(8):920–929.

41. Ogawa LM & Baserga, SJ (2017) Crosstalk between the nucleolus and the DNA damage response. Mol Biosyst 13(3):443–455.

42. Wu ZH, Shi Y, Tibbetts RS, & Miyamoto S (2006) Molecular linkage between the kinase ATM and NF-kappaB signaling in response to genotoxic stimuli. Science 311(5764):1141–1146.

43. Rodier F, et al. (2009) Persistent DNA damage signalling triggers senescence-associated inflammatory cytokine secretion. Nat Cell Biol 11(8):973–979.

44. Karakasilioti I, et al. (2013) DNA damage triggers a chronic autoinflammatory response, leading to fat depletion in NER progeria. Cell Metab 18(3):403–415.

45. Brzostek-Racine S, Gordon C, Van Scoy S, & Reich NC (2011) The DNA damage response induces IFN. J Immunol 187(10):5336–5345.

46. Pilichou K, et al. (2006) Mutations in desmoglein-2 gene are associated with arrhythmogenic right ventricular cardiomyopathy. Circulation 113(9):1171–1179.

47. Pilichou K, et al. (2009) Myocyte necrosis underlies progressive myocardial dystrophy in mouse dsg2-related arrhythmogenic right ventricular cardiomyopathy. J Exp Med 206(8):1787–1802.

48. van Opbergen CJM, den Braven L, Delmar M, & van Veen TAB (2019) Mitochondrial Dysfunction as Substrate for Arrhythmogenic Cardiomyopathy: A Search for New Disease Mechanisms. Front Physiol 10:1496.

49. Gurley KE, Ashley AK, Moser RD, & Kemp CJ (2017) Synergy between Prkdc and Trp53 regulates stem cell proliferation and GI-ARS after irradiation. Cell Death Differ 24(11):1853–1860.

50. Mangerich A & Burkle, A (2012) Pleiotropic cellular functions of PARP1 in longevity and aging: genome maintenance meets inflammation. Oxid Med Cell Longev 2012:321653.

51. Park J, et al. (2015) Mitochondrial ROS govern the LPS-induced pro-inflammatory response in microglia cells by regulating MAPK and NF-kappaB pathways. Neurosci Lett 584:191–196.

52. Kay AM, Simpson CL, & Stewart JA, Jr. (2016) The Role of AGE/RAGE Signaling in Diabetes-Mediated Vascular Calcification. J Diabetes Res 2016:6809703.

53. Goldin A, Beckman JA, Schmidt AM, & Creager MA (2006) Advanced glycation end products: sparking the development of diabetic vascular injury. Circulation 114(6):597–605.

54. Coughlan MT, et al. (2009) RAGE-induced cytosolic ROS promote mitochondrial superoxide generation in diabetes. J Am Soc Nephrol 20(4):742–752.

55. Daffu G, et al. (2013) Radical roles for RAGE in the pathogenesis of oxidative stress in cardiovascular diseases and beyond. Int J Mol Sci 14(10):19891–19910.

56. Mahali S, Raviprakash N, Raghavendra PB, & Manna SK (2011) Advanced glycation end products (AGEs) induce apoptosis via a novel pathway: involvement of Ca2+ mediated by interleukin-8 protein. J Biol Chem 286(40):34903–34913.

57. Sharifi-Sanjani M, et al. (2017) Cardiomyocyte-Specific Telomere Shortening is a Distinct Signature of Heart Failure in Humans. J Am Heart Assoc 6(9).

58. Hagen CM, et al. (2013) MT-CYB mutations in hypertrophic cardiomyopathy. Mol Genet Genomic Med 1(1):54–65.

