## Supplementary material for "Comparative gene expression in cells competent in or lacking DNA-PKcs kinase activity following etoposide exposure reveal differences in gene expression associated with histone modifications, inflammation, cell cycle regulation, Wnt signaling, and differentiation": https://ashleylab.nmsu.edu/supplementary-data/

### Supplemental Figures

**Supplemental Figure 1 :** Differentially regulated genes with major biological pathways in DNA-PKcs variants genotypes followed by  etoposide exposure.

Upregulated genes in KR cells compared to WT

S 1A

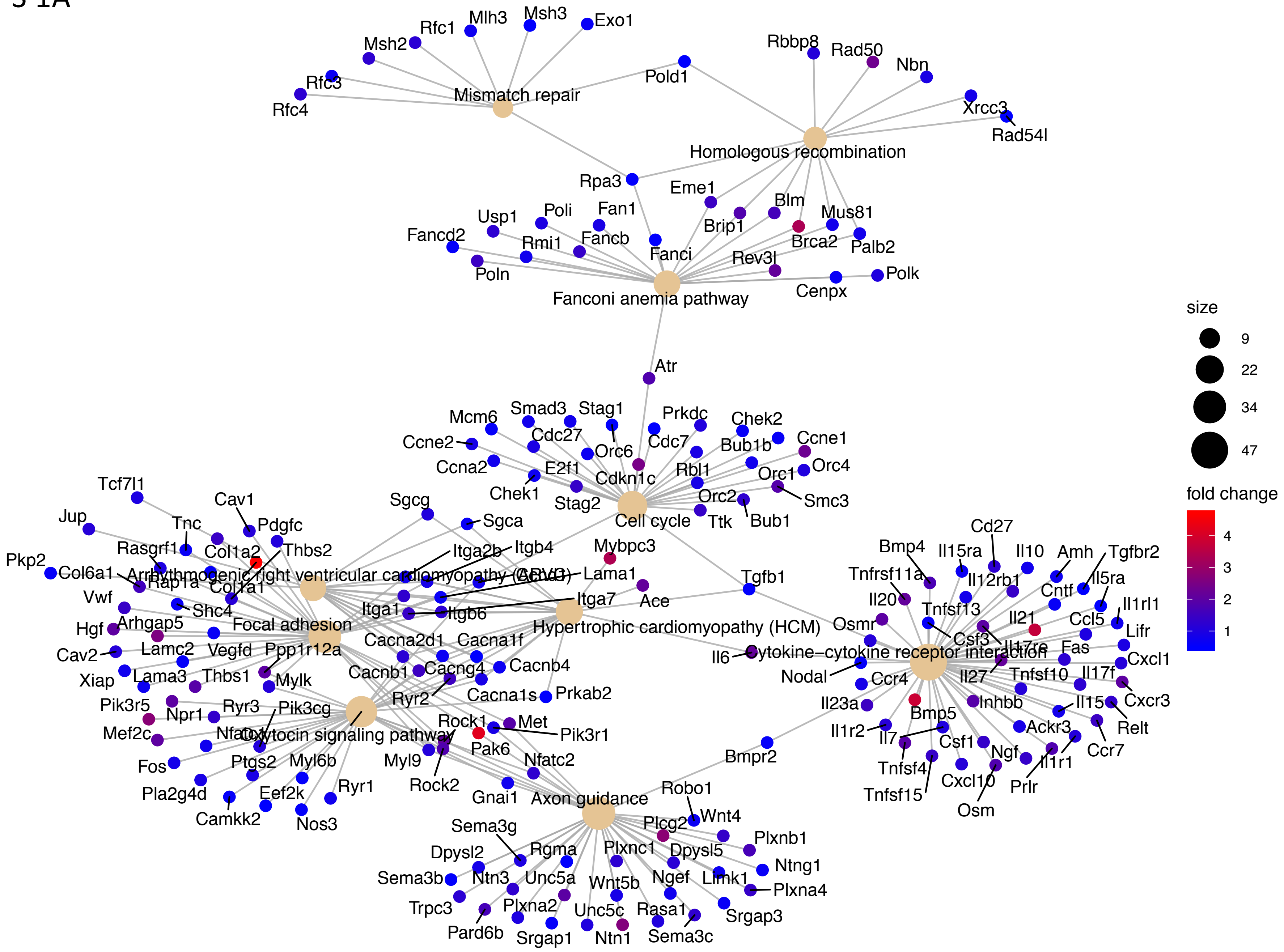

Down regulated genes in KR cells compared to WT

S 1B

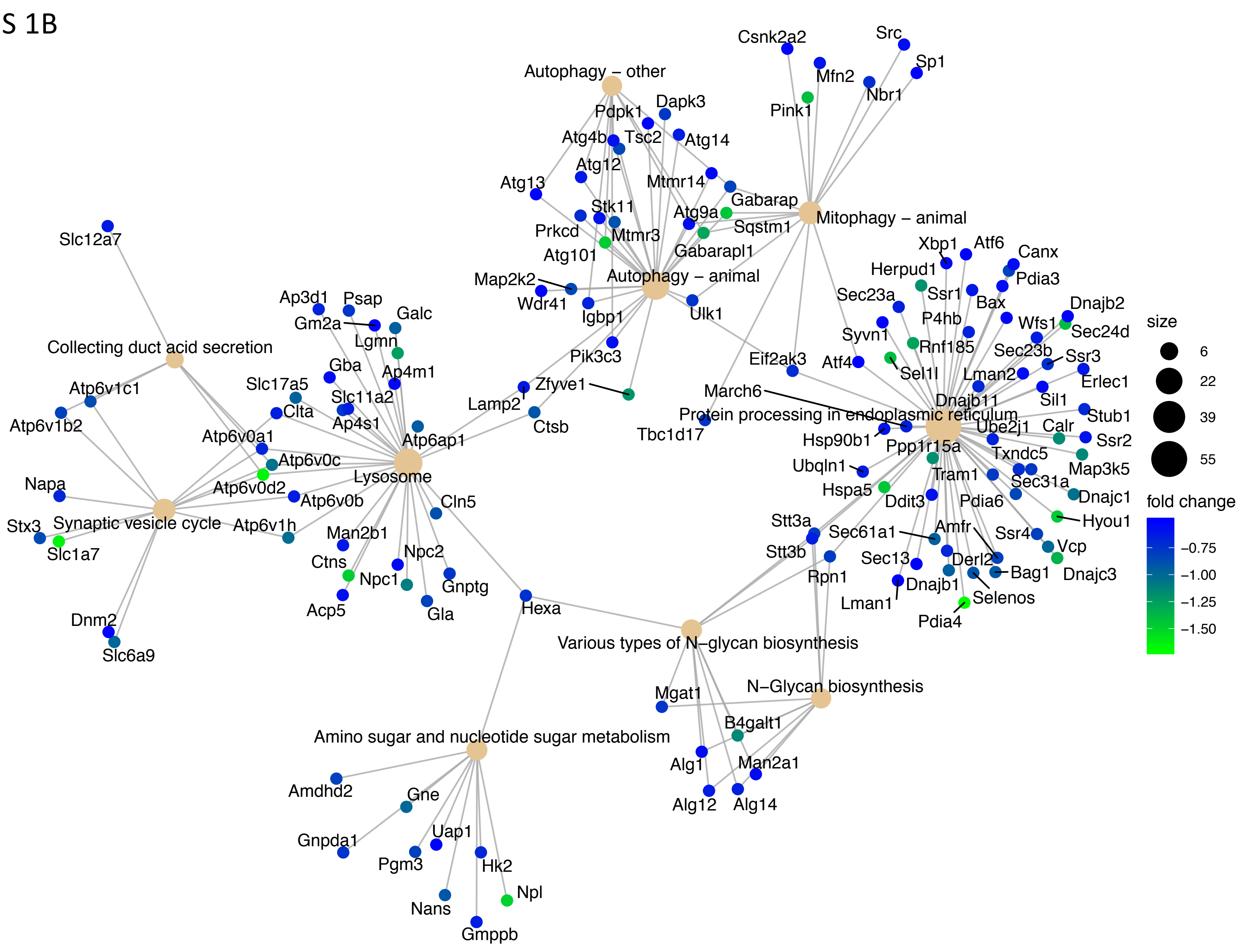

### Upregulated genes in WT cells after etoposide exposure

S 1C

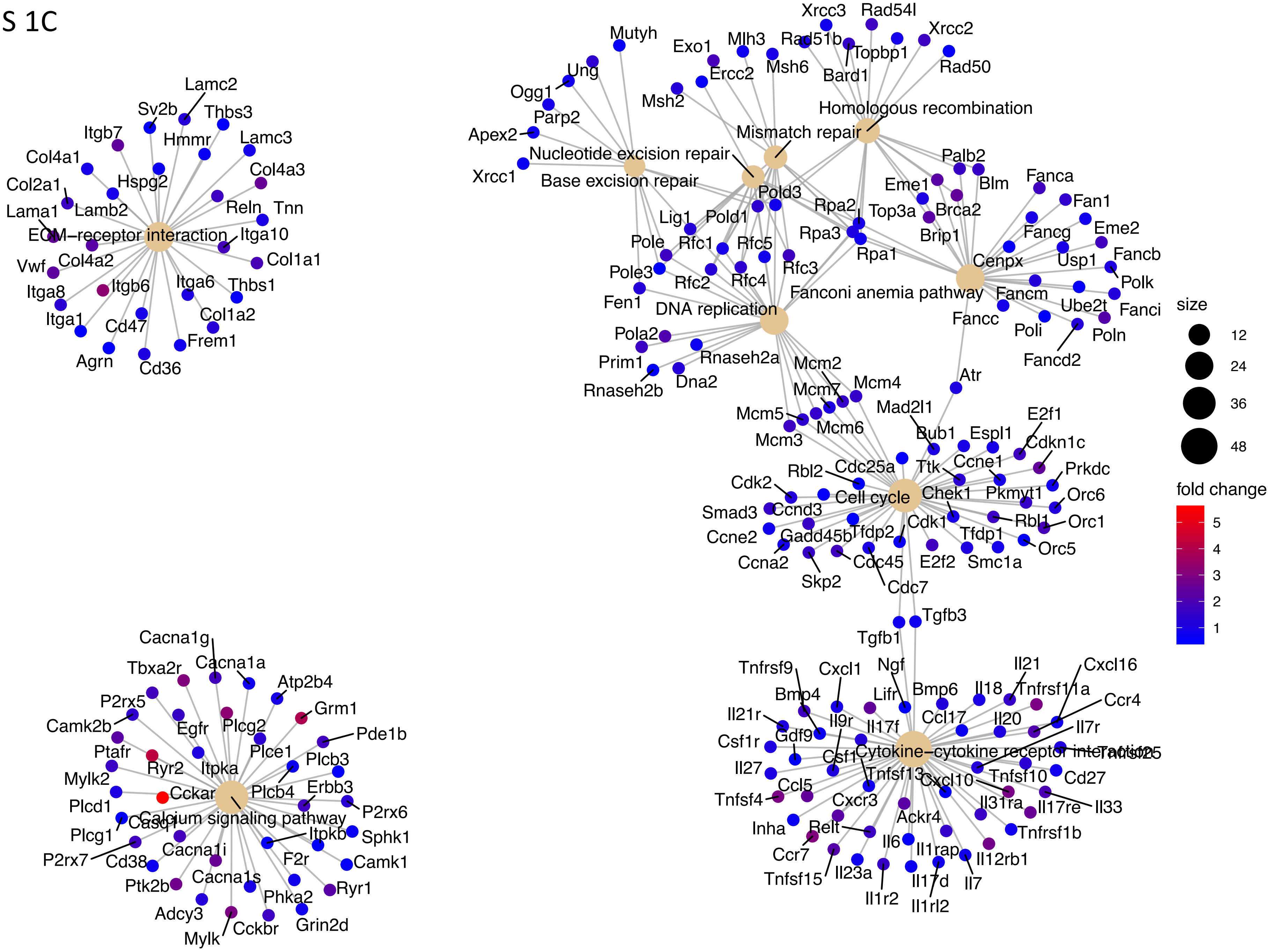

### Down regulated genes in WT cells after etoposide exposure

## S 1D

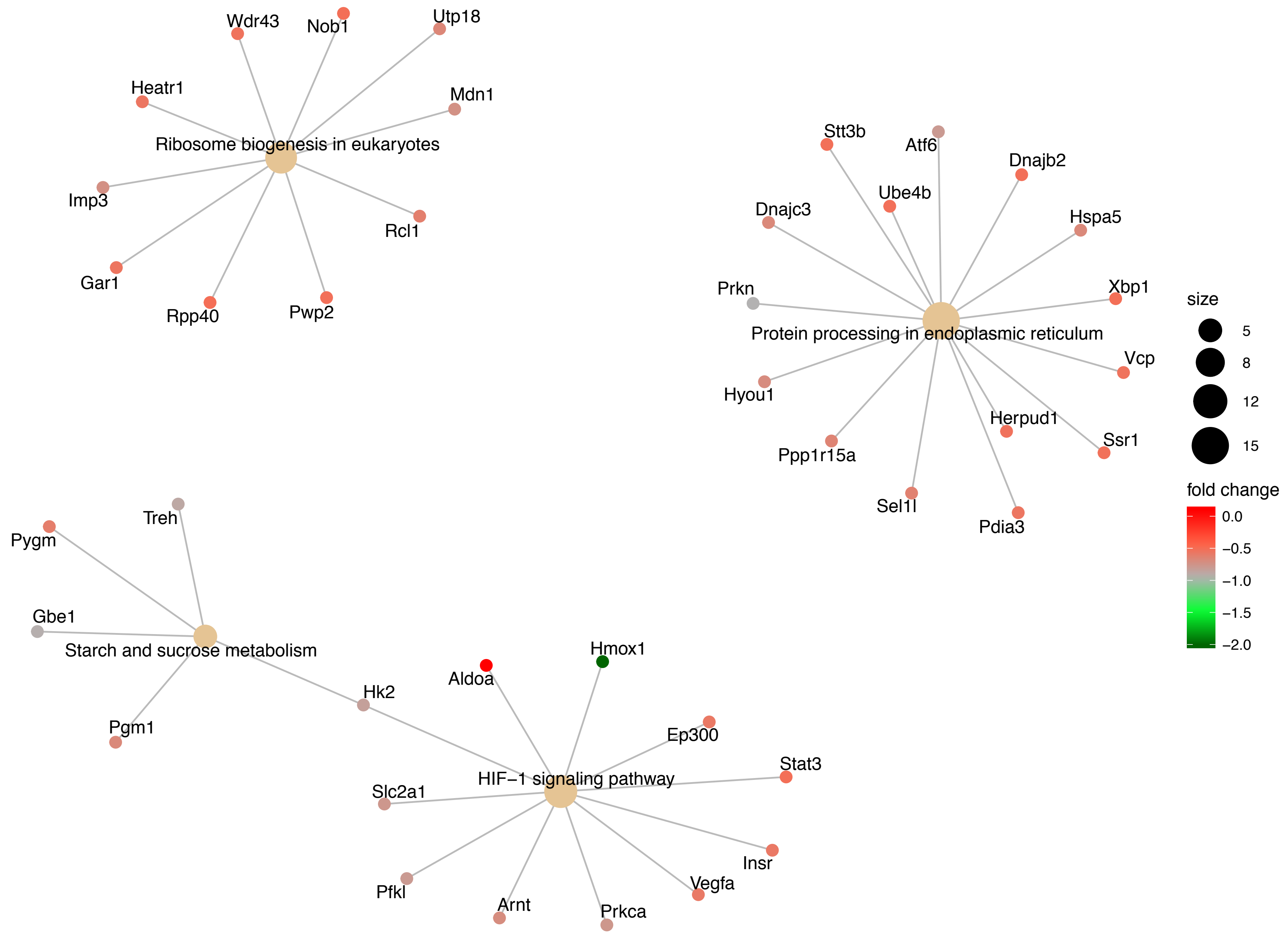

Upregulated genes in KR cells after etoposide exposure

S 1E

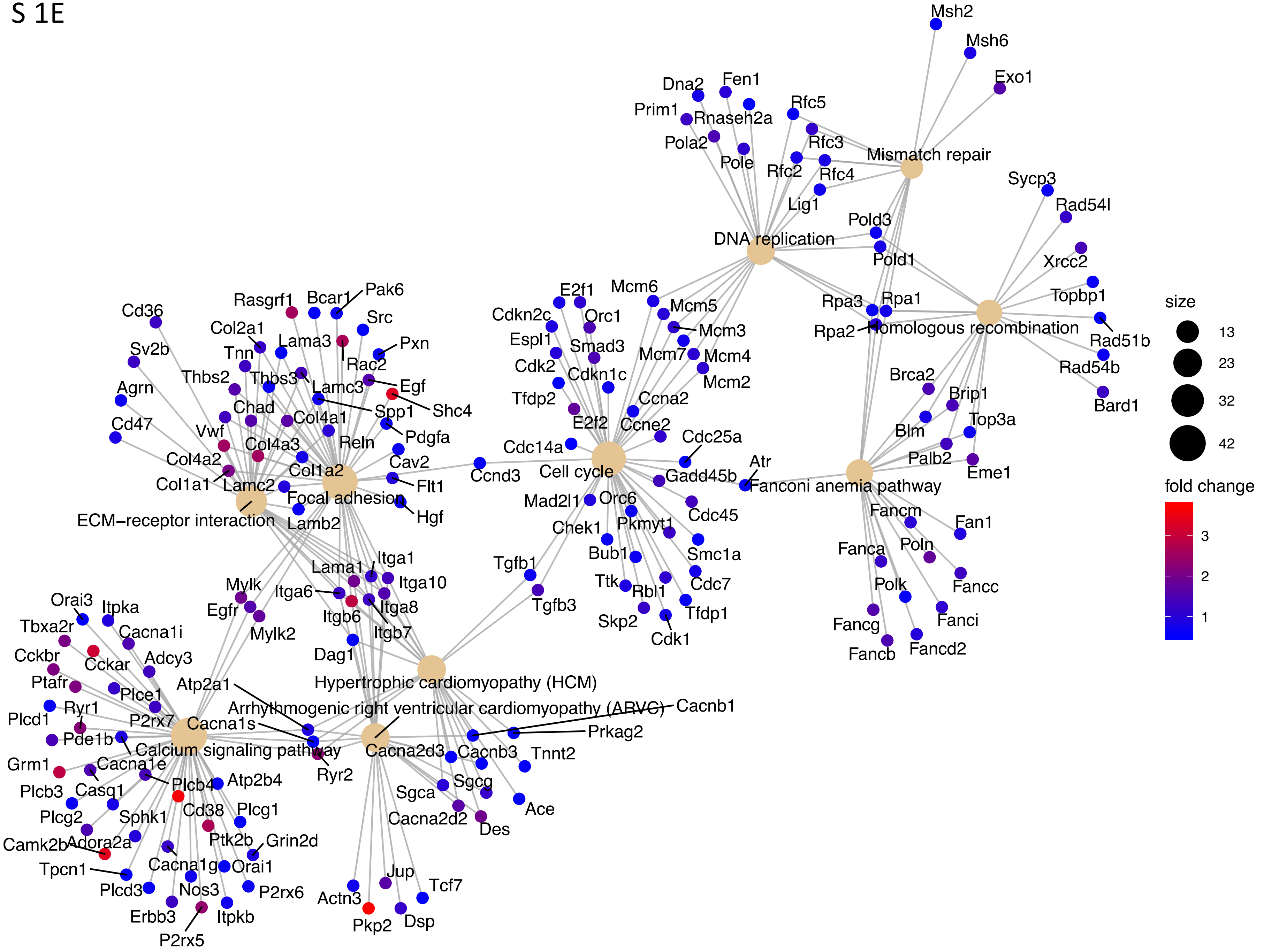

Down regulated genes in KR cells after etoposide exposure

S 1F

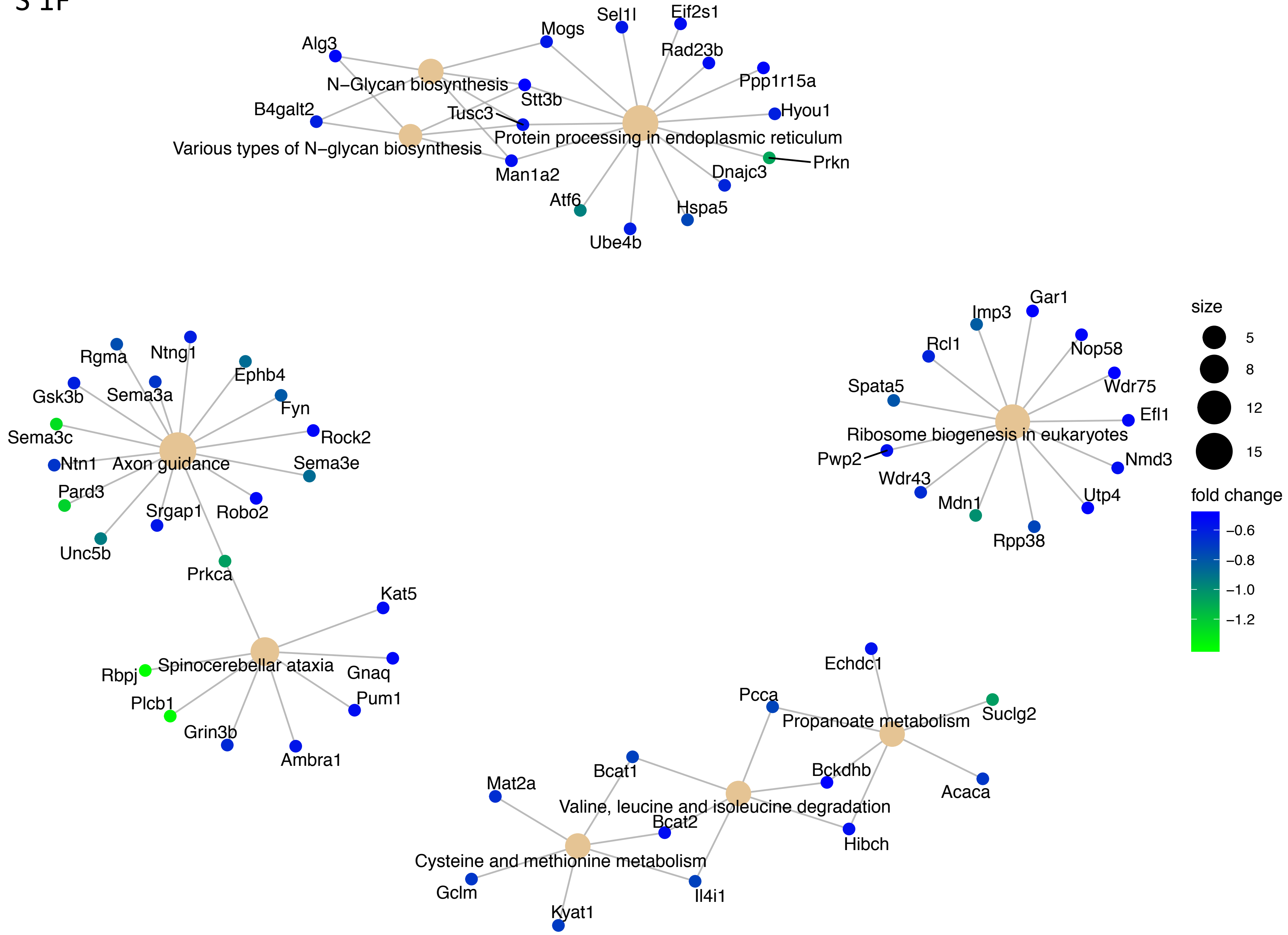

Upregulated genes in etoposide treated KR cells compared to WT

S 1G

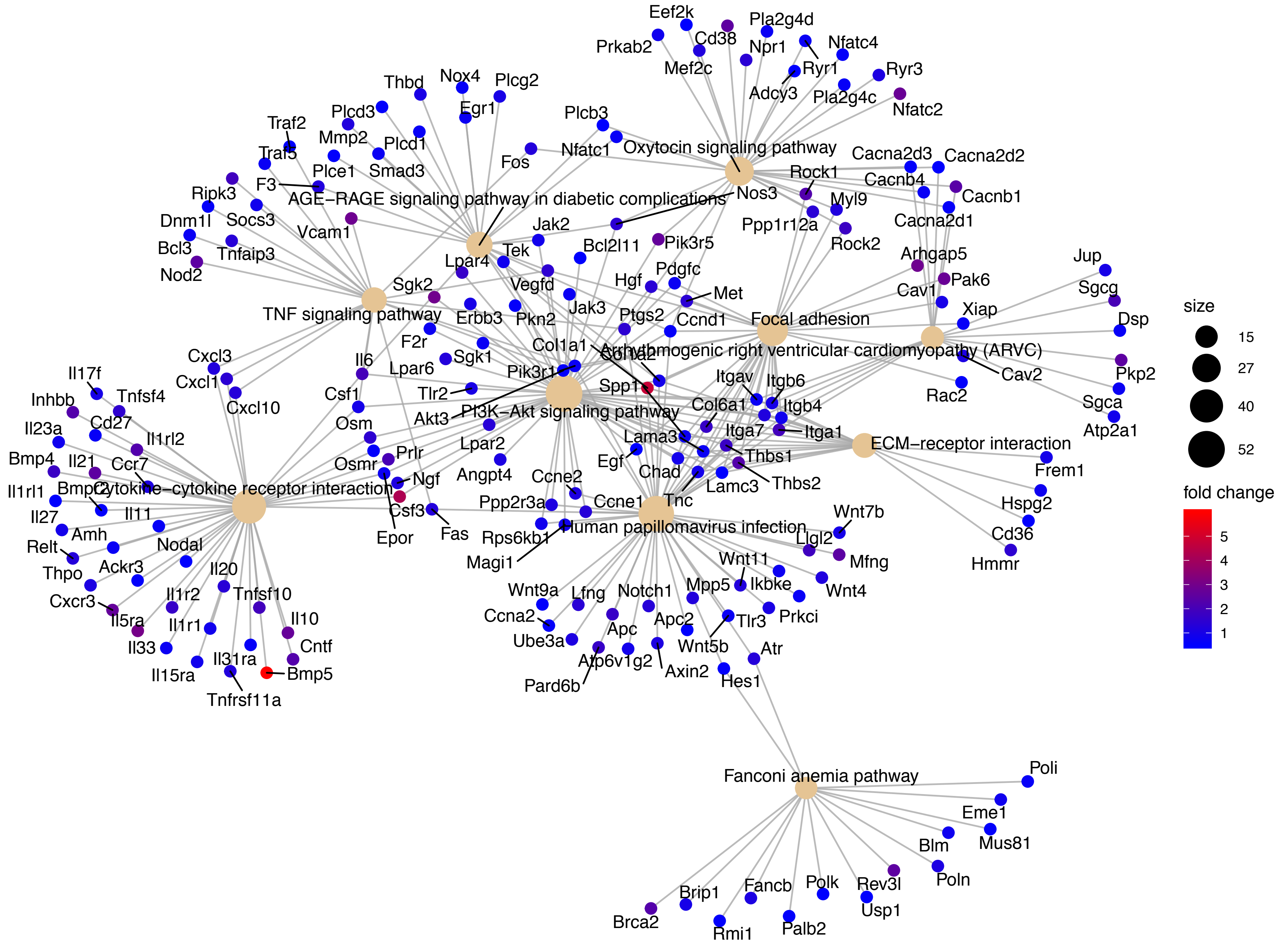

Down regulated genes in etoposide treated KR cells compared to WT

S 1H

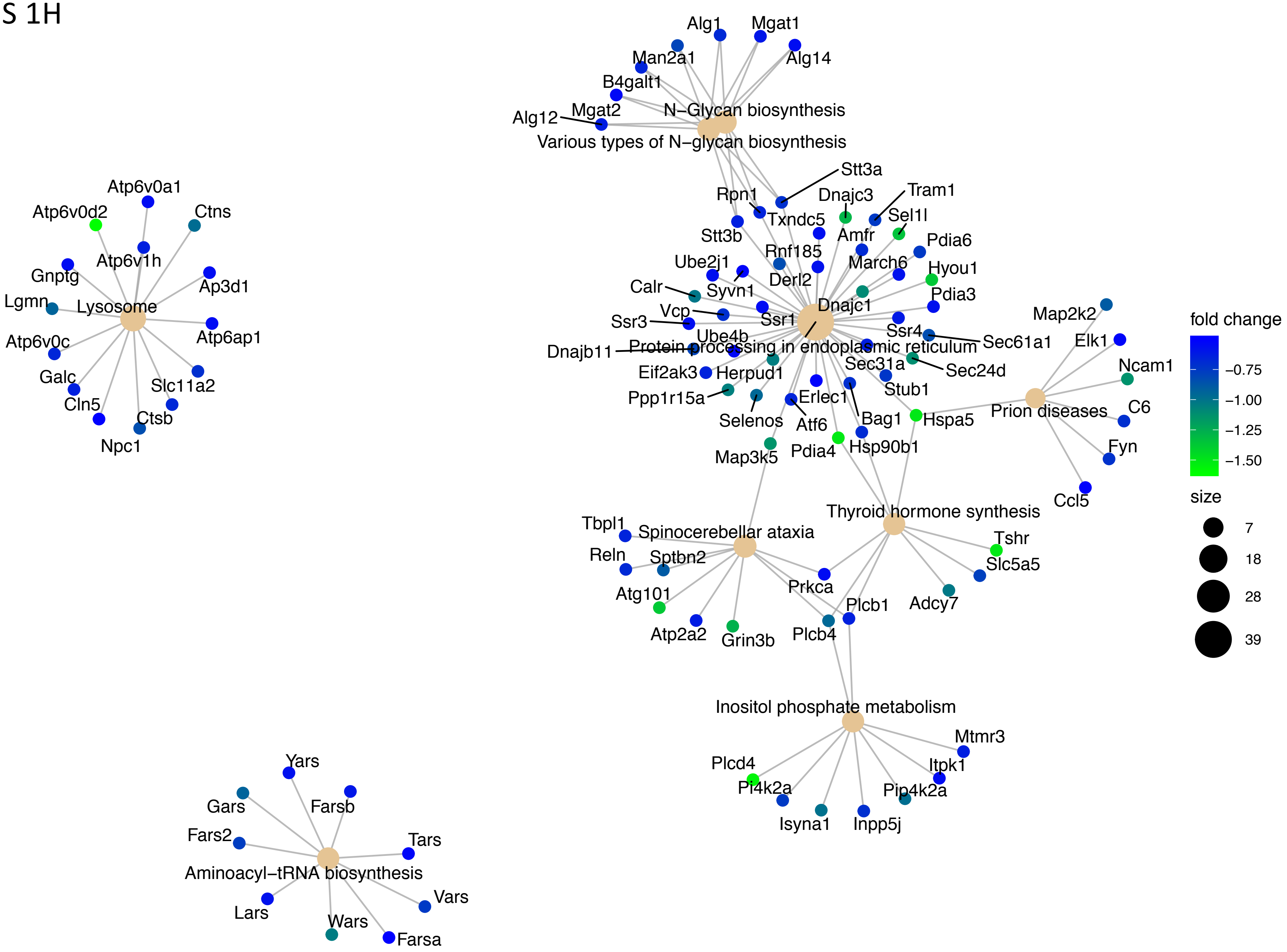

**Supplemental Figure 2 :** Nearest neighbor plots represents major differentially regulated biological pathways in DNA-PKcs variants genotypes followed by  etoposide exposure.

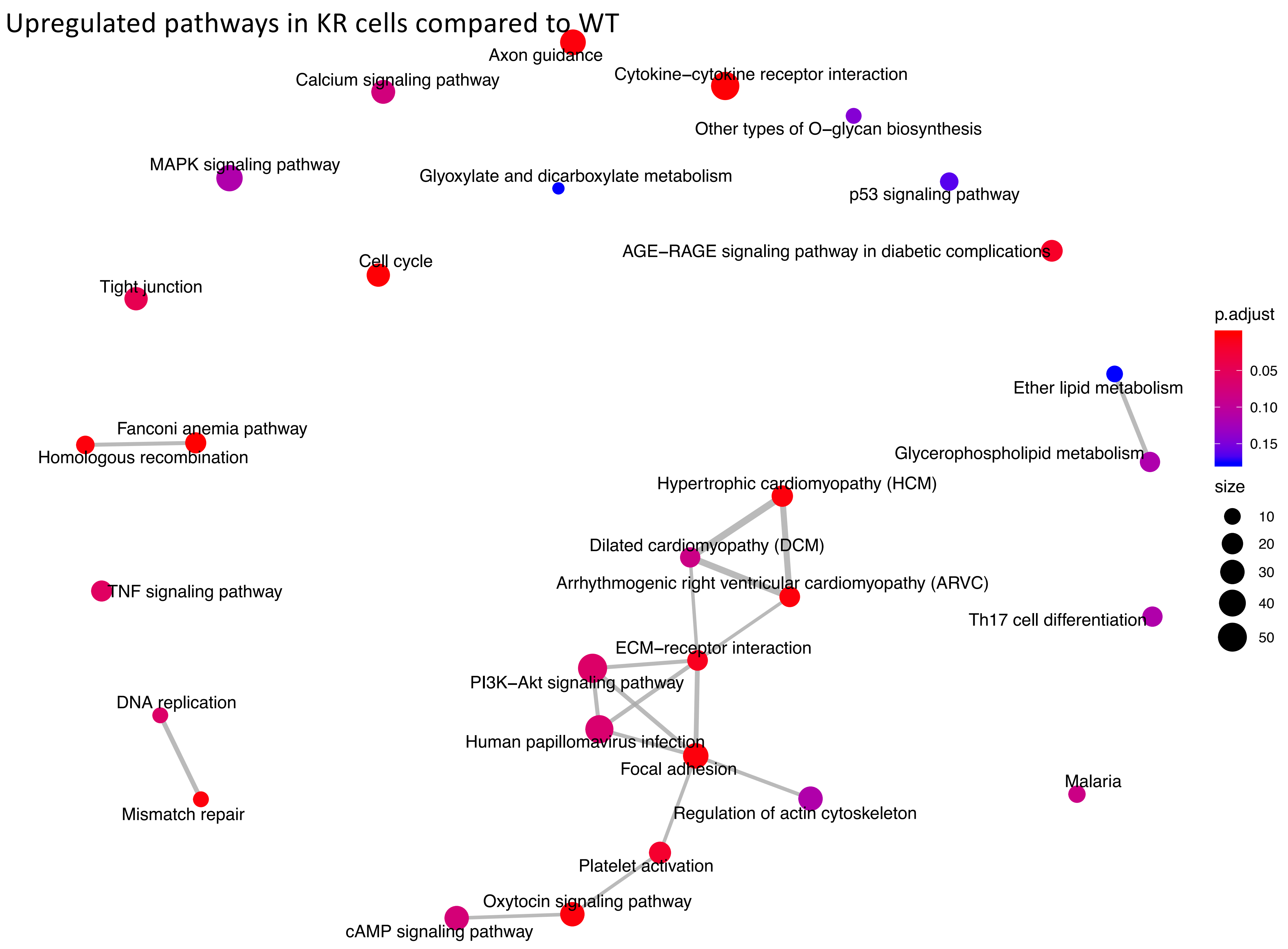
Upregulated pathways in KR cells compared to WT
Axon guidance
Calcium signaling pathway
Cytokine−cytokine receptor interaction
Other types of O−glycan biosynthesis
MAPK signaling pathway
Glyoxylate and dicarboxylate metabolism
p53 signaling pathway
AGE−RAGE signaling pathway in diabetic complications
Cell cycle
Tight junction
p.adjust
0.05
0.10
0.15
Ether lipid metabolism
Fanconi anemia pathway
Homologous recombination
Glycerophospholipid metabolism
Hypertrophic cardiomyopathy (HCM)
size
10
20
30
40
50
Dilated cardiomyopathy (DCM)
Arrhythmogenic right ventricular cardiomyopathy (ARVC)
TNF signaling pathway
Th17 cell differentiation
ECM−receptor interaction
PI3K−Akt signaling pathway
DNA replication
Human papillomavirus infection
Focal adhesion
Malaria
Mismatch repair
Regulation of actin cytoskeleton
Platelet activation
Oxytocin signaling pathway
cAMP signaling pathway

Down regulated pathways in KR cells compared to WT

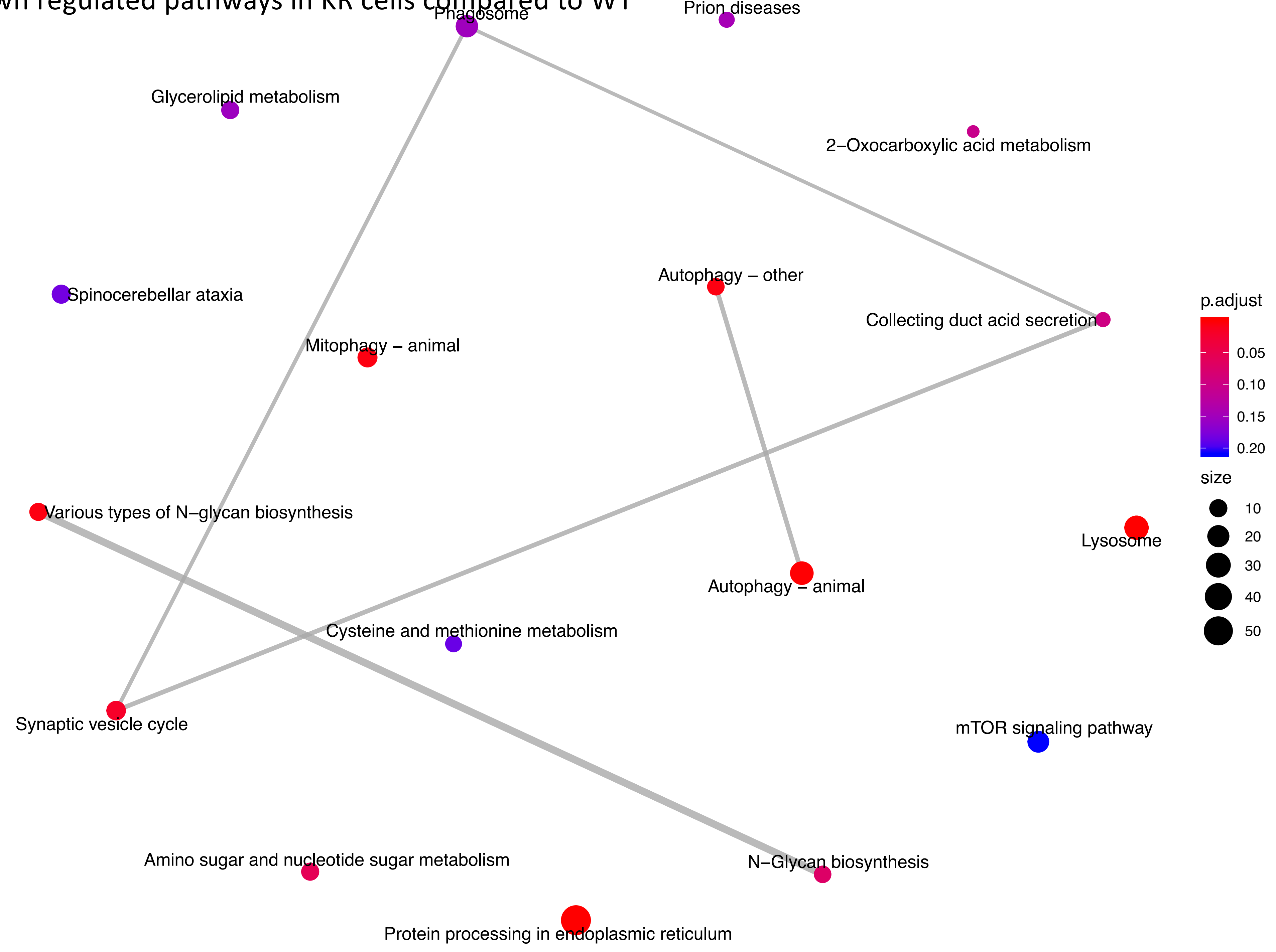

Upregulated pathways in WT cells after etoposide exposure

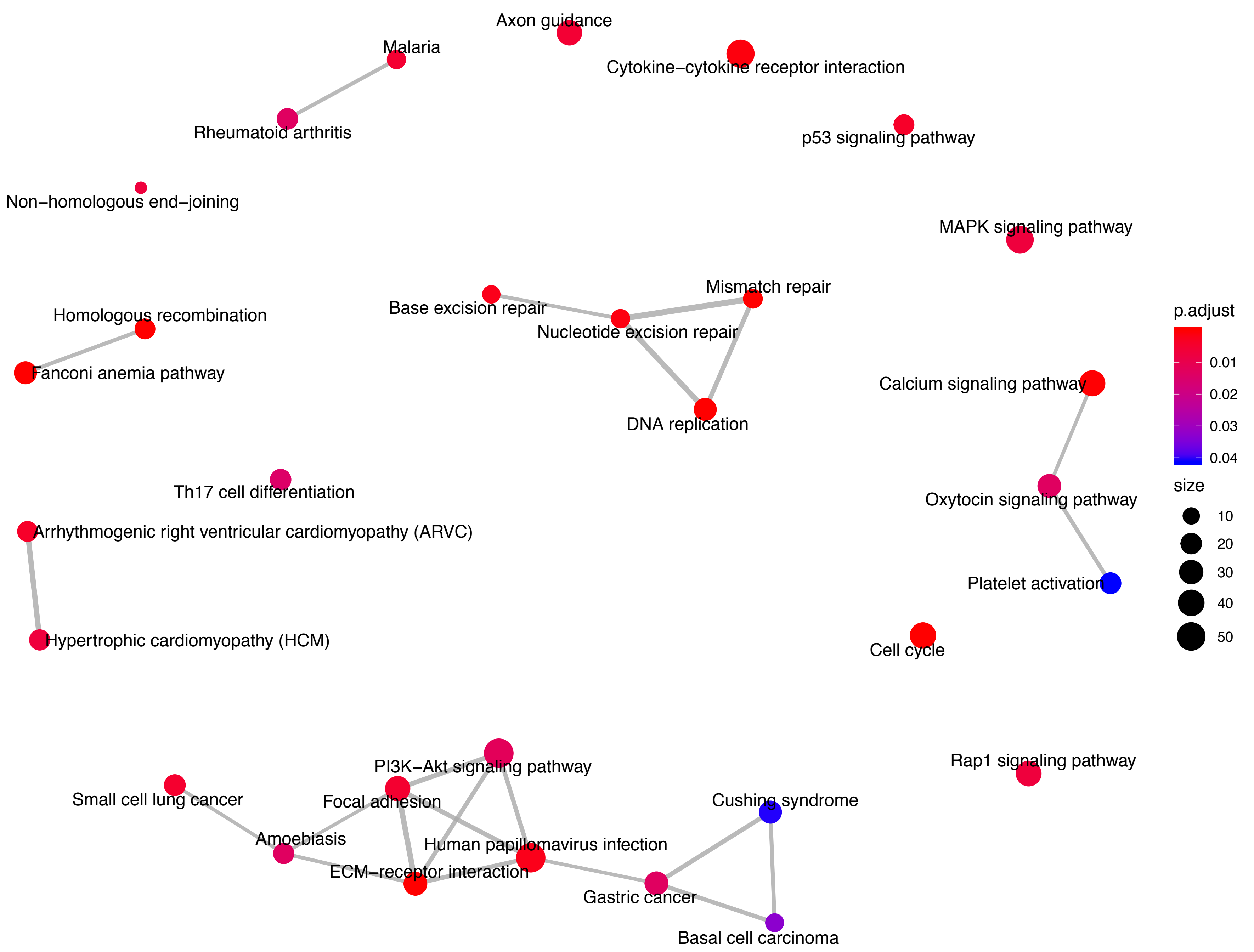

Down regulated pathways in WT cells after etoposide exposure

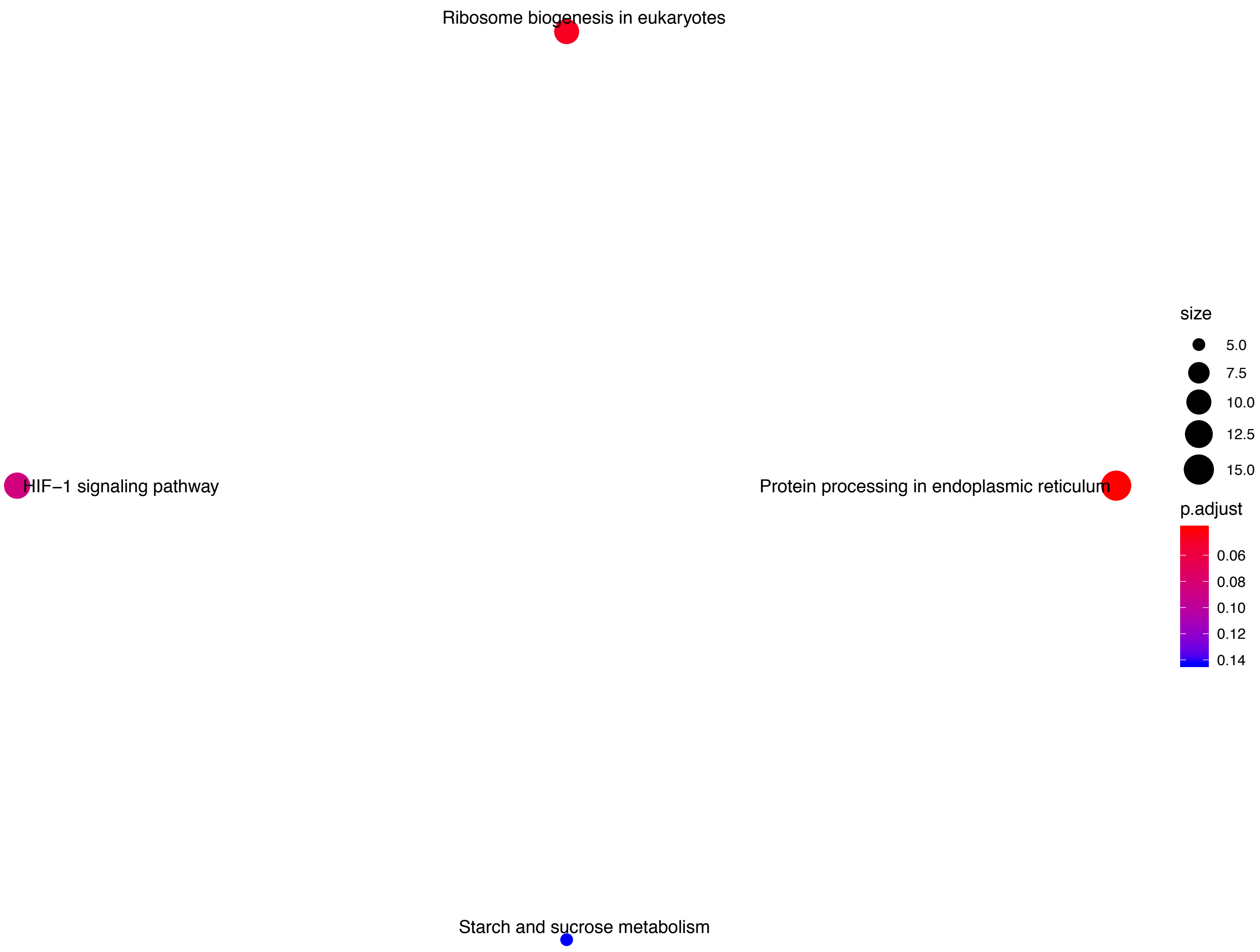

Upregulated pathways in KR cells after etoposide exposure

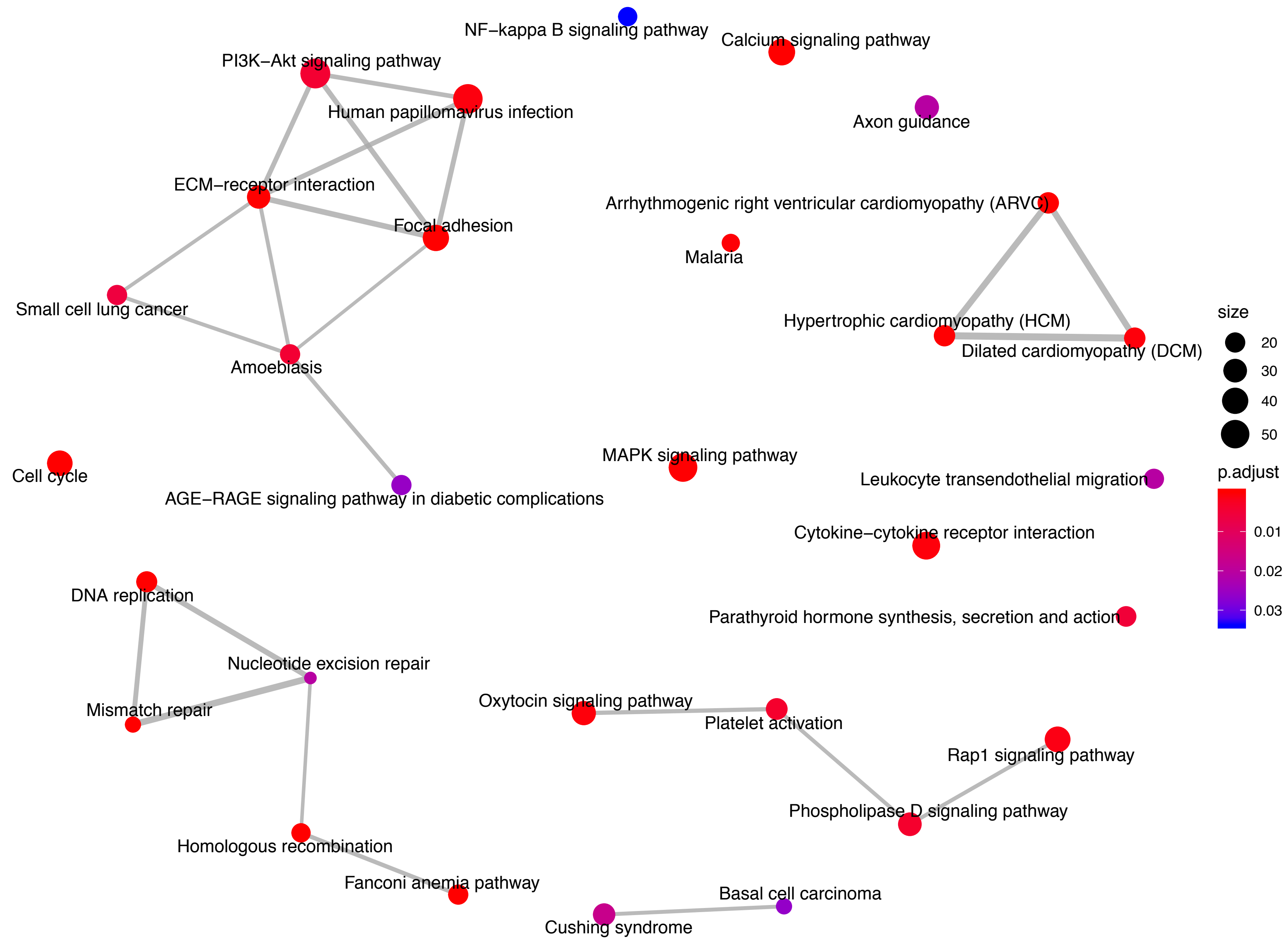

Down regulated pathways in KR cells after etoposide exposure

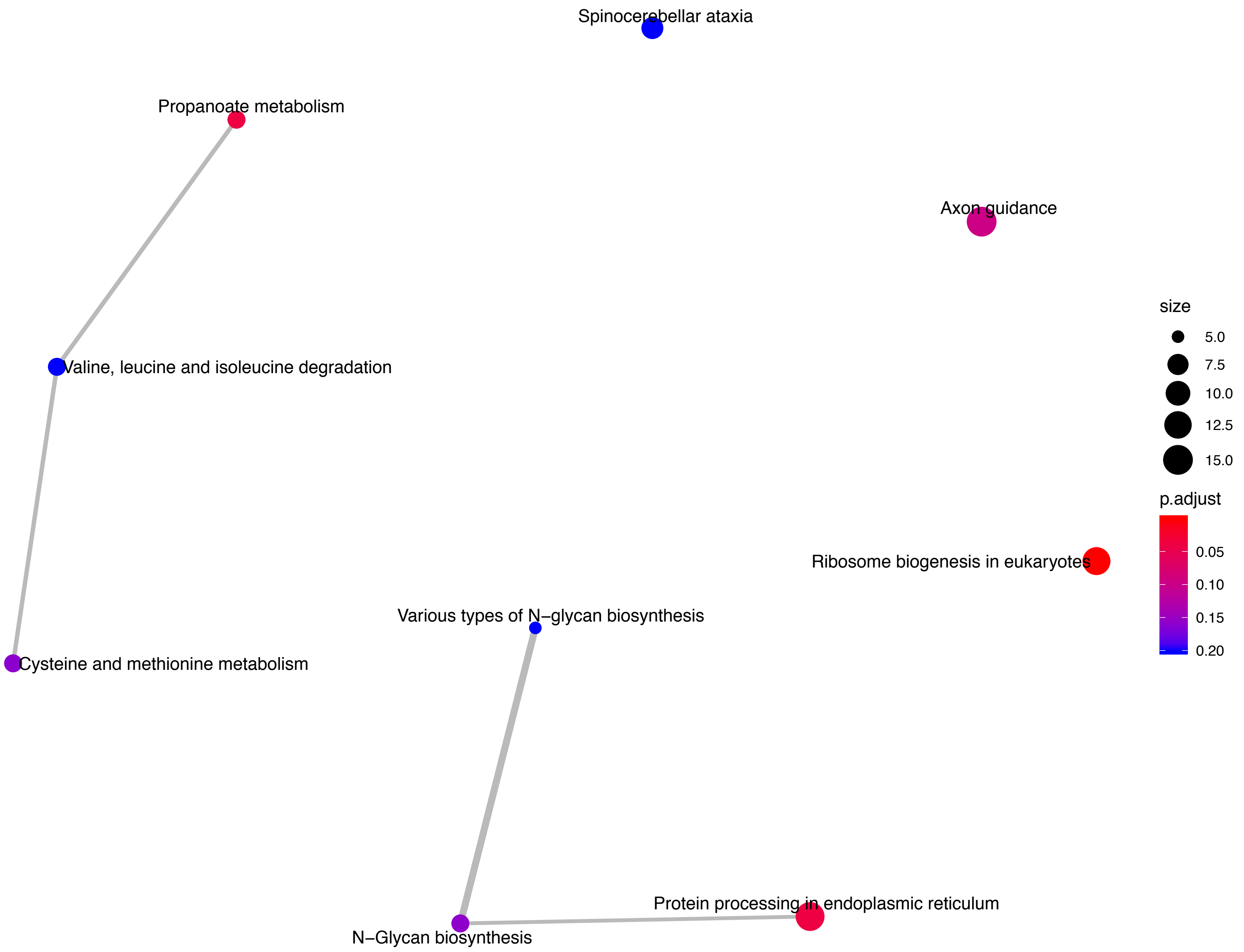

Upregulated pathways in etoposide treated KR cells compared to WT cells

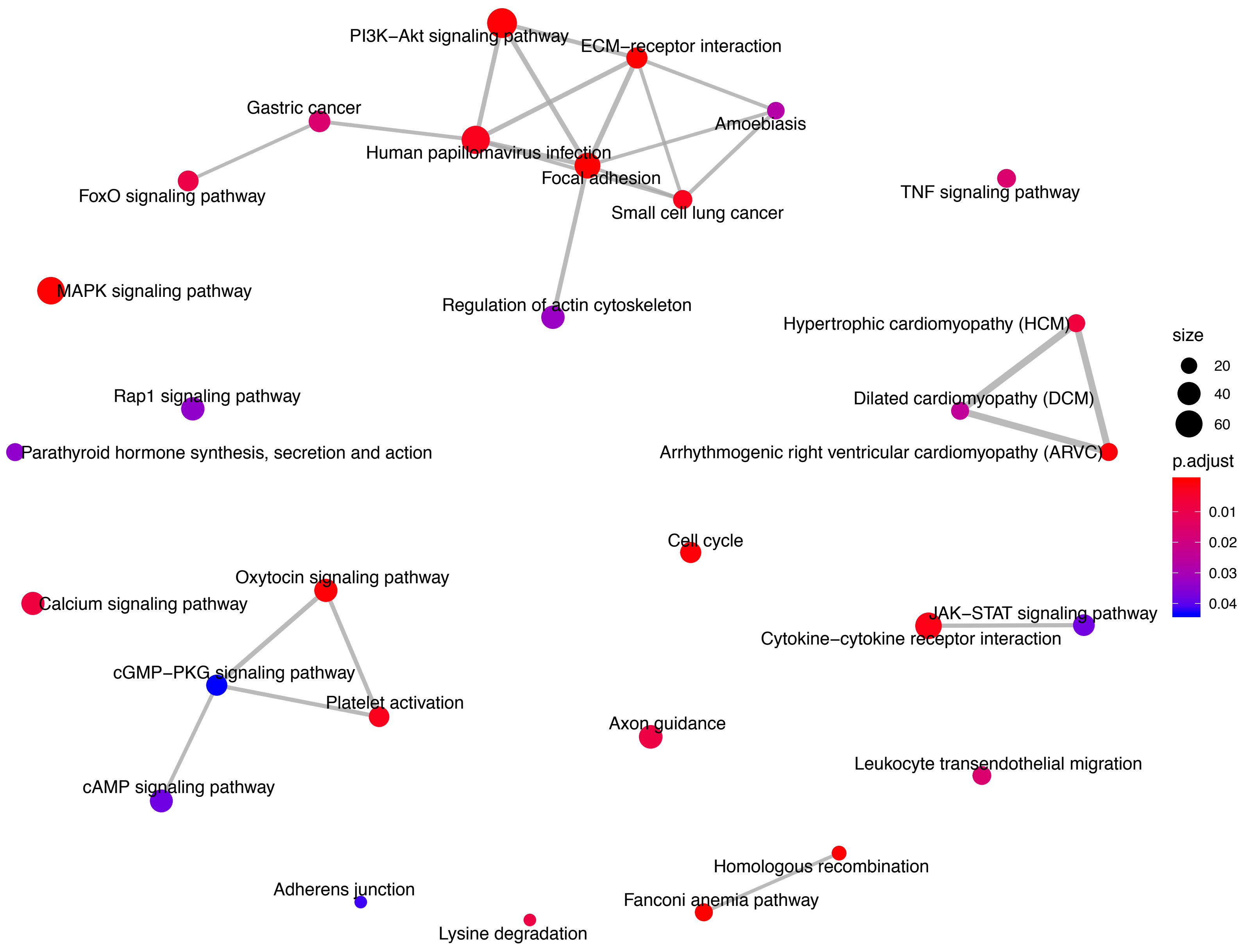

Down regulated pathways in etoposide treated KR cells compared to WT cells

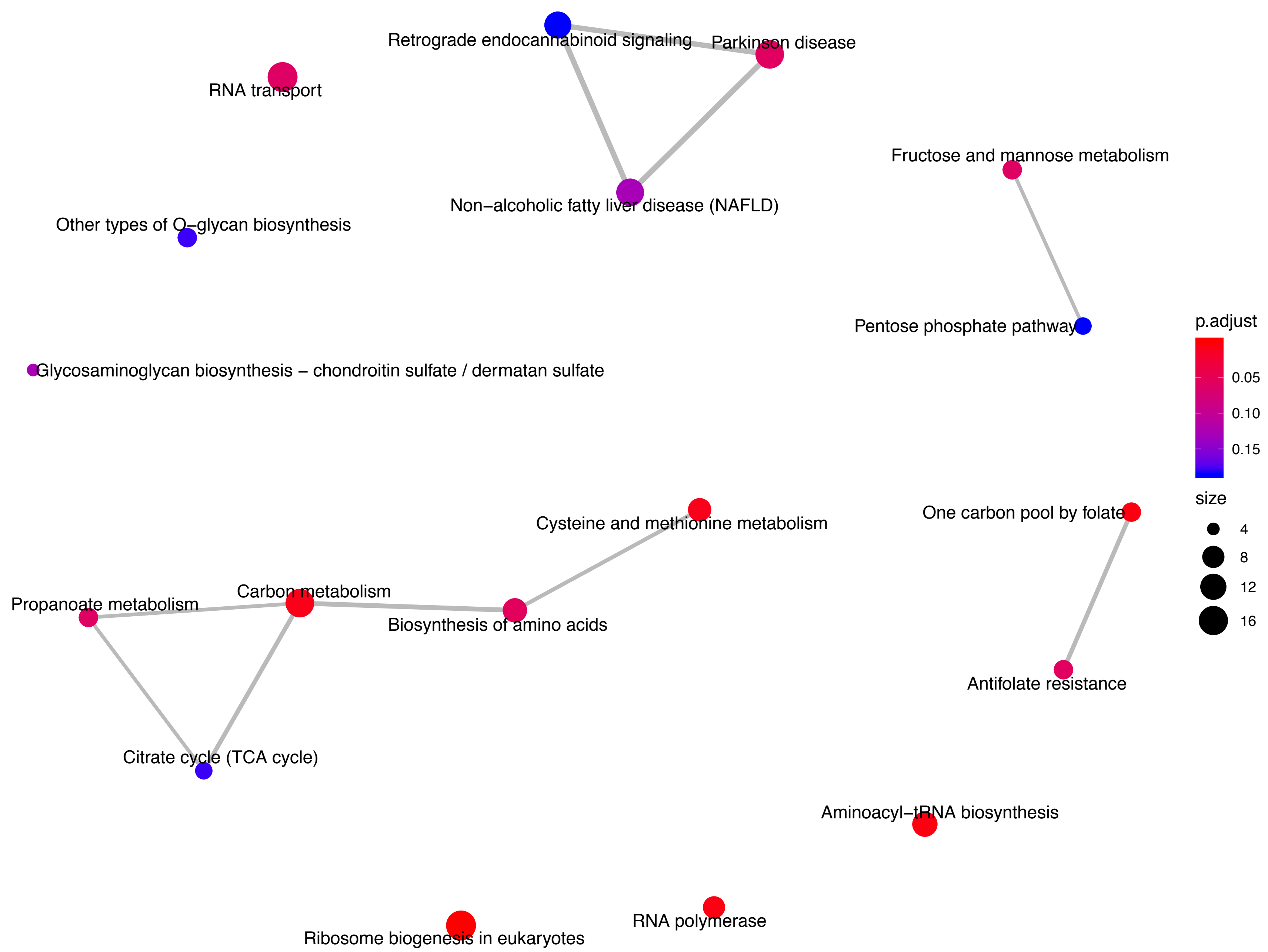

**Supplementary Table 2. Etoposide-induced differences in master regulators in WT cells**

| **Master Regulator** | **Molecule Type** | **Predicted Activation** | **p-value of overlap** |
|---|---|---|---|
| Master Regulator | Molecule Type | Predicted Activa | p-value of overla |
| denileukin diftitox | biologic drug | | 1.68E-14 |
| cdc25C phosphatase (211-221) | biologic drug | | 1.34E-08 |
| vasopressins | biologic drug | Activated | 1.64E-14 |
| desmopressin | biologic drug | Activated | 3.39E-10 |
| desmopressin | biologic drug | Activated | 0.000273 |
| interferon alfa-n1 | biologic drug | Activated | 0.000387 |
| PEG-interferon alfa-2a | biologic drug | Activated | 0.000387 |
| recombinant interferon | biologic drug | Activated | 0.000387 |
| l-asparaginase | biologic drug | Inhibited | 7.43E-27 |
| bivalirudin | biologic drug | Inhibited | 0.000000074 |
| heparin | chemical - endogenous mammalian | | 6.69E-15 |
| 9,13-di-cis-retinoic acid | chemical - endogenous mammalian | | 9.12E-14 |
| 4-hydroxy-all-trans-retinoic acid | chemical - endogenous mammalian | | 2.67E-13 |
| beta-apo-14'-carotenal | chemical - endogenous mammalian | | 2.67E-13 |
| all-trans-4-Oxo-retinoic acid | chemical - endogenous mammalian | | 2.67E-13 |
| 18-hydroxy-retinoic acid | chemical - endogenous mammalian | | 2.67E-13 |
| epinephrine | chemical - endogenous mammalian | | 2.93E-13 |
| formic acid | chemical - endogenous mammalian | | 2.78E-11 |
| propionic acid | chemical - endogenous mammalian | | 3.02E-11 |
| pentanoic acid | chemical - endogenous mammalian | | 3.02E-11 |
| ergocalciferol | chemical - endogenous mammalian | | 0.000000108 |
| calcifediol | chemical - endogenous mammalian | | 0.00000374 |
| cardiolipin | chemical - endogenous mammalian | | 0.00000608 |
| taurodeoxycholic acid | chemical - endogenous mammalian | | 0.0000109 |
| L-carnitine | chemical - endogenous mammalian | | 0.000147 |
| 10E,12Z-octadecadienoic acid | chemical - endogenous mammalian | | 0.00019 |
| 8-epi-prostaglandin F2alpha | chemical - endogenous mammalian | | 0.000764 |
| 19,20-epoxydocosapentaenoic acid | chemical - endogenous mammalian | | 0.00148 |
| 8,9-epoxyeicosatrienoic acid | chemical - endogenous mammalian | | 0.00563 |
| trigonelline | chemical - endogenous mammalian | | 0.0251 |
| SAICAR | chemical - endogenous mammalian | Activated | 7.83E-19 |
| estrone sulfate | chemical - endogenous mammalian | Activated | 1.93E-14 |
| 4-hydroxy-N-desmethyltamoxifen | chemical - endogenous mammalian | Activated | 2.29E-14 |
| UDP-D-glucose | chemical - endogenous mammalian | Activated | 7.04E-14 |
| pregnenolone sulfate | chemical - endogenous mammalian | Activated | 6.56E-13 |
| indican | chemical - endogenous mammalian | Activated | 1.22E-12 |
| acetic acid | chemical - endogenous mammalian | Activated | 3.5E-12 |
| adenine | chemical - endogenous mammalian | Activated | 7.29E-12 |
| uric acid | chemical - endogenous mammalian | Activated | 1.38E-11 |
| acetylcholine | chemical - endogenous mammalian | Activated | 3.82E-10 |
| bombesin | chemical - endogenous mammalian | Activated | 4.43E-08 |

| | | | |
|---|---|---|---|
| androstenedione | chemical - endogenous mammalian | Activated | 0.00000213 |
| 17-hydroxyprogesterone | chemical - endogenous mammalian | Activated | 0.0000156 |
| lactosylceramide | chemical - endogenous mammalian | Activated | 0.000601 |
| phospholipid | chemical - endogenous mammalian | Activated | 0.000912 |
| lysophosphatidylinositol | chemical - endogenous mammalian | Activated | 0.000987 |
| 2'3'-cyclic guanosine monophosphate-adenosine monophosphate | chemical - endogenous mammalian | Activated | 0.00184 |
| glycosaminoglycan | chemical - endogenous mammalian | Inhibited | 1.43E-14 |
| trehalose | chemical - endogenous mammalian | Inhibited | 1.44E-13 |
| chloride | chemical - endogenous mammalian | Inhibited | 2.73E-12 |
| desmethylazelastine | chemical - endogenous mammalian | Inhibited | 4.85E-10 |
| nitro-conjugated linoleic acid | chemical - endogenous mammalian | Inhibited | 0.00000614 |
| 9-nitrooleic acid | chemical - endogenous mammalian | Inhibited | 0.00000614 |
| 3-hydroxybutyric acid | chemical - endogenous mammalian | Inhibited | 0.0138 |
| colcemid | chemical - endogenous non-mammalian | | 9.59E-12 |
| anacardic acid | chemical - endogenous non-mammalian | | 2.56E-10 |
| imperatorin | chemical - endogenous non-mammalian | | 0.00000524 |
| phosphoramidon | chemical - endogenous non-mammalian | | 0.00000906 |
| bis(4-hydroxycinnamoyl)methane | chemical - endogenous non-mammalian | | 0.00549 |
| genipin | chemical - endogenous non-mammalian | | 0.00563 |
| rhodioloside | chemical - endogenous non-mammalian | | 0.0104 |
| ajoene | chemical - endogenous non-mammalian | Activated | 1.34E-08 |
| bis(4-hydroxycinnamoyl)methane | chemical - endogenous non-mammalian | Inhibited | 1.62E-15 |
| bis(4-hydroxycinnamoyl)methane | chemical - endogenous non-mammalian | Inhibited | 2.14E-14 |
| helenalin | chemical - endogenous non-mammalian | Inhibited | 1.23E-13 |
| soraphen-A1alpha | chemical - endogenous non-mammalian | Inhibited | 1.37E-13 |
| sesamin | chemical - endogenous non-mammalian | Inhibited | 5.15E-11 |
| flavone | chemical - endogenous non-mammalian | Inhibited | 2.7E-09 |
| decursin | chemical - endogenous non-mammalian | Inhibited | 7.29E-09 |
| fumagillin | chemical - endogenous non-mammalian | Inhibited | 7.75E-09 |

| | | | |
|---|---|---|---|
| geraniol | chemical - endogenous non-mammalian | Inhibited | 0.000987 |
| dorsomorphin | chemical - kinase inhibitor | | 1.07E-14 |
| IC261 | chemical - kinase inhibitor | | 4.55E-10 |
| CGP 74514A | chemical - kinase inhibitor | | 5.89E-09 |
| SU6657 | chemical - kinase inhibitor | | 0.00000039 |
| THZ1 | chemical - kinase inhibitor | | 0.0000117 |
| alpha-cyano-(3-ethoxy-4-hydroxy-5-phenylthiomethyl)cinnamamide | chemical - kinase inhibitor | | 0.0000159 |
| tyrphostin AG 1295 | chemical - kinase inhibitor | | 0.00188 |
| CBR-470-1 | chemical - kinase inhibitor | | 0.00778 |
| bisindolylmaleimide iv | chemical - kinase inhibitor | Inhibited | 3.58E-13 |
| 4-[(4'-chloro-2'-fluoro)phenylamino]-6,7-dimethoxyquinazoline | chemical - kinase inhibitor | Inhibited | 7.76E-13 |
| JNK-IN-8 | chemical - kinase inhibitor | Inhibited | 2.82E-12 |
| BB4 | chemical - kinase inhibitor | Inhibited | 1.62E-11 |
| Rp-cAMPS | chemical - kinase inhibitor | Inhibited | 9.75E-10 |
| PD 180970 | chemical - kinase inhibitor | Inhibited | 0.000000148 |
| SB203580 | chemical - kinase inhibitor | Inhibited | 0.000000879 |
| 6-aminopyrazolopyrimidine derivative compound II | chemical - kinase inhibitor | Inhibited | 0.0000167 |
| UM101 | chemical - kinase inhibitor | Inhibited | 0.00133 |
| THZ1 | chemical - kinase inhibitor | Inhibited | 0.00326 |
| catecholamine | chemical - other | Activated | 1.72E-11 |
| porphyrin | chemical - other | Inhibited | 8.53E-09 |
| Ac-YVAD-CMK | chemical - protease inhibitor | | 0.125 |
| MMPI-2 | chemical - protease inhibitor | Inhibited | 1.89E-12 |
| isoproterenol | chemical drug | | 8.1E-18 |
| trans-hydroxytamoxifen | chemical drug | | 8.95E-16 |
| acitretin | chemical drug | | 1.77E-15 |
| calcitriol | chemical drug | | 1.65E-14 |
| D-tubocurarine | chemical drug | | 2.9E-14 |
| nadolol | chemical drug | | 8.26E-14 |
| capsaicin | chemical drug | | 1.69E-13 |
| ephedrine | chemical drug | | 1.74E-12 |
| ardeparin | chemical drug | | 1.86E-12 |
| novobiocin | chemical drug | | 2.19E-12 |
| ribavirin | chemical drug | | 2.46E-12 |
| talazoparib | chemical drug | | 3.3E-12 |
| fluphenazine | chemical drug | | 5.63E-12 |
| adaphostin | chemical drug | | 1.22E-11 |
| sorafenib | chemical drug | | 2.35E-11 |
| perphenazine | chemical drug | | 4.85E-11 |
| bendamustine | chemical drug | | 7.27E-10 |
| chlorpropamide | chemical drug | | 0.000000002 |
| imipramine | chemical drug | | 2.39E-08 |

| | | |
|---|---|---|
| inecalcitol | chemical drug | 0.000000026 |
| azathioprine | chemical drug | 0.000000038 |
| pyrilamine | chemical drug | 5.21E-08 |
| ginkgolide B | chemical drug | 5.72E-08 |
| CC 401 | chemical drug | 0.000000105 |
| tazarotene | chemical drug | 0.000000107 |
| enzalutamide | chemical drug | 0.000000158 |
| avasimibe | chemical drug | 0.000000284 |
| doxorubicin | chemical drug | 0.000000288 |
| nicorandil | chemical drug | 0.00000042 |
| pobilukast | chemical drug | 0.000000436 |
| GLPG0187 | chemical drug | 0.00000055 |
| imipramine blue | chemical drug | 0.000000844 |
| bosutinib | chemical drug | 0.000000936 |
| NCX-4040 | chemical drug | 0.0000011 |
| repotrectinib | chemical drug | 0.00000189 |
| chiauranib | chemical drug | 0.00000769 |
| pirfenidone | chemical drug | 0.0000112 |
| acetovanillone | chemical drug | 0.0000181 |
| trans-hydroxytamoxifen | chemical drug | 0.0000349 |
| cerivastatin | chemical drug | 0.000112 |
| allopurinol | chemical drug | 0.000316 |
| triclosan | chemical drug | 0.000318 |
| eprosartan | chemical drug | 0.000451 |
| semaxinib | chemical drug | 0.000891 |
| felodipine | chemical drug | 0.0011 |
| indinavir | chemical drug | 0.00132 |
| tetomilast | chemical drug | 0.00132 |
| arofylline | chemical drug | 0.00132 |
| L 869298 | chemical drug | 0.00132 |
| ramipril | chemical drug | 0.00133 |
| CTS-1027 | chemical drug | 0.00188 |
| Yi-Gan San | chemical drug | 0.00188 |
| L-826,141 | chemical drug | 0.00249 |
| trimetrexate | chemical drug | 0.00328 |
| oxytetracycline | chemical drug | 0.00423 |
| fomepizole | chemical drug | 0.00549 |
| etodolac | chemical drug | 0.00549 |
| 6-mercaptopurine | chemical drug | 0.00778 |
| carvedilol | chemical drug | 0.00812 |
| enoximone | chemical drug | 0.00812 |
| palbociclib | chemical drug | 0.0104 |
| GR-MD-02 | chemical drug | 0.0107 |
| p38 MAP kinase inhibitor | chemical drug | 0.0107 |
| ticrynafen | chemical drug | 0.0107 |
| quinethazone | chemical drug | 0.0168 |
| tranexamic acid | chemical drug | 0.0173 |
| dexamethasone/tobramycin | chemical drug | 0.0434 |

| | | | |
|---|---|---|---|
| HDAC class I inhibitors | chemical drug | | 0.0434 |
| pralnacasan | chemical drug | | 0.0849 |
| alitretinoin | chemical drug | Activated | 8.41E-16 |
| mocetinostat | chemical drug | Activated | 2.21E-14 |
| OKI-179 | chemical drug | Activated | 3.42E-14 |
| cephaloridine | chemical drug | Activated | 6.96E-13 |
| bupropion | chemical drug | Activated | 5.57E-12 |
| SEL120 | chemical drug | Activated | 3.92E-10 |
| BCD-115 | chemical drug | Activated | 3.92E-10 |
| neomycin | chemical drug | Activated | 2.17E-09 |
| cirazoline | chemical drug | Activated | 4.95E-09 |
| terbutaline | chemical drug | Activated | 1.27E-08 |
| seocalcitol | chemical drug | Activated | 3.19E-08 |
| fenoterol | chemical drug | Activated | 8.09E-08 |
| nandrolone | chemical drug | Activated | 0.000000125 |
| estrogen | chemical drug | Activated | 0.000000223 |
| hydroxyurea | chemical drug | Activated | 0.00000206 |
| mibolerone | chemical drug | Activated | 0.00000231 |
| nandrolone decanoate | chemical drug | Activated | 0.00000715 |
| cinacalcet | chemical drug | Activated | 0.0000145 |
| testosterone enanthate | chemical drug | Activated | 0.0000185 |
| pargyline | chemical drug | Activated | 0.0000292 |
| mocetinostat | chemical drug | Activated | 0.0000615 |
| carboplatin | chemical drug | Activated | 0.000419 |
| phenylbutazone | chemical drug | Activated | 0.00107 |
| lestaurtinib | chemical drug | Inhibited | 1.95E-18 |
| PD 153035 | chemical drug | Inhibited | 1.08E-17 |
| cerulenin | chemical drug | Inhibited | 1.86E-17 |
| isoflurophate | chemical drug | Inhibited | 2.28E-16 |
| PLX7486 | chemical drug | Inhibited | 2.44E-15 |
| galunisertib | chemical drug | Inhibited | 3.45E-15 |
| bepridil | chemical drug | Inhibited | 3.47E-15 |
| lenalidomide | chemical drug | Inhibited | 4.08E-15 |
| gallium nitrate | chemical drug | Inhibited | 4.38E-15 |
| astemizole | chemical drug | Inhibited | 6.16E-15 |
| glesatinib | chemical drug | Inhibited | 1.09E-14 |
| thalidomide | chemical drug | Inhibited | 1.46E-14 |
| tioconazole | chemical drug | Inhibited | 3.05E-14 |
| escitalopram | chemical drug | Inhibited | 3.39E-14 |
| CEP-37440 | chemical drug | Inhibited | 3.6E-14 |
| sulconazole | chemical drug | Inhibited | 3.61E-14 |
| candesartan cilexetil | chemical drug | Inhibited | 5.85E-14 |
| altiratinib | chemical drug | Inhibited | 6.13E-14 |
| tranylcypromine | chemical drug | Inhibited | 7.72E-14 |
| caffeic acid phenethyl ester | chemical drug | Inhibited | 1.15E-13 |
| NOV1601 | chemical drug | Inhibited | 1.17E-13 |
| larotrectinib | chemical drug | Inhibited | 1.2E-13 |
| pomalidomide | chemical drug | Inhibited | 1.31E-13 |

| | | | |
|---|---|---|---|
| dasatinib | chemical drug | Inhibited | 1.58E-13 |
| pentamidine | chemical drug | Inhibited | 2.16E-13 |
| methoxsalen | chemical drug | Inhibited | 3.21E-13 |
| DS-6051b | chemical drug | Inhibited | 3.93E-13 |
| tivozanib | chemical drug | Inhibited | 3.98E-13 |
| etiracetam | chemical drug | Inhibited | 4.36E-13 |
| belizatinib | chemical drug | Inhibited | 6.42E-13 |
| acetazolamide | chemical drug | Inhibited | 6.43E-13 |
| strychnine | chemical drug | Inhibited | 6.96E-13 |
| ningetinib | chemical drug | Inhibited | 9.41E-13 |
| famitinib | chemical drug | Inhibited | 1.18E-12 |
| rolipram | chemical drug | Inhibited | 1.65E-12 |
| conteltinib | chemical drug | Inhibited | 1.7E-12 |
| VEGF inhibitor drug | chemical drug | Inhibited | 1.58E-11 |
| mimosine | chemical drug | Inhibited | 4.15E-11 |
| astemizole | chemical drug | Inhibited | 7.41E-11 |
| desloratadine | chemical drug | Inhibited | 8.18E-11 |
| pimozide | chemical drug | Inhibited | 8.2E-11 |
| loratadine | chemical drug | Inhibited | 8.53E-11 |
| axitinib | chemical drug | Inhibited | 8.64E-11 |
| calcitriol | chemical drug | Inhibited | 1.04E-10 |
| ticlopidine | chemical drug | Inhibited | 6.16E-10 |
| niclosamide | chemical drug | Inhibited | 7.1E-10 |
| retinol acetate | chemical drug | Inhibited | 1.64E-09 |
| drospirenone | chemical drug | Inhibited | 1.88E-09 |
| silicon phthalocyanine | chemical drug | Inhibited | 3.51E-09 |
| L 778123 | chemical drug | Inhibited | 5.87E-09 |
| pexmetinib | chemical drug | Inhibited | 7.47E-09 |
| fulvestrant | chemical drug | Inhibited | 3.85E-08 |
| FCN-159 | chemical drug | Inhibited | 4.12E-08 |
| argatroban | chemical drug | Inhibited | 5.55E-08 |
| midazolam | chemical drug | Inhibited | 8.05E-08 |
| betaxolol | chemical drug | Inhibited | 0.000000101 |
| binimetinib | chemical drug | Inhibited | 0.000000524 |
| epothilone B | chemical drug | Inhibited | 0.00000182 |
| saracatinib | chemical drug | Inhibited | 0.00000183 |
| E 6201 | chemical drug | Inhibited | 0.00000191 |
| melagatran | chemical drug | Inhibited | 0.00000216 |
| L-779450 | chemical drug | Inhibited | 0.00000334 |
| enoxaparin | chemical drug | Inhibited | 0.00000462 |
| CXA-10 | chemical drug | Inhibited | 0.00000614 |
| CH5126766 | chemical drug | Inhibited | 0.00000624 |
| nilutamide | chemical drug | Inhibited | 0.0000073 |
| camostat | chemical drug | Inhibited | 0.00000958 |
| BMS-690514 | chemical drug | Inhibited | 0.0000646 |
| halofuginone | chemical drug | Inhibited | 0.000935 |
| vatalanib | chemical drug | Inhibited | 0.00132 |
| T 0070907 | chemical reagent | | 6.67E-14 |

| | | | |
|---|---|---|---|
| Rp-8-pCPT-cGMPS-triethylamine | chemical reagent | | 4.28E-11 |
| (2-(trimethylammonium)ethyl)methanethiosulfonate | chemical reagent | | 3.74E-10 |
| carbonyl cyanide p-(trifluoromethoxy)phenylhydrazone | chemical reagent | | 0.000000192 |
| 2,2'-dipyridyl disulfide | chemical reagent | | 0.00000202 |
| curcumin derivative C817 | chemical reagent | | 0.0000146 |
| mirin | chemical reagent | | 0.0000687 |
| tributyltin | chemical reagent | | 0.000297 |
| proadifen | chemical reagent | | 0.00148 |
| NSOP00313 | chemical reagent | | 0.00549 |
| L-JNK inhibitor I | chemical reagent | | 0.00549 |
| ACS84 | chemical reagent | | 0.00549 |
| 6-amino-4-(4-phenoxyphenylethylamino)quinazoline | chemical reagent | | 0.00563 |
| polymethyl methacrylate | chemical reagent | | 0.0107 |
| proadifen | chemical reagent | | 0.0434 |
| cobalt chloride | chemical reagent | Activated | 6.53E-16 |
| isatoribine | chemical reagent | Activated | 3.33E-14 |
| 4-((4-(3,4-dichlorophenyl)-1,2,5-thiadiazol-3-yl)oxy)butan-1-ol | chemical reagent | Activated | 4.64E-12 |
| I-BOP | chemical reagent | Activated | 1.26E-11 |
| GnRH-A | chemical reagent | Activated | 2.48E-10 |
| lithocholic acid acetate | chemical reagent | Activated | 0.000000113 |
| 3,5-dihydroxyphenylglycine | chemical reagent | Activated | 0.000000607 |
| ITI-078 | chemical reagent | Activated | 0.00000553 |
| 2-bromoethylamine | chemical reagent | Activated | 0.00114 |
| D-2-amino-5-phosphonovaleric acid | chemical reagent | Inhibited | 2.79E-21 |
| 2-(4-amino-1-isopropyl-1H-pyrazolo[3,4-d]pyrimidin-3-yl)-1H-indol-5-ol | chemical reagent | Inhibited | 6.42E-18 |
| sodium bisulfide | chemical reagent | Inhibited | 1.98E-12 |
| ethylene glycol tetraacetic acid | chemical reagent | Inhibited | 1.12E-11 |
| GW3965 | chemical reagent | Inhibited | 3.24E-10 |
| GGTI-2154 | chemical reagent | Inhibited | 6.14E-09 |
| L-JNK inhibitor I | chemical reagent | Inhibited | 0.0000106 |
| purvalanol A | chemical reagent | Inhibited | 0.0000253 |
| latrunculin B | chemical reagent | Inhibited | 0.00061 |
| DMH1 | chemical reagent | Inhibited | 0.00249 |
| pentachlorophenol | chemical toxicant | | 2.77E-15 |
| 1,10-phenanthroline | chemical toxicant | | 6.32E-11 |
| arotinoid acid | chemical toxicant | | 4.05E-08 |
| alpha-amanitin | chemical toxicant | | 0.00000118 |

| | | | |
|---|---|---|---|
| calicheamicin gamma(1)I | chemical toxicant | | 0.0000435 |
| diquat | chemical toxicant | | 0.000764 |
| phorbol 12,13-didecanoate | chemical toxicant | | 0.000891 |
| Ro 25-6760 | chemical toxicant | | 0.00267 |
| biochanin A | chemical toxicant | | 0.00663 |
| kainic acid | chemical toxicant | Activated | 5.48E-14 |
| methylnitronitrosoguanidine | chemical toxicant | Activated | 4.51E-13 |
| carbon tetrachloride | chemical toxicant | Activated | 7.02E-13 |
| latrunculin A | chemical toxicant | Activated | 1.12E-09 |
| Salmonella enterica serotype abortus equi lipopolysaccharide | chemical toxicant | Activated | 1.02E-08 |
| aroclor 1254 | chemical toxicant | Activated | 0.000000041 |
| benzo(a)pyrene 7,8-dihydrodiol | chemical toxicant | Activated | 0.000000156 |
| 2,3',4,4',5-pentachlorobiphenyl | chemical toxicant | Activated | 0.0000003 |
| Salmonella enterica serotype abortus equi lipopolysaccharide | chemical toxicant | Activated | 0.0000012 |
| dimethylnitrosamine | chemical toxicant | Activated | 0.00000223 |
| zearalenone | chemical toxicant | Activated | 0.0000102 |
| thioacetamide | chemical toxicant | Activated | 0.0000182 |
| dimethylnitrosamine | chemical toxicant | Activated | 0.0000345 |
| carbon tetrachloride | chemical toxicant | Activated | 0.00021 |
| cytochalasin B | chemical toxicant | Inhibited | 1.51E-09 |
| norethynodrel | chemical toxicant | Inhibited | 0.000000441 |
| cypermethrin | chemical toxicant | Inhibited | 0.0000115 |
| VitaminD3-VDR-RXR | complex | | 4.07E-16 |
| T3-TR-RXR | complex | | 1.57E-14 |
| SWI-SNF | complex | | 9.42E-14 |
| PKR dimer | complex | | 2.05E-11 |
| APPBP1/UBA3 Complex | complex | | 1.65E-10 |
| PDGF-DD | complex | | 4.28E-08 |
| SWI-SNF | complex | | 0.000000161 |
| MEF2D-NFAT2-p300 | complex | | 0.000000165 |
| Integrin alpha 6 beta 1 | complex | | 0.00000134 |
| Glycoprotein 1B | complex | | 0.0000118 |
| Laminin (complex) | complex | | 0.000451 |
| IgG2b | complex | | 0.00131 |
| histone deacetylase | complex | | 0.0016 |
| PDGF-Rβ dimer | complex | | 0.00184 |
| Bcl10-Card10-Malt1 | complex | | 0.00326 |
| PDGF-Rβ dimer | complex | | 0.0434 |
| Laminin (complex) | complex | Activated | 1.53E-17 |
| Ige | complex | Activated | 1.71E-14 |
| Nodal receptor | complex | Activated | 4.32E-14 |
| IGg-Rheumatoid factor | complex | Activated | 5.76E-13 |
| NMDA Receptor | complex | Activated | 1.48E-12 |
| CD247 dimer | complex | Activated | 2.95E-10 |
| CAK | complex | Activated | 3.2E-09 |
| EIF3 | complex | Activated | 3.92E-09 |

| | | | |
|---|---|---|---|
| Esr1-Esr1-estrogen-estrogen | complex | Activated | 4.19E-08 |
| NFkB1-CRel | complex | Activated | 0.00000789 |
| cyclin h/cdk7 | complex | Activated | 0.0000426 |
| EIF4F | complex | Activated | 0.0000526 |
| APC (complex) | complex | Activated | 0.000149 |
| histone deacetylase | complex | Inhibited | 1.7E-09 |
| I kappa b kinase | complex | Inhibited | 0.0000087 |
| TIMP1 | cytokine | | 0.00267 |
| CXCL11 | cytokine | | 0.00549 |
| CSF2 | cytokine | Activated | 2.77E-18 |
| CXCL12 | cytokine | Activated | 5.07E-18 |
| CSF1 | cytokine | Activated | 7.14E-18 |
| CSF2 | cytokine | Activated | 1.92E-17 |
| IL1B | cytokine | Activated | 1.08E-14 |
| IL20 | cytokine | Activated | 4.19E-14 |
| WNT3A | cytokine | Activated | 1.31E-13 |
| PF4 | cytokine | Activated | 1.54E-09 |
| CXCL12 | cytokine | Activated | 0.0000305 |
| CYP27A1 | enzyme | | 2.51E-15 |
| FBXW11 | enzyme | | 2.6E-15 |
| CYP19A1 | enzyme | | 2.98E-15 |
| RECQL5 | enzyme | | 3.36E-13 |
| RNF19A | enzyme | | 2.08E-12 |
| UHRF2 | enzyme | | 4.64E-12 |
| CUL1 | enzyme | | 4.85E-12 |
| UBE4B | enzyme | | 1.24E-11 |
| OTUB1 | enzyme | | 1.66E-11 |
| RCHY1 | enzyme | | 5.35E-11 |
| RALA | enzyme | | 7.44E-11 |
| POR | enzyme | | 7.63E-11 |
| RALB | enzyme | | 1.03E-10 |
| MLH1 | enzyme | | 1.07E-10 |
| BIRC2 | enzyme | | 2.13E-10 |
| RAD50 | enzyme | | 1.21E-09 |
| RGS6 | enzyme | | 1.52E-09 |
| HPRT1 | enzyme | | 5.18E-08 |
| CAB39 | enzyme | | 0.000000206 |
| MGMT | enzyme | | 0.00000196 |
| ARIH2 | enzyme | | 0.00000689 |
| IHH | enzyme | | 0.000155 |
| AGA | enzyme | | 0.000764 |
| TREX1 | enzyme | | 0.00107 |
| WWOX | enzyme | | 0.00137 |
| IHH | enzyme | | 0.00151 |
| CYP1B1 | enzyme | | 0.00244 |
| PLCD1 | enzyme | | 0.00388 |
| GSR | enzyme | | 0.0042 |
| HERC5 | enzyme | | 0.00549 |

| | | | |
|---|---|---|---|
| PARP14 | enzyme | | 0.00563 |
| HAS2 | enzyme | | 0.00563 |
| NDUFA13 | enzyme | | 0.00778 |
| GNA11 | enzyme | | 0.00778 |
| MBD4 | enzyme | | 0.0107 |
| CHI3L1 | enzyme | | 0.0107 |
| ARF1 | enzyme | | 0.0107 |
| PDE4B | enzyme | | 0.0251 |
| SI | enzyme | | 0.0434 |
| KDM4B | enzyme | Activated | 1.46E-14 |
| SMYD3 | enzyme | Activated | 1.56E-14 |
| RAP2A | enzyme | Activated | 3.45E-14 |
| P4HB | enzyme | Activated | 1.02E-13 |
| HMGA2 | enzyme | Activated | 1.62E-13 |
| PPME1 | enzyme | Activated | 5.6E-13 |
| KDM1A | enzyme | Activated | 5.94E-13 |
| PELI1 | enzyme | Activated | 2.67E-10 |
| GNAS | enzyme | Activated | 8.96E-10 |
| SETD7 | enzyme | Activated | 5.76E-09 |
| TERT | enzyme | Activated | 1.44E-08 |
| PLA2G5 | enzyme | Activated | 0.000000119 |
| KDM8 | enzyme | Activated | 0.00000207 |
| PTGES3 | enzyme | Activated | 0.00000216 |
| RAC3 | enzyme | Activated | 0.00000364 |
| PPID | enzyme | Activated | 0.00000454 |
| AKR1C3 | enzyme | Activated | 0.0000309 |
| HINT1 | enzyme | Inhibited | 4.15E-16 |
| PPIF | enzyme | Inhibited | 6.65E-16 |
| COMT | enzyme | Inhibited | 2.08E-14 |
| PLCL2 | enzyme | Inhibited | 2.41E-12 |
| GNG7 | enzyme | Inhibited | 7.17E-12 |
| FBXO4 | enzyme | Inhibited | 0.000000058 |
| CHFR | enzyme | Inhibited | 0.00000166 |
| DNMT3A | enzyme | Inhibited | 0.0000026 |
| DNMT3A | enzyme | Inhibited | 0.0000059 |
| DNMT3B | enzyme | Inhibited | 0.00033 |
| DNASE2 | enzyme | Inhibited | 0.00267 |
| KDM3B | enzyme | Inhibited | 0.00851 |
| RUNX1-RUNX1T1 | fusion gene/product | | 0.00328 |
| DRD4 | G-protein coupled receptor | | 1.27E-12 |
| HTR6 | G-protein coupled receptor | | 5.12E-11 |
| FFAR3 | G-protein coupled receptor | | 3.19E-10 |
| NMBR | G-protein coupled receptor | | 0.0000046 |
| AVPR2 | G-protein coupled receptor | | 0.0000103 |
| GPR68 | G-protein coupled receptor | | 0.0173 |
| FZD7 | G-protein coupled receptor | | 0.0173 |
| ADGRE5 | G-protein coupled receptor | Activated | 3.15E-13 |
| CXCR3 | G-protein coupled receptor | Activated | 3.2E-12 |

| | | | |
|---|---|---|---|
| P2RY4 | G-protein coupled receptor | Activated | 4.27E-11 |
| F2R | G-protein coupled receptor | Activated | 0.00000357 |
| CASR | G-protein coupled receptor | Activated | 0.0000714 |
| PTGER2 | G-protein coupled receptor | Activated | 0.0012 |
| ADGRG3 | G-protein coupled receptor | Inhibited | 2.46E-13 |
| ACKR2 | G-protein coupled receptor | Inhibited | 0.00203 |
| PI4K | group | | 1.47E-17 |
| MEF2 | group | | 8.62E-15 |
| Lysosomal Protease | group | | 5.91E-14 |
| Alpha Actinin | group | | 6.63E-14 |
| Gata | group | | 6.93E-14 |
| Hdac | group | | 1.93E-13 |
| Focal adhesion kinase | group | | 2E-13 |
| Cdc2 | group | | 1.61E-11 |
| B56 | group | | 6.99E-11 |
| ALDH | group | | 3.66E-10 |
| Pkar2 | group | | 8.68E-09 |
| Calcineurin A | group | | 1.71E-08 |
| MEF2 | group | | 0.000000204 |
| DNA-methyltransferase | group | | 0.000000234 |
| DUB | group | | 0.000018 |
| ADRB | group | | 0.000037 |
| tryptase | group | | 0.000111 |
| G-Actin | group | | 0.00233 |
| Ikk (family) | group | | 0.00549 |
| Diap | group | | 0.00549 |
| Nfat (family) | group | Activated | 8.55E-19 |
| Vegf | group | Activated | 7.69E-18 |
| E2f | group | Activated | 7.71E-18 |
| estrogen receptor | group | Activated | 4.22E-15 |
| PDZGEF | group | Activated | 3.36E-13 |
| Raf | group | Activated | 8.64E-13 |
| DRD1/5 | group | Activated | 8.04E-12 |
| receptor protein tyrosine kinase | group | Activated | 1.3E-11 |
| IL-1R | group | Activated | 2.02E-10 |
| Par | group | Activated | 9.4E-10 |
| SYK/ZAP | group | Activated | 1.09E-09 |
| Vegf | group | Activated | 1.85E-09 |
| Gα12/13 | group | Activated | 0.000000398 |
| MIR99A-LET7C-MIR125B2 | group | Inhibited | 2.87E-16 |
| MIR100-LET7A2-MIR125B1 | group | Inhibited | 2.87E-16 |
| MIRLET7 | group | Inhibited | 3.94E-16 |
| SFRP | group | Inhibited | 7.93E-15 |
| Lefty | group | Inhibited | 1.48E-13 |
| Npm | group | Inhibited | 2.13E-13 |
| Endophilin | group | Inhibited | 1.21E-12 |
| Rb | group | Inhibited | 2.08E-12 |
| MIRLET7 | group | Inhibited | 7.37E-11 |

| | | | |
|---|---|---|---|
| MIR99A-LET7C-MIR125B2 | group | Inhibited | 1.49E-09 |
| MIR100-LET7A2-MIR125B1 | group | Inhibited | 1.49E-09 |
| Gamma tubulin | group | Inhibited | 1.64E-08 |
| Alpha 1 antitrypsin | group | Inhibited | 0.000000892 |
| FGF3 | growth factor | | 1.13E-11 |
| GDF9 | growth factor | | 0.000451 |
| BMP4 | growth factor | Activated | 1.35E-15 |
| NDP | growth factor | Activated | 1.46E-14 |
| HDGF | growth factor | Activated | 3.6E-13 |
| FGF4 | growth factor | Activated | 1.38E-09 |
| HGF | growth factor | Activated | 4.32E-08 |
| FGF6 | growth factor | Activated | 0.000000138 |
| FGF3 | growth factor | Activated | 0.000111 |
| JAG1 | growth factor | Activated | 0.000419 |
| RASA3 | ion channel | | 7.54E-12 |
| Cacnb1 | ion channel | | 0.00549 |
| CACNA2D1 | ion channel | | 0.00549 |
| DYRK1A | kinase | | 6.81E-19 |
| MAK | kinase | | 2.03E-18 |
| DYRK1A | kinase | | 1.43E-16 |
| STK38L | kinase | | 4.34E-15 |
| MYLK2 | kinase | | 4.49E-13 |
| PAK6 | kinase | | 2.32E-11 |
| CAMK2G | kinase | | 5.4E-11 |
| AURKB | kinase | | 6.31E-10 |
| STRADA | kinase | | 3.53E-08 |
| Brd4 | kinase | | 0.000000324 |
| CDC7 | kinase | | 0.000000845 |
| STK11 | kinase | | 0.00000365 |
| MAPK1 | kinase | | 0.0000204 |
| CPNE3 | kinase | | 0.0000271 |
| ATM | kinase | | 0.0000479 |
| CHEK1 | kinase | | 0.00011 |
| MAPKAPK2 | kinase | | 0.000112 |
| AURKB | kinase | | 0.000151 |
| TBK1 | kinase | | 0.00037 |
| PLK2 | kinase | | 0.000384 |
| CDK12 | kinase | | 0.000764 |
| NTRK2 | kinase | | 0.0013 |
| FGFR2 | kinase | | 0.00156 |
| PTK6 | kinase | | 0.0107 |
| TRPM7 | kinase | | 0.0434 |
| HIPK3 | kinase | | 0.0434 |
| PKM | kinase | Activated | 3.18E-21 |
| ERBB2 | kinase | Activated | 2.01E-17 |
| PRKCZ | kinase | Activated | 6.14E-16 |
| MAP3K8 | kinase | Activated | 1.02E-14 |
| STK26 | kinase | Activated | 1.37E-14 |

| | | | |
|---|---|---|---|
| MAP3K3 | kinase | Activated | 4.87E-14 |
| JAK1 | kinase | Activated | 2.37E-13 |
| MAPK11 | kinase | Activated | 2.78E-13 |
| MAP3K2 | kinase | Activated | 9.83E-13 |
| CAMK2A | kinase | Activated | 5.13E-12 |
| PRKCI | kinase | Activated | 1.14E-10 |
| MOS | kinase | Activated | 3.82E-10 |
| CKS1B | kinase | Activated | 2.12E-08 |
| TAOK2 | kinase | Activated | 5.62E-08 |
| BRD4 | kinase | Activated | 0.000000216 |
| Brd4 | kinase | Activated | 0.00000301 |
| MAP2K1 | kinase | Activated | 0.00000544 |
| CCNK | kinase | Activated | 0.0000396 |
| CRKL | kinase | Activated | 0.00328 |
| MARK2 | kinase | Inhibited | 2.21E-15 |
| CDK8 | kinase | Inhibited | 9.39E-10 |
| ITPKB | kinase | Inhibited | 5.13E-09 |
| CDKN1A | kinase | Inhibited | 0.000000256 |
| CAMK2N1 | kinase | Inhibited | 0.000000263 |
| LATS1 | kinase | Inhibited | 0.00000207 |
| DYRK1A | kinase | Inhibited | 0.0000273 |
| PLK1 | kinase | Inhibited | 0.00534 |
| RARA | ligand-dependent nuclear receptor | | 0.0000459 |
| ESR1 | ligand-dependent nuclear receptor | Activated | 3.15E-14 |
| RORC | ligand-dependent nuclear receptor | Activated | 5.24E-12 |
| AR | ligand-dependent nuclear receptor | Activated | 0.0000376 |
| NR2E1 | ligand-dependent nuclear receptor | Inhibited | 0.0000276 |
| miR-515-5p (and other miRNAs w/seed UCUCCAA) | mature microRNA | | 0.0107 |
| miR-382-5p (miRNAs w/seed AAGUUGU) | mature microRNA | | 0.0173 |
| miR-105-5p (and other miRNAs w/seed CAAAUGC) | mature microRNA | | 0.0434 |
| miR-148a-3p (and other miRNAs w/seed CAGUGCA) | mature microRNA | Inhibited | 0.00022 |
| miR-34a-5p (and other miRNAs w/seed GGCAGUG) | mature microRNA | Inhibited | 0.00195 |
| miR-24-3p (and other miRNAs w/seed GGCUCAG) | mature microRNA | Inhibited | 0.00653 |
| mir-192 | microRNA | | 0.000000242 |
| mir-19 | microRNA | | 0.000188 |
| mir-503 | microRNA | | 0.00392 |
| mir-340 | microRNA | | 0.0434 |
| let-7 | microRNA | Inhibited | 1.95E-14 |
| mir-17 | microRNA | Inhibited | 2.57E-11 |
| let-7 | microRNA | Inhibited | 1.99E-09 |
| mir-21 | microRNA | Inhibited | 0.00000454 |
| ARRDC3 | other | | 6.32E-14 |

| | | |
|---|---|---|
| TG | other | 3.4E-13 |
| RALGAPB | other | 6.62E-13 |
| INCENP | other | 2.67E-11 |
| TNRC6A | other | 3.89E-11 |
| DDIT4 | other | 5.35E-11 |
| ELAVL1 | other | 5.19E-10 |
| YWHAQ | other | 4.71E-09 |
| CUL4A | other | 3.74E-08 |
| FRAT1 | other | 4.16E-08 |
| FADD | other | 5.72E-08 |
| HLA-A | other | 0.000000103 |
| SELPLG | other | 0.000000107 |
| BNIP3L | other | 0.000000165 |
| USO1 | other | 0.000000221 |
| NLRP3 | other | 0.00000801 |
| CD59 | other | 0.0000109 |
| SYF2 | other | 0.0000115 |
| GP9 | other | 0.0000133 |
| SV2C | other | 0.0000175 |
| CDC6 | other | 0.0000196 |
| PVR | other | 0.0000361 |
| PALB2 | other | 0.0000479 |
| VWC2 | other | 0.0000499 |
| CBX7 | other | 0.0000819 |
| SDC3 | other | 0.000147 |
| HOTAIR | other | 0.000632 |
| MKI67 | other | 0.000764 |
| VGLL3 | other | 0.00137 |
| SOCS6 | other | 0.00174 |
| DCTN4 | other | 0.00188 |
| LAMA1 | other | 0.00188 |
| DLL4 | other | 0.00203 |
| TNFAIP6 | other | 0.00217 |
| CLDN7 | other | 0.00249 |
| SERPINH1 | other | 0.0025 |
| Ptgs2os2 | other | 0.0025 |
| ATG13 | other | 0.00423 |
| FBXO42 | other | 0.00534 |
| SFN | other | 0.00556 |
| CDKN1C | other | 0.00563 |
| VGLL3 | other | 0.00663 |
| CDH2 | other | 0.00778 |
| Nppb | other | 0.0107 |
| LSP1 | other | 0.0107 |
| SCIN | other | 0.0107 |
| RB1CC1 | other | 0.0134 |
| TRIM38 | other | 0.0168 |
| TRIM58 | other | 0.0173 |

| | | | |
|---|---|---|---|
| C4A/C4B | other | | 0.0251 |
| KRT1 | other | | 0.0434 |
| FSIP1 | other | | 0.0434 |
| LINC02599 | other | | 0.0434 |
| OPCML | other | | 0.0434 |
| USP50 | other | | 0.0434 |
| LINC00261 | other | | 0.0434 |
| CEP78 | other | | 0.0434 |
| SYN2 | other | | 0.0434 |
| NLRP9 | other | | 0.0434 |
| KRT3 | other | | 0.0434 |
| CD151 | other | Activated | 6.35E-22 |
| CALCA | other | Activated | 4.15E-20 |
| POMC | other | Activated | 1.21E-17 |
| APBB1IP | other | Activated | 1.39E-13 |
| KLHL42 | other | Activated | 4.53E-13 |
| RADIL | other | Activated | 5.2E-13 |
| EFHD2 | other | Activated | 4.79E-12 |
| BLNK | other | Activated | 7.82E-12 |
| IL34 | other | Activated | 1.3E-11 |
| RABL6 | other | Activated | 2.74E-11 |
| BAK1 | other | Activated | 3.53E-11 |
| BEX2 | other | Activated | 4.81E-11 |
| HOTAIR | other | Activated | 5.09E-11 |
| S100A8 | other | Activated | 1.48E-10 |
| S100A9 | other | Activated | 5.18E-10 |
| SORBS3 | other | Activated | 7.1E-10 |
| CRK | other | Activated | 7.49E-10 |
| MBL2 | other | Activated | 1.05E-09 |
| EP400 | other | Activated | 1.09E-09 |
| IGFBP6 | other | Activated | 1.81E-09 |
| RETNLB | other | Activated | 2.75E-09 |
| RANBP9 | other | Activated | 7.58E-09 |
| LAMTOR3 | other | Activated | 3.39E-08 |
| FBLN5 | other | Activated | 0.000000184 |
| ARHGEF11 | other | Activated | 0.000000224 |
| CYTH2 | other | Activated | 0.00000047 |
| INIP | other | Activated | 0.000000593 |
| INTS3 | other | Activated | 0.000000593 |
| SASH1 | other | Activated | 0.000000788 |
| FGD1 | other | Activated | 0.00000266 |
| BOP1 | other | Activated | 0.00000379 |
| RNF123 | other | Activated | 0.00000533 |
| CLEC12A | other | Activated | 0.0000181 |
| BRAT1 | other | Activated | 0.0000341 |
| ACTB | other | Activated | 0.0000356 |
| SPTAN1 | other | Activated | 0.000043 |
| PTH | other | Activated | 0.0000447 |

| | | | |
|---|---|---|---|
| SASH1 | other | Activated | 0.00023 |
| Bvht | other | Activated | 0.000333 |
| ARHGAP21 | other | Activated | 0.000459 |
| STING1 | other | Activated | 0.000556 |
| SAMSN1 | other | Activated | 0.000561 |
| DSCAM | other | Activated | 0.00251 |
| CDH11 | other | Activated | 0.00653 |
| EWSR1 | other | Activated | 0.0101 |
| TRIM65 | other | Inhibited | 3.32E-18 |
| LEMD3 | other | Inhibited | 1.59E-16 |
| SAFB | other | Inhibited | 8.26E-16 |
| CBX7 | other | Inhibited | 9.66E-16 |
| SAFB2 | other | Inhibited | 3.86E-15 |
| IFRD1 | other | Inhibited | 9.9E-15 |
| KRIT1 | other | Inhibited | 1.2E-14 |
| SPARC | other | Inhibited | 1.24E-14 |
| TINCR | other | Inhibited | 1.28E-14 |
| CGN | other | Inhibited | 2.8E-13 |
| SPINT2 | other | Inhibited | 2.98E-13 |
| SERPING1 | other | Inhibited | 1.71E-12 |
| AMOT | other | Inhibited | 2.98E-12 |
| TRIM65 | other | Inhibited | 4.54E-10 |
| SERPIND1 | other | Inhibited | 6.96E-10 |
| CCNC | other | Inhibited | 2.61E-09 |
| PINX1 | other | Inhibited | 9.92E-09 |
| SPINT1 | other | Inhibited | 1.26E-08 |
| Cdc2b | other | Inhibited | 1.37E-08 |
| KLHL40 | other | Inhibited | 1.78E-08 |
| SPRY4 | other | Inhibited | 1.87E-08 |
| Irgm1 | other | Inhibited | 0.000000236 |
| MMS22L | other | Inhibited | 0.000000593 |
| FRMD6 | other | Inhibited | 0.000000736 |
| Irgm1 | other | Inhibited | 0.000002 |
| SPARC | other | Inhibited | 0.00000456 |
| SFPQ | other | Inhibited | 0.00000637 |
| SERPINA5 | other | Inhibited | 0.00000749 |
| TFPI2 | other | Inhibited | 0.00000984 |
| RASSF6 | other | Inhibited | 0.00002 |
| SPINK5 | other | Inhibited | 0.000289 |
| 1810019D21Rik | other | Inhibited | 0.000941 |
| PCBP2 | other | Inhibited | 0.000947 |
| CDON | other | Inhibited | 0.00132 |
| RCE1 | peptidase | | 6.4E-13 |
| F10 | peptidase | | 5.9E-12 |
| CAST | peptidase | | 1.63E-10 |
| EPS8 | peptidase | | 4.84E-09 |
| USP24 | peptidase | | 0.000000135 |
| USP12 | peptidase | | 0.000000302 |

| | | | |
|---|---|---|---|
| CAPN3 | peptidase | | 0.00328 |
| USP1 | peptidase | | 0.00549 |
| ZFYVE9 | peptidase | | 0.0107 |
| USP6 | peptidase | Activated | 1.45E-13 |
| CTSC | peptidase | Activated | 6.87E-12 |
| GZMB | peptidase | Activated | 6.79E-11 |
| ST14 | peptidase | Activated | 4.96E-10 |
| F11 | peptidase | Activated | 1.65E-09 |
| UCHL5 | peptidase | Activated | 2.89E-09 |
| USP5 | peptidase | Activated | 0.000000513 |
| F2 | peptidase | Activated | 0.0000121 |
| ADAMTS18 | peptidase | Activated | 0.0000396 |
| PLG | peptidase | Activated | 0.00386 |
| CCAR2 | peptidase | Inhibited | 6.24E-08 |
| PSMB11 | peptidase | Inhibited | 7.29E-08 |
| USP38 | peptidase | Inhibited | 0.000000644 |
| CPN1 | peptidase | Inhibited | 0.000422 |
| PPM1A | phosphatase | | 1.69E-11 |
| CDC25B | phosphatase | | 1.08E-10 |
| CDC14A | phosphatase | | 1.05E-09 |
| CDC14B | phosphatase | | 1.09E-09 |
| PPEF2 | phosphatase | | 0.00000238 |
| PPP3R1 | phosphatase | | 0.00152 |
| TNS3 | phosphatase | Activated | 9.72E-13 |
| PTPN22 | phosphatase | Inhibited | 7.96E-18 |
| PTPN9 | phosphatase | Inhibited | 1.58E-17 |
| SOX2 | transcription regulator | | 5.31E-18 |
| GATA1 | transcription regulator | | 1.2E-14 |
| BCL6 | transcription regulator | | 4.71E-14 |
| ID2 | transcription regulator | | 8.31E-13 |
| MEF2C | transcription regulator | | 1.03E-12 |
| NFKB1 | transcription regulator | | 1.03E-12 |
| ZBTB16 | transcription regulator | | 1.22E-12 |
| HMGA1 | transcription regulator | | 2.98E-12 |
| SOX2 | transcription regulator | | 3.35E-12 |
| SMARCA4 | transcription regulator | | 3.56E-12 |
| CALR | transcription regulator | | 4.18E-12 |
| KLF8 | transcription regulator | | 6.29E-12 |
| H2AX | transcription regulator | | 1.04E-11 |
| TAF1 | transcription regulator | | 2.19E-11 |
| FLI1 | transcription regulator | | 2.54E-11 |
| HEXIM1 | transcription regulator | | 6.92E-11 |
| SMARCA2 | transcription regulator | | 3.83E-10 |
| NKX2-5 | transcription regulator | | 5.11E-10 |
| SPI1 | transcription regulator | | 2.1E-09 |
| CBFA2T3 | transcription regulator | | 3.04E-09 |
| GATA1 | transcription regulator | | 3.4E-09 |
| TCF7L2 | transcription regulator | | 8.06E-09 |

| | | | |
|---|---|---|---|
| TMF1 | transcription regulator | | 1.44E-08 |
| GATA1 | transcription regulator | | 5.59E-08 |
| IKZF1 | transcription regulator | | 0.000000102 |
| MEF2D | transcription regulator | | 0.000000319 |
| GATA2 | transcription regulator | | 0.00000183 |
| SMARCB1 | transcription regulator | | 0.00000385 |
| SOX2 | transcription regulator | | 0.0000114 |
| TONSL | transcription regulator | | 0.0000134 |
| VDR | transcription regulator | | 0.0000163 |
| KDM5B | transcription regulator | | 0.0000223 |
| WWTR1 | transcription regulator | | 0.0000904 |
| NOTCH4 | transcription regulator | | 0.000118 |
| HDAC2 | transcription regulator | | 0.000208 |
| UXT | transcription regulator | | 0.000369 |
| SMARCA2 | transcription regulator | | 0.000459 |
| CBFB | transcription regulator | | 0.000591 |
| GABPA | transcription regulator | | 0.000829 |
| ZNF217 | transcription regulator | | 0.000855 |
| FOXN4 | transcription regulator | | 0.000855 |
| FLI1 | transcription regulator | | 0.000987 |
| MED16 | transcription regulator | | 0.00132 |
| SKI | transcription regulator | | 0.00137 |
| E2F7 | transcription regulator | | 0.00148 |
| UBP1 | transcription regulator | | 0.00188 |
| NKX2-5 | transcription regulator | | 0.00191 |
| TOB1 | transcription regulator | | 0.00442 |
| HAND2 | transcription regulator | | 0.00442 |
| ZHX2 | transcription regulator | | 0.00549 |
| LPXN | transcription regulator | | 0.00549 |
| HMG20B | transcription regulator | | 0.00549 |
| RBBP7 | transcription regulator | | 0.00549 |
| NEO1 | transcription regulator | | 0.00563 |
| ARID2 | transcription regulator | | 0.0107 |
| DR1 | transcription regulator | | 0.0251 |
| DEPDC1 | transcription regulator | | 0.0434 |
| TFCP2L1 | transcription regulator | | 0.0434 |
| TBX18 | transcription regulator | | 0.0434 |
| HDAC2 | transcription regulator | Activated | 2.41E-15 |
| TP63 | transcription regulator | Activated | 5.54E-15 |
| CTNNB1 | transcription regulator | Activated | 2.23E-14 |
| MITF | transcription regulator | Activated | 3.11E-14 |
| SMAD2 | transcription regulator | Activated | 6.76E-13 |
| CBX4 | transcription regulator | Activated | 1.51E-12 |
| TBX2 | transcription regulator | Activated | 4.49E-12 |
| SP3 | transcription regulator | Activated | 7.45E-11 |
| ZNF217 | transcription regulator | Activated | 1.48E-10 |
| PURA | transcription regulator | Activated | 1.59E-09 |
| SERTAD1 | transcription regulator | Activated | 2.16E-09 |

| | | | |
|---|---|---|---|
| ETV1 | transcription regulator | Activated | 5.04E-09 |
| NFIC | transcription regulator | Activated | 9.58E-09 |
| TFDP2 | transcription regulator | Activated | 1.64E-08 |
| E2F1 | transcription regulator | Activated | 0.00000004 |
| UHRF1 | transcription regulator | Activated | 0.000000133 |
| CCND1 | transcription regulator | Activated | 0.000000172 |
| SMARCE1 | transcription regulator | Activated | 0.000000223 |
| TP63 | transcription regulator | Activated | 0.000000892 |
| CCNE1 | transcription regulator | Activated | 0.000000969 |
| YAP1 | transcription regulator | Activated | 0.00000228 |
| E2F2 | transcription regulator | Activated | 0.00000584 |
| EHF | transcription regulator | Activated | 0.0000103 |
| E2F3 | transcription regulator | Activated | 0.0000104 |
| SMARCA4 | transcription regulator | Activated | 0.0000496 |
| NOTCH3 | transcription regulator | Activated | 0.000227 |
| FOXM1 | transcription regulator | Activated | 0.000377 |
| KDM3A | transcription regulator | Activated | 0.000388 |
| TAL1 | transcription regulator | Activated | 0.000485 |
| E2F2 | transcription regulator | Activated | 0.000737 |
| ZNF281 | transcription regulator | Activated | 0.00109 |
| TCF7L2 | transcription regulator | Activated | 0.00144 |
| REL | transcription regulator | Activated | 0.00155 |
| HOXA9 | transcription regulator | Activated | 0.00212 |
| WBP2 | transcription regulator | Activated | 0.00224 |
| ETS1 | transcription regulator | Inhibited | 1.27E-23 |
| MEOX1 | transcription regulator | Inhibited | 1.94E-15 |
| CDKN2A | transcription regulator | Inhibited | 3.06E-15 |
| HDAC7 | transcription regulator | Inhibited | 3.07E-15 |
| ZFHX3 | transcription regulator | Inhibited | 3.08E-15 |
| SOX11 | transcription regulator | Inhibited | 4.21E-15 |
| HDAC1 | transcription regulator | Inhibited | 1.28E-13 |
| MEOX1 | transcription regulator | Inhibited | 2.13E-13 |
| ZFP36 | transcription regulator | Inhibited | 3.17E-13 |
| CDKN2A | transcription regulator | Inhibited | 3.28E-13 |
| E2F6 | transcription regulator | Inhibited | 5.85E-12 |
| NUPR1 | transcription regulator | Inhibited | 1.99E-10 |
| SPDEF | transcription regulator | Inhibited | 5.07E-09 |
| NFX1 | transcription regulator | Inhibited | 7.26E-09 |
| RBL1 | transcription regulator | Inhibited | 0.000000064 |
| TRERF1 | transcription regulator | Inhibited | 0.000000122 |
| DNMT3L | transcription regulator | Inhibited | 0.00000017 |
| ZIC2 | transcription regulator | Inhibited | 0.000000229 |
| PSIP1 | transcription regulator | Inhibited | 0.00000058 |
| NUPR1 | transcription regulator | Inhibited | 0.000000731 |
| HOXA4 | transcription regulator | Inhibited | 0.00000167 |
| HEYL | transcription regulator | Inhibited | 0.0000171 |
| FOXH1 | transcription regulator | Inhibited | 0.0000185 |
| RB1 | transcription regulator | Inhibited | 0.0000894 |

| | | | |
|---|---|---|---|
| FOXP3 | transcription regulator | Inhibited | 0.000295 |
| PTTG1 | transcription regulator | Inhibited | 0.000295 |
| HDAC11 | transcription regulator | Inhibited | 0.000386 |
| GLIS2 | transcription regulator | Inhibited | 0.000815 |
| ZFP36 | transcription regulator | Inhibited | 0.00118 |
| NOSTRIN | transcription regulator | Inhibited | 0.00227 |
| Bc1 | translation regulator | | 0.0107 |
| EIF4G1 | translation regulator | Activated | 0.00217 |
| TNFRSF1A | transmembrane receptor | | 8.42E-12 |
| FAS | transmembrane receptor | | 0.000112 |
| PLAUR | transmembrane receptor | | 0.000787 |
| FCGR1A | transmembrane receptor | | 0.0025 |
| ENG | transmembrane receptor | | 0.00267 |
| GP1BA | transmembrane receptor | Activated | 4.49E-15 |
| IL21R | transmembrane receptor | Activated | 3.99E-14 |
| ITGAV | transmembrane receptor | Activated | 5.41E-14 |
| ITGB3 | transmembrane receptor | Activated | 2.13E-13 |
| KLRF1 | transmembrane receptor | Activated | 3.79E-13 |
| CLEC1B | transmembrane receptor | Activated | 5.51E-13 |
| ITGB1 | transmembrane receptor | Activated | 1.13E-12 |
| IL17RA | transmembrane receptor | Activated | 6.02E-10 |
| ITGA2 | transmembrane receptor | Activated | 0.00000001 |
| TREM1 | transmembrane receptor | Activated | 1.14E-08 |
| FCGR1A | transmembrane receptor | Activated | 1.46E-08 |
| IL1R1 | transmembrane receptor | Activated | 0.00114 |
| SNX9 | transporter | | 4.42E-12 |
| RACGAP1 | transporter | | 0.00000612 |
| SYN1 | transporter | | 0.0434 |
| GJA3 | transporter | | 0.0434 |
| SNX5 | transporter | Inhibited | 4.84E-13 |
| AMBP | transporter | Inhibited | 3.04E-09 |

**Supplementary Table 2. Etoposide-induced differences in master regulators in WT cells**

| **Master Regulator** | **Molecule Type** | **Predicted** | **p-value of** |
|---|---|---|---|
| Master Regulator | Molecule Type | Predicted | p-value of |
| denileukin diftitox | biologic drug | | 1.68E-14 |
| cdc25C phosphatase (211-221) | biologic drug | | 1.34E-08 |
| vasopressins | biologic drug | Activated | 1.64E-14 |
| desmopressin | biologic drug | Activated | 3.39E-10 |
| desmopressin | biologic drug | Activated | 0.000273 |
| interferon alfa-n1 | biologic drug | Activated | 0.000387 |
| PEG-interferon alfa-2a | biologic drug | Activated | 0.000387 |
| recombinant interferon | biologic drug | Activated | 0.000387 |
| l-asparaginase | biologic drug | Inhibited | 7.43E-27 |
| bivalirudin | biologic drug | Inhibited | 7.4E-08 |
| heparin | chemical - endogenous mammalian | | 6.69E-15 |
| 9,13-di-cis-retinoic acid | chemical - endogenous mammalian | | 9.12E-14 |
| 4-hydroxy-all-trans-retinoic acid | chemical - endogenous mammalian | | 2.67E-13 |
| beta-apo-14'-carotenal | chemical - endogenous mammalian | | 2.67E-13 |
| all-trans-4-Oxo-retinoic acid | chemical - endogenous mammalian | | 2.67E-13 |
| 18-hydroxy-retinoic acid | chemical - endogenous mammalian | | 2.67E-13 |
| epinephrine | chemical - endogenous mammalian | | 2.93E-13 |
| formic acid | chemical - endogenous mammalian | | 2.78E-11 |
| propionic acid | chemical - endogenous mammalian | | 3.02E-11 |
| pentanoic acid | chemical - endogenous mammalian | | 3.02E-11 |
| ergocalciferol | chemical - endogenous mammalian | | 1.08E-07 |
| calcifediol | chemical - endogenous mammalian | | 3.74E-06 |
| cardiolipin | chemical - endogenous mammalian | | 6.08E-06 |
| taurodeoxycholic acid | chemical - endogenous mammalian | | 1.09E-05 |
| L-carnitine | chemical - endogenous mammalian | | 0.000147 |
| 10E,12Z-octadecadienoic acid | chemical - endogenous mammalian | | 0.00019 |
| 8-epi-prostaglandin F2alpha | chemical - endogenous mammalian | | 0.000764 |
| 19,20-epoxydocosapentaenoic acid | chemical - endogenous mammalian | | 0.00148 |
| 8,9-epoxyeicosatrienoic acid | chemical - endogenous mammalian | | 0.00563 |
| trigonelline | chemical - endogenous mammalian | | 0.0251 |
| SAICAR | chemical - endogenous mammalian | Activated | 7.83E-19 |
| estrone sulfate | chemical - endogenous mammalian | Activated | 1.93E-14 |
| 4-hydroxy-N-desmethyltamoxifen | chemical - endogenous mammalian | Activated | 2.29E-14 |
| UDP-D-glucose | chemical - endogenous mammalian | Activated | 7.04E-14 |
| pregnenolone sulfate | chemical - endogenous mammalian | Activated | 6.56E-13 |
| indican | chemical - endogenous mammalian | Activated | 1.22E-12 |
| acetic acid | chemical - endogenous mammalian | Activated | 3.5E-12 |
| adenine | chemical - endogenous mammalian | Activated | 7.29E-12 |
| uric acid | chemical - endogenous mammalian | Activated | 1.38E-11 |
| acetylcholine | chemical - endogenous mammalian | Activated | 3.82E-10 |
| bombesin | chemical - endogenous mammalian | Activated | 4.43E-08 |
| androstenedione | chemical - endogenous mammalian | Activated | 2.13E-06 |

| | | | |
|---|---|---|---|
| 17-hydroxyprogesterone | chemical - endogenous mammalian | Activated | 1.56E-05 |
| lactosylceramide | chemical - endogenous mammalian | Activated | 0.000601 |
| phospholipid | chemical - endogenous mammalian | Activated | 0.000912 |
| lysophosphatidylinositol | chemical - endogenous mammalian | Activated | 0.000987 |
| 2'3'-cyclic guanosine monophosphate-adenosine monophosphate | chemical - endogenous mammalian | Activated | 0.00184 |
| glycosaminoglycan | chemical - endogenous mammalian | Inhibited | 1.43E-14 |
| trehalose | chemical - endogenous mammalian | Inhibited | 1.44E-13 |
| chloride | chemical - endogenous mammalian | Inhibited | 2.73E-12 |
| desmethylazelastine | chemical - endogenous mammalian | Inhibited | 4.85E-10 |
| nitro-conjugated linoleic acid | chemical - endogenous mammalian | Inhibited | 6.14E-06 |
| 9-nitrooleic acid | chemical - endogenous mammalian | Inhibited | 6.14E-06 |
| 3-hydroxybutyric acid | chemical - endogenous mammalian | Inhibited | 0.0138 |
| colcemid | chemical - endogenous non-mammalian | | 9.59E-12 |
| anacardic acid | chemical - endogenous non-mammalian | | 2.56E-10 |
| imperatorin | chemical - endogenous non-mammalian | | 5.24E-06 |
| phosphoramidon | chemical - endogenous non-mammalian | | 9.06E-06 |
| bis(4-hydroxycinnamoyl)methane | chemical - endogenous non-mammalian | | 0.00549 |
| genipin | chemical - endogenous non-mammalian | | 0.00563 |
| rhodioloside | chemical - endogenous non-mammalian | | 0.0104 |
| ajoene | chemical - endogenous non-mammalian | Activated | 1.34E-08 |
| bis(4-hydroxycinnamoyl)methane | chemical - endogenous non-mammalian | Inhibited | 1.62E-15 |
| bis(4-hydroxycinnamoyl)methane | chemical - endogenous non-mammalian | Inhibited | 2.14E-14 |
| helenalin | chemical - endogenous non-mammalian | Inhibited | 1.23E-13 |
| soraphen-A1alpha | chemical - endogenous non-mammalian | Inhibited | 1.37E-13 |
| sesamin | chemical - endogenous non-mammalian | Inhibited | 5.15E-11 |
| flavone | chemical - endogenous non-mammalian | Inhibited | 2.7E-09 |
| decursin | chemical - endogenous non-mammalian | Inhibited | 7.29E-09 |
| fumagillin | chemical - endogenous non-mammalian | Inhibited | 7.75E-09 |
| geraniol | chemical - endogenous non-mammalian | Inhibited | 0.000987 |

| | | | |
|---|---|---|---|
| dorsomorphin | chemical - kinase inhibitor | | 1.07E-14 |
| IC261 | chemical - kinase inhibitor | | 4.55E-10 |
| CGP 74514A | chemical - kinase inhibitor | | 5.89E-09 |
| SU6657 | chemical - kinase inhibitor | | 3.9E-07 |
| THZ1 | chemical - kinase inhibitor | | 1.17E-05 |
| alpha-cyano-(3-ethoxy-4-hydroxy-5-phenylthiomethyl)cinnamamide | chemical - kinase inhibitor | | 1.59E-05 |
| tyrphostin AG 1295 | chemical - kinase inhibitor | | 0.00188 |
| CBR-470-1 | chemical - kinase inhibitor | | 0.00778 |
| bisindolylmaleimide iv | chemical - kinase inhibitor | Inhibited | 3.58E-13 |
| 4-[(4'-chloro-2'-fluoro)phenylamino]-6,7-dimethoxyquinazoline | chemical - kinase inhibitor | Inhibited | 7.76E-13 |
| JNK-IN-8 | chemical - kinase inhibitor | Inhibited | 2.82E-12 |
| BB4 | chemical - kinase inhibitor | Inhibited | 1.62E-11 |
| Rp-cAMPS | chemical - kinase inhibitor | Inhibited | 9.75E-10 |
| PD 180970 | chemical - kinase inhibitor | Inhibited | 1.48E-07 |
| SB203580 | chemical - kinase inhibitor | Inhibited | 8.79E-07 |
| 6-aminopyrazolopyrimidine derivative compound II | chemical - kinase inhibitor | Inhibited | 1.67E-05 |
| UM101 | chemical - kinase inhibitor | Inhibited | 0.00133 |
| THZ1 | chemical - kinase inhibitor | Inhibited | 0.00326 |
| catecholamine | chemical - other | Activated | 1.72E-11 |
| porphyrin | chemical - other | Inhibited | 8.53E-09 |
| Ac-YVAD-CMK | chemical - protease inhibitor | | 0.125 |
| MMPI-2 | chemical - protease inhibitor | Inhibited | 1.89E-12 |
| isoproterenol | chemical drug | | 8.1E-18 |
| trans-hydroxytamoxifen | chemical drug | | 8.95E-16 |
| acitretin | chemical drug | | 1.77E-15 |
| calcitriol | chemical drug | | 1.65E-14 |
| D-tubocurarine | chemical drug | | 2.9E-14 |
| nadolol | chemical drug | | 8.26E-14 |
| capsaicin | chemical drug | | 1.69E-13 |
| ephedrine | chemical drug | | 1.74E-12 |
| ardeparin | chemical drug | | 1.86E-12 |
| novobiocin | chemical drug | | 2.19E-12 |
| ribavirin | chemical drug | | 2.46E-12 |
| talazoparib | chemical drug | | 3.3E-12 |
| fluphenazine | chemical drug | | 5.63E-12 |
| adaphostin | chemical drug | | 1.22E-11 |
| sorafenib | chemical drug | | 2.35E-11 |
| perphenazine | chemical drug | | 4.85E-11 |
| bendamustine | chemical drug | | 7.27E-10 |
| chlorpropamide | chemical drug | | 2E-09 |
| imipramine | chemical drug | | 2.39E-08 |
| inecalcitol | chemical drug | | 2.6E-08 |
| azathioprine | chemical drug | | 3.8E-08 |

| | | |
|---|---|---|
| pyrilamine | chemical drug | 5.21E-08 |
| ginkgolide B | chemical drug | 5.72E-08 |
| CC 401 | chemical drug | 1.05E-07 |
| tazarotene | chemical drug | 1.07E-07 |
| enzalutamide | chemical drug | 1.58E-07 |
| avasimibe | chemical drug | 2.84E-07 |
| doxorubicin | chemical drug | 2.88E-07 |
| nicorandil | chemical drug | 4.2E-07 |
| pobilukast | chemical drug | 4.36E-07 |
| GLPG0187 | chemical drug | 5.5E-07 |
| imipramine blue | chemical drug | 8.44E-07 |
| bosutinib | chemical drug | 9.36E-07 |
| NCX-4040 | chemical drug | 1.1E-06 |
| repotrectinib | chemical drug | 1.89E-06 |
| chiauranib | chemical drug | 7.69E-06 |
| pirfenidone | chemical drug | 1.12E-05 |
| acetovanillone | chemical drug | 1.81E-05 |
| trans-hydroxytamoxifen | chemical drug | 3.49E-05 |
| cerivastatin | chemical drug | 0.000112 |
| allopurinol | chemical drug | 0.000316 |
| triclosan | chemical drug | 0.000318 |
| eprosartan | chemical drug | 0.000451 |
| semaxinib | chemical drug | 0.000891 |
| felodipine | chemical drug | 0.0011 |
| indinavir | chemical drug | 0.00132 |
| tetomilast | chemical drug | 0.00132 |
| arofylline | chemical drug | 0.00132 |
| L 869298 | chemical drug | 0.00132 |
| ramipril | chemical drug | 0.00133 |
| CTS-1027 | chemical drug | 0.00188 |
| Yi-Gan San | chemical drug | 0.00188 |
| L-826,141 | chemical drug | 0.00249 |
| trimetrexate | chemical drug | 0.00328 |
| oxytetracycline | chemical drug | 0.00423 |
| fomepizole | chemical drug | 0.00549 |
| etodolac | chemical drug | 0.00549 |
| 6-mercaptopurine | chemical drug | 0.00778 |
| carvedilol | chemical drug | 0.00812 |
| enoximone | chemical drug | 0.00812 |
| palbociclib | chemical drug | 0.0104 |
| GR-MD-02 | chemical drug | 0.0107 |
| p38 MAP kinase inhibitor | chemical drug | 0.0107 |
| ticrynafen | chemical drug | 0.0107 |
| quinethazone | chemical drug | 0.0168 |
| tranexamic acid | chemical drug | 0.0173 |
| dexamethasone/tobramycin | chemical drug | 0.0434 |
| HDAC class I inhibitors | chemical drug | 0.0434 |
| pralnacasan | chemical drug | 0.0849 |

| | | | |
|---|---|---|---|
| alitretinoin | chemical drug | Activated | 8.41E-16 |
| mocetinostat | chemical drug | Activated | 2.21E-14 |
| OKI-179 | chemical drug | Activated | 3.42E-14 |
| cephaloridine | chemical drug | Activated | 6.96E-13 |
| bupropion | chemical drug | Activated | 5.57E-12 |
| SEL120 | chemical drug | Activated | 3.92E-10 |
| BCD-115 | chemical drug | Activated | 3.92E-10 |
| neomycin | chemical drug | Activated | 2.17E-09 |
| cirazoline | chemical drug | Activated | 4.95E-09 |
| terbutaline | chemical drug | Activated | 1.27E-08 |
| seocalcitol | chemical drug | Activated | 3.19E-08 |
| fenoterol | chemical drug | Activated | 8.09E-08 |
| nandrolone | chemical drug | Activated | 1.25E-07 |
| estrogen | chemical drug | Activated | 2.23E-07 |
| hydroxyurea | chemical drug | Activated | 2.06E-06 |
| mibolerone | chemical drug | Activated | 2.31E-06 |
| nandrolone decanoate | chemical drug | Activated | 7.15E-06 |
| cinacalcet | chemical drug | Activated | 1.45E-05 |
| testosterone enanthate | chemical drug | Activated | 1.85E-05 |
| pargyline | chemical drug | Activated | 2.92E-05 |
| mocetinostat | chemical drug | Activated | 6.15E-05 |
| carboplatin | chemical drug | Activated | 0.000419 |
| phenylbutazone | chemical drug | Activated | 0.00107 |
| lestaurtinib | chemical drug | Inhibited | 1.95E-18 |
| PD 153035 | chemical drug | Inhibited | 1.08E-17 |
| cerulenin | chemical drug | Inhibited | 1.86E-17 |
| isoflurophate | chemical drug | Inhibited | 2.28E-16 |
| PLX7486 | chemical drug | Inhibited | 2.44E-15 |
| galunisertib | chemical drug | Inhibited | 3.45E-15 |
| bepridil | chemical drug | Inhibited | 3.47E-15 |
| lenalidomide | chemical drug | Inhibited | 4.08E-15 |
| gallium nitrate | chemical drug | Inhibited | 4.38E-15 |
| astemizole | chemical drug | Inhibited | 6.16E-15 |
| glesatinib | chemical drug | Inhibited | 1.09E-14 |
| thalidomide | chemical drug | Inhibited | 1.46E-14 |
| tioconazole | chemical drug | Inhibited | 3.05E-14 |
| escitalopram | chemical drug | Inhibited | 3.39E-14 |
| CEP-37440 | chemical drug | Inhibited | 3.6E-14 |
| sulconazole | chemical drug | Inhibited | 3.61E-14 |
| candesartan cilexetil | chemical drug | Inhibited | 5.85E-14 |
| altiratinib | chemical drug | Inhibited | 6.13E-14 |
| tranylcypromine | chemical drug | Inhibited | 7.72E-14 |
| caffeic acid phenethyl ester | chemical drug | Inhibited | 1.15E-13 |
| NOV1601 | chemical drug | Inhibited | 1.17E-13 |
| larotrectinib | chemical drug | Inhibited | 1.2E-13 |
| pomalidomide | chemical drug | Inhibited | 1.31E-13 |
| dasatinib | chemical drug | Inhibited | 1.58E-13 |
| pentamidine | chemical drug | Inhibited | 2.16E-13 |

| | | | |
|---|---|---|---|
| methoxsalen | chemical drug | Inhibited | 3.21E-13 |
| DS-6051b | chemical drug | Inhibited | 3.93E-13 |
| tivozanib | chemical drug | Inhibited | 3.98E-13 |
| etiracetam | chemical drug | Inhibited | 4.36E-13 |
| belizatinib | chemical drug | Inhibited | 6.42E-13 |
| acetazolamide | chemical drug | Inhibited | 6.43E-13 |
| strychnine | chemical drug | Inhibited | 6.96E-13 |
| ningetinib | chemical drug | Inhibited | 9.41E-13 |
| famitinib | chemical drug | Inhibited | 1.18E-12 |
| rolipram | chemical drug | Inhibited | 1.65E-12 |
| conteltinib | chemical drug | Inhibited | 1.7E-12 |
| VEGF inhibitor drug | chemical drug | Inhibited | 1.58E-11 |
| mimosine | chemical drug | Inhibited | 4.15E-11 |
| astemizole | chemical drug | Inhibited | 7.41E-11 |
| desloratadine | chemical drug | Inhibited | 8.18E-11 |
| pimozide | chemical drug | Inhibited | 8.2E-11 |
| loratadine | chemical drug | Inhibited | 8.53E-11 |
| axitinib | chemical drug | Inhibited | 8.64E-11 |
| calcitriol | chemical drug | Inhibited | 1.04E-10 |
| ticlopidine | chemical drug | Inhibited | 6.16E-10 |
| niclosamide | chemical drug | Inhibited | 7.1E-10 |
| retinol acetate | chemical drug | Inhibited | 1.64E-09 |
| drospirenone | chemical drug | Inhibited | 1.88E-09 |
| silicon phthalocyanine | chemical drug | Inhibited | 3.51E-09 |
| L 778123 | chemical drug | Inhibited | 5.87E-09 |
| pexmetinib | chemical drug | Inhibited | 7.47E-09 |
| fulvestrant | chemical drug | Inhibited | 3.85E-08 |
| FCN-159 | chemical drug | Inhibited | 4.12E-08 |
| argatroban | chemical drug | Inhibited | 5.55E-08 |
| midazolam | chemical drug | Inhibited | 8.05E-08 |
| betaxolol | chemical drug | Inhibited | 1.01E-07 |
| binimetinib | chemical drug | Inhibited | 5.24E-07 |
| epothilone B | chemical drug | Inhibited | 1.82E-06 |
| saracatinib | chemical drug | Inhibited | 1.83E-06 |
| E 6201 | chemical drug | Inhibited | 1.91E-06 |
| melagatran | chemical drug | Inhibited | 2.16E-06 |
| L-779450 | chemical drug | Inhibited | 3.34E-06 |
| enoxaparin | chemical drug | Inhibited | 4.62E-06 |
| CXA-10 | chemical drug | Inhibited | 6.14E-06 |
| CH5126766 | chemical drug | Inhibited | 6.24E-06 |
| nilutamide | chemical drug | Inhibited | 7.3E-06 |
| camostat | chemical drug | Inhibited | 9.58E-06 |
| BMS-690514 | chemical drug | Inhibited | 6.46E-05 |
| halofuginone | chemical drug | Inhibited | 0.000935 |
| vatalanib | chemical drug | Inhibited | 0.00132 |
| T 0070907 | chemical reagent | | 6.67E-14 |
| Rp-8-pCPT-cGMPS-triethylamine | chemical reagent | | 4.28E-11 |

| | | | |
|---|---|---|---|
| (2-(trimethylammonium)ethyl)methanethiosulfonate | chemical reagent | | 3.74E-10 |
| carbonyl cyanide p-(trifluoromethoxy)phenylhydrazone | chemical reagent | | 1.92E-07 |
| 2,2'-dipyridyl disulfide | chemical reagent | | 2.02E-06 |
| curcumin derivative C817 | chemical reagent | | 1.46E-05 |
| mirin | chemical reagent | | 6.87E-05 |
| tributyltin | chemical reagent | | 0.000297 |
| proadifen | chemical reagent | | 0.00148 |
| NSOP00313 | chemical reagent | | 0.00549 |
| L-JNK inhibitor I | chemical reagent | | 0.00549 |
| ACS84 | chemical reagent | | 0.00549 |
| 6-amino-4-(4-phenoxyphenylethylamino)quinazoline | chemical reagent | | 0.00563 |
| polymethyl methacrylate | chemical reagent | | 0.0107 |
| proadifen | chemical reagent | | 0.0434 |
| cobalt chloride | chemical reagent | Activated | 6.53E-16 |
| isatoribine | chemical reagent | Activated | 3.33E-14 |
| 4-((4-(3,4-dichlorophenyl)-1,2,5-thiadiazol-3-yl)oxy)butan-1-ol | chemical reagent | Activated | 4.64E-12 |
| I-BOP | chemical reagent | Activated | 1.26E-11 |
| GnRH-A | chemical reagent | Activated | 2.48E-10 |
| lithocholic acid acetate | chemical reagent | Activated | 1.13E-07 |
| 3,5-dihydroxyphenylglycine | chemical reagent | Activated | 6.07E-07 |
| ITI-078 | chemical reagent | Activated | 5.53E-06 |
| 2-bromoethylamine | chemical reagent | Activated | 0.00114 |
| D-2-amino-5-phosphonovaleric acid | chemical reagent | Inhibited | 2.79E-21 |
| 2-(4-amino-1-isopropyl-1H-pyrazolo[3,4-d]pyrimidin-3-yl)-1H-indol-5-ol | chemical reagent | Inhibited | 6.42E-18 |
| sodium bisulfide | chemical reagent | Inhibited | 1.98E-12 |
| ethylene glycol tetraacetic acid | chemical reagent | Inhibited | 1.12E-11 |
| GW3965 | chemical reagent | Inhibited | 3.24E-10 |
| GGTI-2154 | chemical reagent | Inhibited | 6.14E-09 |
| L-JNK inhibitor I | chemical reagent | Inhibited | 1.06E-05 |
| purvalanol A | chemical reagent | Inhibited | 2.53E-05 |
| latrunculin B | chemical reagent | Inhibited | 0.00061 |
| DMH1 | chemical reagent | Inhibited | 0.00249 |
| pentachlorophenol | chemical toxicant | | 2.77E-15 |
| 1,10-phenanthroline | chemical toxicant | | 6.32E-11 |
| arotinoid acid | chemical toxicant | | 4.05E-08 |
| alpha-amanitin | chemical toxicant | | 1.18E-06 |
| calicheamicin gamma(1)I | chemical toxicant | | 4.35E-05 |
| diquat | chemical toxicant | | 0.000764 |

| | | | |
|---|---|---|---|
| phorbol 12,13-didecanoate | chemical toxicant | | 0.000891 |
| Ro 25-6760 | chemical toxicant | | 0.00267 |
| biochanin A | chemical toxicant | | 0.00663 |
| kainic acid | chemical toxicant | Activated | 5.48E-14 |
| methylnitronitrosoguanidine | chemical toxicant | Activated | 4.51E-13 |
| carbon tetrachloride | chemical toxicant | Activated | 7.02E-13 |
| latrunculin A | chemical toxicant | Activated | 1.12E-09 |
| Salmonella enterica serotype abortus equi lipopolysaccharide | chemical toxicant | Activated | 1.02E-08 |
| aroclor 1254 | chemical toxicant | Activated | 4.1E-08 |
| benzo(a)pyrene 7,8-dihydrodiol | chemical toxicant | Activated | 1.56E-07 |
| 2,3',4,4',5-pentachlorobiphenyl | chemical toxicant | Activated | 3E-07 |
| Salmonella enterica serotype abortus equi lipopolysaccharide | chemical toxicant | Activated | 1.2E-06 |
| dimethylnitrosamine | chemical toxicant | Activated | 2.23E-06 |
| zearalenone | chemical toxicant | Activated | 1.02E-05 |
| thioacetamide | chemical toxicant | Activated | 1.82E-05 |
| dimethylnitrosamine | chemical toxicant | Activated | 3.45E-05 |
| carbon tetrachloride | chemical toxicant | Activated | 0.00021 |
| cytochalasin B | chemical toxicant | Inhibited | 1.51E-09 |
| norethynodrel | chemical toxicant | Inhibited | 4.41E-07 |
| cypermethrin | chemical toxicant | Inhibited | 1.15E-05 |
| VitaminD3-VDR-RXR | complex | | 4.07E-16 |
| T3-TR-RXR | complex | | 1.57E-14 |
| SWI-SNF | complex | | 9.42E-14 |
| PKR dimer | complex | | 2.05E-11 |
| APPBP1/UBA3 Complex | complex | | 1.65E-10 |
| PDGF-DD | complex | | 4.28E-08 |
| SWI-SNF | complex | | 1.61E-07 |
| MEF2D-NFAT2-p300 | complex | | 1.65E-07 |
| Integrin alpha 6 beta 1 | complex | | 1.34E-06 |
| Glycoprotein 1B | complex | | 1.18E-05 |
| Laminin (complex) | complex | | 0.000451 |
| IgG2b | complex | | 0.00131 |
| histone deacetylase | complex | | 0.0016 |
| PDGF-Rβ dimer | complex | | 0.00184 |
| Bcl10-Card10-Malt1 | complex | | 0.00326 |
| PDGF-Rβ dimer | complex | | 0.0434 |
| Laminin (complex) | complex | Activated | 1.53E-17 |
| Ige | complex | Activated | 1.71E-14 |
| Nodal receptor | complex | Activated | 4.32E-14 |
| IGg-Rheumatoid factor | complex | Activated | 5.76E-13 |
| NMDA Receptor | complex | Activated | 1.48E-12 |
| CD247 dimer | complex | Activated | 2.95E-10 |
| CAK | complex | Activated | 3.2E-09 |
| EIF3 | complex | Activated | 3.92E-09 |
| Esr1-Esr1-estrogen-estrogen | complex | Activated | 4.19E-08 |
| NFkB1-CRel | complex | Activated | 7.89E-06 |

| | | | |
|---|---|---|---|
| cyclin h/cdk7 | complex | Activated | 4.26E-05 |
| EIF4F | complex | Activated | 5.26E-05 |
| APC (complex) | complex | Activated | 0.000149 |
| histone deacetylase | complex | Inhibited | 1.7E-09 |
| I kappa b kinase | complex | Inhibited | 8.7E-06 |
| TIMP1 | cytokine | | 0.00267 |
| CXCL11 | cytokine | | 0.00549 |
| CSF2 | cytokine | Activated | 2.77E-18 |
| CXCL12 | cytokine | Activated | 5.07E-18 |
| CSF1 | cytokine | Activated | 7.14E-18 |
| CSF2 | cytokine | Activated | 1.92E-17 |
| IL1B | cytokine | Activated | 1.08E-14 |
| IL20 | cytokine | Activated | 4.19E-14 |
| WNT3A | cytokine | Activated | 1.31E-13 |
| PF4 | cytokine | Activated | 1.54E-09 |
| CXCL12 | cytokine | Activated | 3.05E-05 |
| CYP27A1 | enzyme | | 2.51E-15 |
| FBXW11 | enzyme | | 2.6E-15 |
| CYP19A1 | enzyme | | 2.98E-15 |
| RECQL5 | enzyme | | 3.36E-13 |
| RNF19A | enzyme | | 2.08E-12 |
| UHRF2 | enzyme | | 4.64E-12 |
| CUL1 | enzyme | | 4.85E-12 |
| UBE4B | enzyme | | 1.24E-11 |
| OTUB1 | enzyme | | 1.66E-11 |
| RCHY1 | enzyme | | 5.35E-11 |
| RALA | enzyme | | 7.44E-11 |
| POR | enzyme | | 7.63E-11 |
| RALB | enzyme | | 1.03E-10 |
| MLH1 | enzyme | | 1.07E-10 |
| BIRC2 | enzyme | | 2.13E-10 |
| RAD50 | enzyme | | 1.21E-09 |
| RGS6 | enzyme | | 1.52E-09 |
| HPRT1 | enzyme | | 5.18E-08 |
| CAB39 | enzyme | | 2.06E-07 |
| MGMT | enzyme | | 1.96E-06 |
| ARIH2 | enzyme | | 6.89E-06 |
| IHH | enzyme | | 0.000155 |
| AGA | enzyme | | 0.000764 |
| TREX1 | enzyme | | 0.00107 |
| WWOX | enzyme | | 0.00137 |
| IHH | enzyme | | 0.00151 |
| CYP1B1 | enzyme | | 0.00244 |
| PLCD1 | enzyme | | 0.00388 |
| GSR | enzyme | | 0.0042 |
| HERC5 | enzyme | | 0.00549 |
| PARP14 | enzyme | | 0.00563 |
| HAS2 | enzyme | | 0.00563 |

| | | | |
|---|---|---|---|
| NDUFA13 | enzyme | | 0.00778 |
| GNA11 | enzyme | | 0.00778 |
| MBD4 | enzyme | | 0.0107 |
| CHI3L1 | enzyme | | 0.0107 |
| ARF1 | enzyme | | 0.0107 |
| PDE4B | enzyme | | 0.0251 |
| SI | enzyme | | 0.0434 |
| KDM4B | enzyme | Activated | 1.46E-14 |
| SMYD3 | enzyme | Activated | 1.56E-14 |
| RAP2A | enzyme | Activated | 3.45E-14 |
| P4HB | enzyme | Activated | 1.02E-13 |
| HMGA2 | enzyme | Activated | 1.62E-13 |
| PPME1 | enzyme | Activated | 5.6E-13 |
| KDM1A | enzyme | Activated | 5.94E-13 |
| PELI1 | enzyme | Activated | 2.67E-10 |
| GNAS | enzyme | Activated | 8.96E-10 |
| SETD7 | enzyme | Activated | 5.76E-09 |
| TERT | enzyme | Activated | 1.44E-08 |
| PLA2G5 | enzyme | Activated | 1.19E-07 |
| KDM8 | enzyme | Activated | 2.07E-06 |
| PTGES3 | enzyme | Activated | 2.16E-06 |
| RAC3 | enzyme | Activated | 3.64E-06 |
| PPID | enzyme | Activated | 4.54E-06 |
| AKR1C3 | enzyme | Activated | 3.09E-05 |
| HINT1 | enzyme | Inhibited | 4.15E-16 |
| PPIF | enzyme | Inhibited | 6.65E-16 |
| COMT | enzyme | Inhibited | 2.08E-14 |
| PLCL2 | enzyme | Inhibited | 2.41E-12 |
| GNG7 | enzyme | Inhibited | 7.17E-12 |
| FBXO4 | enzyme | Inhibited | 5.8E-08 |
| CHFR | enzyme | Inhibited | 1.66E-06 |
| DNMT3A | enzyme | Inhibited | 2.6E-06 |
| DNMT3A | enzyme | Inhibited | 5.9E-06 |
| DNMT3B | enzyme | Inhibited | 0.00033 |
| DNASE2 | enzyme | Inhibited | 0.00267 |
| KDM3B | enzyme | Inhibited | 0.00851 |
| RUNX1-RUNX1T1 | fusion gene/product | | 0.00328 |
| DRD4 | G-protein coupled receptor | | 1.27E-12 |
| HTR6 | G-protein coupled receptor | | 5.12E-11 |
| FFAR3 | G-protein coupled receptor | | 3.19E-10 |
| NMBR | G-protein coupled receptor | | 4.6E-06 |
| AVPR2 | G-protein coupled receptor | | 1.03E-05 |
| GPR68 | G-protein coupled receptor | | 0.0173 |
| FZD7 | G-protein coupled receptor | | 0.0173 |
| ADGRE5 | G-protein coupled receptor | Activated | 3.15E-13 |
| CXCR3 | G-protein coupled receptor | Activated | 3.2E-12 |
| P2RY4 | G-protein coupled receptor | Activated | 4.27E-11 |
| F2R | G-protein coupled receptor | Activated | 3.57E-06 |

| | | | |
|---|---|---|---|
| CASR | G-protein coupled receptor | Activated | 7.14E-05 |
| PTGER2 | G-protein coupled receptor | Activated | 0.0012 |
| ADGRG3 | G-protein coupled receptor | Inhibited | 2.46E-13 |
| ACKR2 | G-protein coupled receptor | Inhibited | 0.00203 |
| PI4K | group | | 1.47E-17 |
| MEF2 | group | | 8.62E-15 |
| Lysosomal Protease | group | | 5.91E-14 |
| Alpha Actinin | group | | 6.63E-14 |
| Gata | group | | 6.93E-14 |
| Hdac | group | | 1.93E-13 |
| Focal adhesion kinase | group | | 2E-13 |
| Cdc2 | group | | 1.61E-11 |
| B56 | group | | 6.99E-11 |
| ALDH | group | | 3.66E-10 |
| Pkar2 | group | | 8.68E-09 |
| Calcineurin A | group | | 1.71E-08 |
| MEF2 | group | | 2.04E-07 |
| DNA-methyltransferase | group | | 2.34E-07 |
| DUB | group | | 0.000018 |
| ADRB | group | | 0.000037 |
| tryptase | group | | 0.000111 |
| G-Actin | group | | 0.00233 |
| Ikk (family) | group | | 0.00549 |
| Diap | group | | 0.00549 |
| Nfat (family) | group | Activated | 8.55E-19 |
| Vegf | group | Activated | 7.69E-18 |
| E2f | group | Activated | 7.71E-18 |
| estrogen receptor | group | Activated | 4.22E-15 |
| PDZGEF | group | Activated | 3.36E-13 |
| Raf | group | Activated | 8.64E-13 |
| DRD1/5 | group | Activated | 8.04E-12 |
| receptor protein tyrosine kinase | group | Activated | 1.3E-11 |
| IL-1R | group | Activated | 2.02E-10 |
| Par | group | Activated | 9.4E-10 |
| SYK/ZAP | group | Activated | 1.09E-09 |
| Vegf | group | Activated | 1.85E-09 |
| Gα12/13 | group | Activated | 3.98E-07 |
| MIR99A-LET7C-MIR125B2 | group | Inhibited | 2.87E-16 |
| MIR100-LET7A2-MIR125B1 | group | Inhibited | 2.87E-16 |
| MIRLET7 | group | Inhibited | 3.94E-16 |
| SFRP | group | Inhibited | 7.93E-15 |
| Lefty | group | Inhibited | 1.48E-13 |
| Npm | group | Inhibited | 2.13E-13 |
| Endophilin | group | Inhibited | 1.21E-12 |
| Rb | group | Inhibited | 2.08E-12 |
| MIRLET7 | group | Inhibited | 7.37E-11 |
| MIR99A-LET7C-MIR125B2 | group | Inhibited | 1.49E-09 |
| MIR100-LET7A2-MIR125B1 | group | Inhibited | 1.49E-09 |

| | | | |
|---|---|---|---|
| Gamma tubulin | group | Inhibited | 1.64E-08 |
| Alpha 1 antitrypsin | group | Inhibited | 8.92E-07 |
| FGF3 | growth factor | | 1.13E-11 |
| GDF9 | growth factor | | 0.000451 |
| BMP4 | growth factor | Activated | 1.35E-15 |
| NDP | growth factor | Activated | 1.46E-14 |
| HDGF | growth factor | Activated | 3.6E-13 |
| FGF4 | growth factor | Activated | 1.38E-09 |
| HGF | growth factor | Activated | 4.32E-08 |
| FGF6 | growth factor | Activated | 1.38E-07 |
| FGF3 | growth factor | Activated | 0.000111 |
| JAG1 | growth factor | Activated | 0.000419 |
| RASA3 | ion channel | | 7.54E-12 |
| Cacnb1 | ion channel | | 0.00549 |
| CACNA2D1 | ion channel | | 0.00549 |
| DYRK1A | kinase | | 6.81E-19 |
| MAK | kinase | | 2.03E-18 |
| DYRK1A | kinase | | 1.43E-16 |
| STK38L | kinase | | 4.34E-15 |
| MYLK2 | kinase | | 4.49E-13 |
| PAK6 | kinase | | 2.32E-11 |
| CAMK2G | kinase | | 5.4E-11 |
| AURKB | kinase | | 6.31E-10 |
| STRADA | kinase | | 3.53E-08 |
| Brd4 | kinase | | 3.24E-07 |
| CDC7 | kinase | | 8.45E-07 |
| STK11 | kinase | | 3.65E-06 |
| MAPK1 | kinase | | 2.04E-05 |
| CPNE3 | kinase | | 2.71E-05 |
| ATM | kinase | | 4.79E-05 |
| CHEK1 | kinase | | 0.00011 |
| MAPKAPK2 | kinase | | 0.000112 |
| AURKB | kinase | | 0.000151 |
| TBK1 | kinase | | 0.00037 |
| PLK2 | kinase | | 0.000384 |
| CDK12 | kinase | | 0.000764 |
| NTRK2 | kinase | | 0.0013 |
| FGFR2 | kinase | | 0.00156 |
| PTK6 | kinase | | 0.0107 |
| TRPM7 | kinase | | 0.0434 |
| HIPK3 | kinase | | 0.0434 |
| PKM | kinase | Activated | 3.18E-21 |
| ERBB2 | kinase | Activated | 2.01E-17 |
| PRKCZ | kinase | Activated | 6.14E-16 |
| MAP3K8 | kinase | Activated | 1.02E-14 |
| STK26 | kinase | Activated | 1.37E-14 |
| MAP3K3 | kinase | Activated | 4.87E-14 |
| JAK1 | kinase | Activated | 2.37E-13 |

| | | | |
|---|---|---|---|
| MAPK11 | kinase | Activated | 2.78E-13 |
| MAP3K2 | kinase | Activated | 9.83E-13 |
| CAMK2A | kinase | Activated | 5.13E-12 |
| PRKCI | kinase | Activated | 1.14E-10 |
| MOS | kinase | Activated | 3.82E-10 |
| CKS1B | kinase | Activated | 2.12E-08 |
| TAOK2 | kinase | Activated | 5.62E-08 |
| BRD4 | kinase | Activated | 2.16E-07 |
| Brd4 | kinase | Activated | 3.01E-06 |
| MAP2K1 | kinase | Activated | 5.44E-06 |
| CCNK | kinase | Activated | 3.96E-05 |
| CRKL | kinase | Activated | 0.00328 |
| MARK2 | kinase | Inhibited | 2.21E-15 |
| CDK8 | kinase | Inhibited | 9.39E-10 |
| ITPKB | kinase | Inhibited | 5.13E-09 |
| CDKN1A | kinase | Inhibited | 2.56E-07 |
| CAMK2N1 | kinase | Inhibited | 2.63E-07 |
| LATS1 | kinase | Inhibited | 2.07E-06 |
| DYRK1A | kinase | Inhibited | 2.73E-05 |
| PLK1 | kinase | Inhibited | 0.00534 |
| RARA | ligand-dependent nuclear receptor | | 4.59E-05 |
| ESR1 | ligand-dependent nuclear receptor | Activated | 3.15E-14 |
| RORC | ligand-dependent nuclear receptor | Activated | 5.24E-12 |
| AR | ligand-dependent nuclear receptor | Activated | 3.76E-05 |
| NR2E1 | ligand-dependent nuclear receptor | Inhibited | 2.76E-05 |
| miR-515-5p (and other miRNAs w/seed UCUCCAA) | mature microRNA | | 0.0107 |
| miR-382-5p (miRNAs w/seed AAGUUGU) | mature microRNA | | 0.0173 |
| miR-105-5p (and other miRNAs w/seed CAAAUGC) | mature microRNA | | 0.0434 |
| miR-148a-3p (and other miRNAs w/seed CAGUGCA) | mature microRNA | Inhibited | 0.00022 |
| miR-34a-5p (and other miRNAs w/seed GGCAGUG) | mature microRNA | Inhibited | 0.00195 |
| miR-24-3p (and other miRNAs w/seed GGCUCAG) | mature microRNA | Inhibited | 0.00653 |
| mir-192 | microRNA | | 2.42E-07 |
| mir-19 | microRNA | | 0.000188 |
| mir-503 | microRNA | | 0.00392 |
| mir-340 | microRNA | | 0.0434 |
| let-7 | microRNA | Inhibited | 1.95E-14 |
| mir-17 | microRNA | Inhibited | 2.57E-11 |
| let-7 | microRNA | Inhibited | 1.99E-09 |
| mir-21 | microRNA | Inhibited | 4.54E-06 |
| ARRDC3 | other | | 6.32E-14 |
| TG | other | | 3.4E-13 |
| RALGAPB | other | | 6.62E-13 |

| | | |
|---|---|---|
| INCENP | other | 2.67E-11 |
| TNRC6A | other | 3.89E-11 |
| DDIT4 | other | 5.35E-11 |
| ELAVL1 | other | 5.19E-10 |
| YWHAQ | other | 4.71E-09 |
| CUL4A | other | 3.74E-08 |
| FRAT1 | other | 4.16E-08 |
| FADD | other | 5.72E-08 |
| HLA-A | other | 1.03E-07 |
| SELPLG | other | 1.07E-07 |
| BNIP3L | other | 1.65E-07 |
| USO1 | other | 2.21E-07 |
| NLRP3 | other | 8.01E-06 |
| CD59 | other | 1.09E-05 |
| SYF2 | other | 1.15E-05 |
| GP9 | other | 1.33E-05 |
| SV2C | other | 1.75E-05 |
| CDC6 | other | 1.96E-05 |
| PVR | other | 3.61E-05 |
| PALB2 | other | 4.79E-05 |
| VWC2 | other | 4.99E-05 |
| CBX7 | other | 8.19E-05 |
| SDC3 | other | 0.000147 |
| HOTAIR | other | 0.000632 |
| MKI67 | other | 0.000764 |
| VGLL3 | other | 0.00137 |
| SOCS6 | other | 0.00174 |
| DCTN4 | other | 0.00188 |
| LAMA1 | other | 0.00188 |
| DLL4 | other | 0.00203 |
| TNFAIP6 | other | 0.00217 |
| CLDN7 | other | 0.00249 |
| SERPINH1 | other | 0.0025 |
| Ptgs2os2 | other | 0.0025 |
| ATG13 | other | 0.00423 |
| FBXO42 | other | 0.00534 |
| SFN | other | 0.00556 |
| CDKN1C | other | 0.00563 |
| VGLL3 | other | 0.00663 |
| CDH2 | other | 0.00778 |
| Nppb | other | 0.0107 |
| LSP1 | other | 0.0107 |
| SCIN | other | 0.0107 |
| RB1CC1 | other | 0.0134 |
| TRIM38 | other | 0.0168 |
| TRIM58 | other | 0.0173 |
| C4A/C4B | other | 0.0251 |
| KRT1 | other | 0.0434 |

| | | | |
|---|---|---|---|
| FSIP1 | other | | 0.0434 |
| LINC02599 | other | | 0.0434 |
| OPCML | other | | 0.0434 |
| USP50 | other | | 0.0434 |
| LINC00261 | other | | 0.0434 |
| CEP78 | other | | 0.0434 |
| SYN2 | other | | 0.0434 |
| NLRP9 | other | | 0.0434 |
| KRT3 | other | | 0.0434 |
| CD151 | other | Activated | 6.35E-22 |
| CALCA | other | Activated | 4.15E-20 |
| POMC | other | Activated | 1.21E-17 |
| APBB1IP | other | Activated | 1.39E-13 |
| KLHL42 | other | Activated | 4.53E-13 |
| RADIL | other | Activated | 5.2E-13 |
| EFHD2 | other | Activated | 4.79E-12 |
| BLNK | other | Activated | 7.82E-12 |
| IL34 | other | Activated | 1.3E-11 |
| RABL6 | other | Activated | 2.74E-11 |
| BAK1 | other | Activated | 3.53E-11 |
| BEX2 | other | Activated | 4.81E-11 |
| HOTAIR | other | Activated | 5.09E-11 |
| S100A8 | other | Activated | 1.48E-10 |
| S100A9 | other | Activated | 5.18E-10 |
| SORBS3 | other | Activated | 7.1E-10 |
| CRK | other | Activated | 7.49E-10 |
| MBL2 | other | Activated | 1.05E-09 |
| EP400 | other | Activated | 1.09E-09 |
| IGFBP6 | other | Activated | 1.81E-09 |
| RETNLB | other | Activated | 2.75E-09 |
| RANBP9 | other | Activated | 7.58E-09 |
| LAMTOR3 | other | Activated | 3.39E-08 |
| FBLN5 | other | Activated | 1.84E-07 |
| ARHGEF11 | other | Activated | 2.24E-07 |
| CYTH2 | other | Activated | 4.7E-07 |
| INIP | other | Activated | 5.93E-07 |
| INTS3 | other | Activated | 5.93E-07 |
| SASH1 | other | Activated | 7.88E-07 |
| FGD1 | other | Activated | 2.66E-06 |
| BOP1 | other | Activated | 3.79E-06 |
| RNF123 | other | Activated | 5.33E-06 |
| CLEC12A | other | Activated | 1.81E-05 |
| BRAT1 | other | Activated | 3.41E-05 |
| ACTB | other | Activated | 3.56E-05 |
| SPTAN1 | other | Activated | 0.000043 |
| PTH | other | Activated | 4.47E-05 |
| SASH1 | other | Activated | 0.00023 |
| Bvht | other | Activated | 0.000333 |

| | | | |
|---|---|---|---|
| ARHGAP21 | other | Activated | 0.000459 |
| STING1 | other | Activated | 0.000556 |
| SAMSN1 | other | Activated | 0.000561 |
| DSCAM | other | Activated | 0.00251 |
| CDH11 | other | Activated | 0.00653 |
| EWSR1 | other | Activated | 0.0101 |
| TRIM65 | other | Inhibited | 3.32E-18 |
| LEMD3 | other | Inhibited | 1.59E-16 |
| SAFB | other | Inhibited | 8.26E-16 |
| CBX7 | other | Inhibited | 9.66E-16 |
| SAFB2 | other | Inhibited | 3.86E-15 |
| IFRD1 | other | Inhibited | 9.9E-15 |
| KRIT1 | other | Inhibited | 1.2E-14 |
| SPARC | other | Inhibited | 1.24E-14 |
| TINCR | other | Inhibited | 1.28E-14 |
| CGN | other | Inhibited | 2.8E-13 |
| SPINT2 | other | Inhibited | 2.98E-13 |
| SERPING1 | other | Inhibited | 1.71E-12 |
| AMOT | other | Inhibited | 2.98E-12 |
| TRIM65 | other | Inhibited | 4.54E-10 |
| SERPIND1 | other | Inhibited | 6.96E-10 |
| CCNC | other | Inhibited | 2.61E-09 |
| PINX1 | other | Inhibited | 9.92E-09 |
| SPINT1 | other | Inhibited | 1.26E-08 |
| Cdc2b | other | Inhibited | 1.37E-08 |
| KLHL40 | other | Inhibited | 1.78E-08 |
| SPRY4 | other | Inhibited | 1.87E-08 |
| Irgm1 | other | Inhibited | 2.36E-07 |
| MMS22L | other | Inhibited | 5.93E-07 |
| FRMD6 | other | Inhibited | 7.36E-07 |
| Irgm1 | other | Inhibited | 0.000002 |
| SPARC | other | Inhibited | 4.56E-06 |
| SFPQ | other | Inhibited | 6.37E-06 |
| SERPINA5 | other | Inhibited | 7.49E-06 |
| TFPI2 | other | Inhibited | 9.84E-06 |
| RASSF6 | other | Inhibited | 0.00002 |
| SPINK5 | other | Inhibited | 0.000289 |
| 1810019D21Rik | other | Inhibited | 0.000941 |
| PCBP2 | other | Inhibited | 0.000947 |
| CDON | other | Inhibited | 0.00132 |
| RCE1 | peptidase | | 6.4E-13 |
| F10 | peptidase | | 5.9E-12 |
| CAST | peptidase | | 1.63E-10 |
| EPS8 | peptidase | | 4.84E-09 |
| USP24 | peptidase | | 1.35E-07 |
| USP12 | peptidase | | 3.02E-07 |
| CAPN3 | peptidase | | 0.00328 |
| USP1 | peptidase | | 0.00549 |

| | | | |
|---|---|---|---|
| ZFYVE9 | peptidase | | 0.0107 |
| USP6 | peptidase | Activated | 1.45E-13 |
| CTSC | peptidase | Activated | 6.87E-12 |
| GZMB | peptidase | Activated | 6.79E-11 |
| ST14 | peptidase | Activated | 4.96E-10 |
| F11 | peptidase | Activated | 1.65E-09 |
| UCHL5 | peptidase | Activated | 2.89E-09 |
| USP5 | peptidase | Activated | 5.13E-07 |
| F2 | peptidase | Activated | 1.21E-05 |
| ADAMTS18 | peptidase | Activated | 3.96E-05 |
| PLG | peptidase | Activated | 0.00386 |
| CCAR2 | peptidase | Inhibited | 6.24E-08 |
| PSMB11 | peptidase | Inhibited | 7.29E-08 |
| USP38 | peptidase | Inhibited | 6.44E-07 |
| CPN1 | peptidase | Inhibited | 0.000422 |
| PPM1A | phosphatase | | 1.69E-11 |
| CDC25B | phosphatase | | 1.08E-10 |
| CDC14A | phosphatase | | 1.05E-09 |
| CDC14B | phosphatase | | 1.09E-09 |
| PPEF2 | phosphatase | | 2.38E-06 |
| PPP3R1 | phosphatase | | 0.00152 |
| TNS3 | phosphatase | Activated | 9.72E-13 |
| PTPN22 | phosphatase | Inhibited | 7.96E-18 |
| PTPN9 | phosphatase | Inhibited | 1.58E-17 |
| SOX2 | transcription regulator | | 5.31E-18 |
| GATA1 | transcription regulator | | 1.2E-14 |
| BCL6 | transcription regulator | | 4.71E-14 |
| ID2 | transcription regulator | | 8.31E-13 |
| MEF2C | transcription regulator | | 1.03E-12 |
| NFKB1 | transcription regulator | | 1.03E-12 |
| ZBTB16 | transcription regulator | | 1.22E-12 |
| HMGA1 | transcription regulator | | 2.98E-12 |
| SOX2 | transcription regulator | | 3.35E-12 |
| SMARCA4 | transcription regulator | | 3.56E-12 |
| CALR | transcription regulator | | 4.18E-12 |
| KLF8 | transcription regulator | | 6.29E-12 |
| H2AX | transcription regulator | | 1.04E-11 |
| TAF1 | transcription regulator | | 2.19E-11 |
| FLI1 | transcription regulator | | 2.54E-11 |
| HEXIM1 | transcription regulator | | 6.92E-11 |
| SMARCA2 | transcription regulator | | 3.83E-10 |
| NKX2-5 | transcription regulator | | 5.11E-10 |
| SPI1 | transcription regulator | | 2.1E-09 |
| CBFA2T3 | transcription regulator | | 3.04E-09 |
| GATA1 | transcription regulator | | 3.4E-09 |
| TCF7L2 | transcription regulator | | 8.06E-09 |
| TMF1 | transcription regulator | | 1.44E-08 |
| GATA1 | transcription regulator | | 5.59E-08 |

| | | | |
|---|---|---|---|
| IKZF1 | transcription regulator | | 1.02E-07 |
| MEF2D | transcription regulator | | 3.19E-07 |
| GATA2 | transcription regulator | | 1.83E-06 |
| SMARCB1 | transcription regulator | | 3.85E-06 |
| SOX2 | transcription regulator | | 1.14E-05 |
| TONSL | transcription regulator | | 1.34E-05 |
| VDR | transcription regulator | | 1.63E-05 |
| KDM5B | transcription regulator | | 2.23E-05 |
| WWTR1 | transcription regulator | | 9.04E-05 |
| NOTCH4 | transcription regulator | | 0.000118 |
| HDAC2 | transcription regulator | | 0.000208 |
| UXT | transcription regulator | | 0.000369 |
| SMARCA2 | transcription regulator | | 0.000459 |
| CBFB | transcription regulator | | 0.000591 |
| GABPA | transcription regulator | | 0.000829 |
| ZNF217 | transcription regulator | | 0.000855 |
| FOXN4 | transcription regulator | | 0.000855 |
| FLI1 | transcription regulator | | 0.000987 |
| MED16 | transcription regulator | | 0.00132 |
| SKI | transcription regulator | | 0.00137 |
| E2F7 | transcription regulator | | 0.00148 |
| UBP1 | transcription regulator | | 0.00188 |
| NKX2-5 | transcription regulator | | 0.00191 |
| TOB1 | transcription regulator | | 0.00442 |
| HAND2 | transcription regulator | | 0.00442 |
| ZHX2 | transcription regulator | | 0.00549 |
| LPXN | transcription regulator | | 0.00549 |
| HMG20B | transcription regulator | | 0.00549 |
| RBBP7 | transcription regulator | | 0.00549 |
| NEO1 | transcription regulator | | 0.00563 |
| ARID2 | transcription regulator | | 0.0107 |
| DR1 | transcription regulator | | 0.0251 |
| DEPDC1 | transcription regulator | | 0.0434 |
| TFCP2L1 | transcription regulator | | 0.0434 |
| TBX18 | transcription regulator | | 0.0434 |
| HDAC2 | transcription regulator | Activated | 2.41E-15 |
| TP63 | transcription regulator | Activated | 5.54E-15 |
| CTNNB1 | transcription regulator | Activated | 2.23E-14 |
| MITF | transcription regulator | Activated | 3.11E-14 |
| SMAD2 | transcription regulator | Activated | 6.76E-13 |
| CBX4 | transcription regulator | Activated | 1.51E-12 |
| TBX2 | transcription regulator | Activated | 4.49E-12 |
| SP3 | transcription regulator | Activated | 7.45E-11 |
| ZNF217 | transcription regulator | Activated | 1.48E-10 |
| PURA | transcription regulator | Activated | 1.59E-09 |
| SERTAD1 | transcription regulator | Activated | 2.16E-09 |
| ETV1 | transcription regulator | Activated | 5.04E-09 |
| NFIC | transcription regulator | Activated | 9.58E-09 |

| | | | |
|---|---|---|---|
| TFDP2 | transcription regulator | Activated | 1.64E-08 |
| E2F1 | transcription regulator | Activated | 4E-08 |
| UHRF1 | transcription regulator | Activated | 1.33E-07 |
| CCND1 | transcription regulator | Activated | 1.72E-07 |
| SMARCE1 | transcription regulator | Activated | 2.23E-07 |
| TP63 | transcription regulator | Activated | 8.92E-07 |
| CCNE1 | transcription regulator | Activated | 9.69E-07 |
| YAP1 | transcription regulator | Activated | 2.28E-06 |
| E2F2 | transcription regulator | Activated | 5.84E-06 |
| EHF | transcription regulator | Activated | 1.03E-05 |
| E2F3 | transcription regulator | Activated | 1.04E-05 |
| SMARCA4 | transcription regulator | Activated | 4.96E-05 |
| NOTCH3 | transcription regulator | Activated | 0.000227 |
| FOXM1 | transcription regulator | Activated | 0.000377 |
| KDM3A | transcription regulator | Activated | 0.000388 |
| TAL1 | transcription regulator | Activated | 0.000485 |
| E2F2 | transcription regulator | Activated | 0.000737 |
| ZNF281 | transcription regulator | Activated | 0.00109 |
| TCF7L2 | transcription regulator | Activated | 0.00144 |
| REL | transcription regulator | Activated | 0.00155 |
| HOXA9 | transcription regulator | Activated | 0.00212 |
| WBP2 | transcription regulator | Activated | 0.00224 |
| ETS1 | transcription regulator | Inhibited | 1.27E-23 |
| MEOX1 | transcription regulator | Inhibited | 1.94E-15 |
| CDKN2A | transcription regulator | Inhibited | 3.06E-15 |
| HDAC7 | transcription regulator | Inhibited | 3.07E-15 |
| ZFHX3 | transcription regulator | Inhibited | 3.08E-15 |
| SOX11 | transcription regulator | Inhibited | 4.21E-15 |
| HDAC1 | transcription regulator | Inhibited | 1.28E-13 |
| MEOX1 | transcription regulator | Inhibited | 2.13E-13 |
| ZFP36 | transcription regulator | Inhibited | 3.17E-13 |
| CDKN2A | transcription regulator | Inhibited | 3.28E-13 |
| E2F6 | transcription regulator | Inhibited | 5.85E-12 |
| NUPR1 | transcription regulator | Inhibited | 1.99E-10 |
| SPDEF | transcription regulator | Inhibited | 5.07E-09 |
| NFX1 | transcription regulator | Inhibited | 7.26E-09 |
| RBL1 | transcription regulator | Inhibited | 6.4E-08 |
| TRERF1 | transcription regulator | Inhibited | 1.22E-07 |
| DNMT3L | transcription regulator | Inhibited | 1.7E-07 |
| ZIC2 | transcription regulator | Inhibited | 2.29E-07 |
| PSIP1 | transcription regulator | Inhibited | 5.8E-07 |
| NUPR1 | transcription regulator | Inhibited | 7.31E-07 |
| HOXA4 | transcription regulator | Inhibited | 1.67E-06 |
| HEYL | transcription regulator | Inhibited | 1.71E-05 |
| FOXH1 | transcription regulator | Inhibited | 1.85E-05 |
| RB1 | transcription regulator | Inhibited | 8.94E-05 |
| FOXP3 | transcription regulator | Inhibited | 0.000295 |
| PTTG1 | transcription regulator | Inhibited | 0.000295 |

| | | | |
|---|---|---|---|
| HDAC11 | transcription regulator | Inhibited | 0.000386 |
| GLIS2 | transcription regulator | Inhibited | 0.000815 |
| ZFP36 | transcription regulator | Inhibited | 0.00118 |
| NOSTRIN | transcription regulator | Inhibited | 0.00227 |
| Bc1 | translation regulator | | 0.0107 |
| EIF4G1 | translation regulator | Activated | 0.00217 |
| TNFRSF1A | transmembrane receptor | | 8.42E-12 |
| FAS | transmembrane receptor | | 0.000112 |
| PLAUR | transmembrane receptor | | 0.000787 |
| FCGR1A | transmembrane receptor | | 0.0025 |
| ENG | transmembrane receptor | | 0.00267 |
| GP1BA | transmembrane receptor | Activated | 4.49E-15 |
| IL21R | transmembrane receptor | Activated | 3.99E-14 |
| ITGAV | transmembrane receptor | Activated | 5.41E-14 |
| ITGB3 | transmembrane receptor | Activated | 2.13E-13 |
| KLRF1 | transmembrane receptor | Activated | 3.79E-13 |
| CLEC1B | transmembrane receptor | Activated | 5.51E-13 |
| ITGB1 | transmembrane receptor | Activated | 1.13E-12 |
| IL17RA | transmembrane receptor | Activated | 6.02E-10 |
| ITGA2 | transmembrane receptor | Activated | 1E-08 |
| TREM1 | transmembrane receptor | Activated | 1.14E-08 |
| FCGR1A | transmembrane receptor | Activated | 1.46E-08 |
| IL1R1 | transmembrane receptor | Activated | 0.00114 |
| SNX9 | transporter | | 4.42E-12 |
| RACGAP1 | transporter | | 6.12E-06 |
| SYN1 | transporter | | 0.0434 |
| GJA3 | transporter | | 0.0434 |
| SNX5 | transporter | Inhibited | 4.84E-13 |
| AMBP | transporter | Inhibited | 3.04E-09 |

**Supplementary Table 3. Etoposide-induced differences in master regulators in KR cells**

| Master Regulator | Molecule Type | Predicted Activation | p-value of overlap |
|---|---|---|---|
| CD151 | other | Activated | 6.88E-25 |
| PKM | kinase | Activated | 4.98E-21 |
| D-2-amino-5-phosphonovaleric acid | chemical reagent | Inhibited | 5.44E-21 |
| CAMLG | other | Activated | 9.84E-21 |
| Nfat (family) | group | Activated | 1.45E-20 |
| CDH1 | other | | 4.4E-20 |
| NRG4 | growth factor | Activated | 1.24E-19 |
| PTPN9 | phosphatase | Inhibited | 1.76E-19 |
| IGFBP2 | other | Activated | 1.77E-19 |
| ETV7 | transcription regulator | Activated | 3.02E-19 |
| TOB1 | transcription regulator | Inhibited | 3.33E-19 |
| SMAD5 | transcription regulator | Activated | 3.47E-19 |
| DYRK1A | kinase | | 7.3E-19 |
| TGM2 | enzyme | Activated | 1E-18 |
| CALCA | other | | 1.33E-18 |
| Laminin (complex) | complex | Activated | 1.43E-18 |
| pelitinib | chemical drug | Inhibited | 1.55E-18 |
| POMC | other | | 1.77E-18 |
| SAICAR | chemical - endogenous mammalian | Activated | 2.01E-18 |
| CYP27A1 | enzyme | | 2.32E-18 |
| BMP4 | growth factor | Activated | 2.65E-18 |
| 3-amino-N-(carboxycarbonyl)-L-alanine | chemical toxicant | Activated | 3.71E-18 |
| vasopressins | biologic drug | Activated | 5.04E-18 |
| tyrosine kinase | group | Activated | 5.09E-18 |
| NRG4 | growth factor | Activated | 5.17E-18 |
| KCNA3 | ion channel | | 5.37E-18 |
| mir-148 | microRNA | Inhibited | 7.19E-18 |
| PPP1R1B | phosphatase | Activated | 8E-18 |
| CAR ligand-CAR-Retinoic acid-RXRα | complex | Activated | 9.11E-18 |
| ZNF703 | other | Inhibited | 1.11E-17 |
| ANO1 | ion channel | Activated | 1.13E-17 |
| SH3GL1 | other | Activated | 1.24E-17 |
| CHMP6 | other | Inhibited | 1.24E-17 |
| panitumumab | biologic drug | Inhibited | 1.24E-17 |
| VPS25 | other | Inhibited | 1.24E-17 |
| quinine | chemical drug | | 1.26E-17 |
| isoproterenol | chemical drug | | 1.51E-17 |
| ZGPAT | transcription regulator | Inhibited | 1.54E-17 |
| MAFK | transcription regulator | Inhibited | 1.74E-17 |
| TRIM65 | other | Inhibited | 1.88E-17 |

| | | | |
|---|---|---|---|
| WNT3A | cytokine | Activated | 2.2E-17 |
| EGFR | kinase | Activated | 2.32E-17 |
| Ap1 | complex | Activated | 2.83E-17 |
| let-7 | microRNA | Inhibited | 2.84E-17 |
| mustard gas | chemical toxicant | Activated | 2.95E-17 |
| Hdac-Rcor-Rest-Sin3a | complex | Inhibited | 3.81E-17 |
| cobalt chloride | chemical reagent | Activated | 3.82E-17 |
| S100A8 | other | Activated | 3.88E-17 |
| lipooligosaccharide | chemical toxicant | Inhibited | 3.91E-17 |
| TADA3 | transcription regulator | | 4.63E-17 |
| SOX2 | transcription regulator | | 4.65E-17 |
| T3-TR-RXR | complex | | 4.87E-17 |
| PD184352 | chemical drug | Inhibited | 5.06E-17 |
| Hdac | group | Inhibited | 5.37E-17 |
| AG 879 | chemical - kinase inhibitor | Inhibited | 6.47E-17 |
| SOX11 | transcription regulator | Inhibited | 6.88E-17 |
| CG | complex | Activated | 7.51E-17 |
| c-Src | group | Activated | 7.54E-17 |
| D-tubocurarine | chemical drug | | 7.85E-17 |
| PI4K | group | | 8.33E-17 |
| ETS | group | Inhibited | 8.39E-17 |
| TP73 | transcription regulator | Inhibited | 1.02E-16 |
| NFAT5 | transcription regulator | | 1.08E-16 |
| lenalidomide | chemical drug | Inhibited | 1.09E-16 |
| EHMT1 | transcription regulator | | 1.18E-16 |
| Rsk | group | Activated | 1.2E-16 |
| Has | group | Activated | 1.34E-16 |
| IFRD1 | other | Inhibited | 1.51E-16 |
| ZFP36 | transcription regulator | Inhibited | 1.72E-16 |
| ETV6 | transcription regulator | Inhibited | 1.72E-16 |
| receptor protein tyrosine kinase | group | Activated | 1.84E-16 |
| KN-62 | chemical - kinase inhibitor | Inhibited | 1.85E-16 |
| denileukin diftitox | biologic drug | | 2.14E-16 |
| octreotide | biologic drug | Inhibited | 2.26E-16 |
| KDM4B | enzyme | Activated | 2.26E-16 |
| NRG (family) | group | Activated | 2.29E-16 |
| barasertib | chemical drug | | 2.46E-16 |
| DICER1 | enzyme | | 2.77E-16 |
| denatonium benzoate | chemical drug | | 2.95E-16 |
| BECN1 | other | Activated | 3.23E-16 |
| CTNNB1 | transcription regulator | Activated | 3.5E-16 |
| PTPRG | phosphatase | Inhibited | 3.64E-16 |
| rolipram | chemical drug | | 3.94E-16 |
| Shc | group | Activated | 4.07E-16 |
| Ige | complex | Activated | 4.09E-16 |
| Gata | group | Activated | 4.17E-16 |
| TAB2/TAB3 | group | Activated | 4.3E-16 |

| | | | |
|---|---|---|---|
| CD74-NRG1 | fusion gene/product | Activated | 4.36E-16 |
| ZNF366 | transcription regulator | | 4.77E-16 |
| SNX5 | transporter | Inhibited | 4.99E-16 |
| PTPRB | phosphatase | Inhibited | 5.08E-16 |
| MYLK2 | kinase | Activated | 5.17E-16 |
| IL6ST | transmembrane receptor | Activated | 5.32E-16 |
| lipoteichoic acid | chemical - endogenous non-mammalian | Activated | 5.64E-16 |
| RING1 | transcription regulator | | 5.91E-16 |
| Lefty | group | Inhibited | 5.93E-16 |
| MEF2 | group | | 6E-16 |
| ITM2B | other | Activated | 6.16E-16 |
| MUC4 | other | Activated | 6.27E-16 |
| DYRK1A | kinase | | 7.02E-16 |
| FAT1 | other | | 7.1E-16 |
| APH1A | peptidase | | 7.34E-16 |
| PSENEN | peptidase | | 7.58E-16 |
| CSF2 | cytokine | Activated | 7.71E-16 |
| SSRP1 | transcription regulator | | 8.66E-16 |
| PTPRK | phosphatase | Inhibited | 9.12E-16 |
| PD 153035 | chemical drug | Inhibited | 9.43E-16 |
| decitabine | chemical drug | Activated | 9.47E-16 |
| ASGR2 | transmembrane receptor | Activated | 9.64E-16 |
| rolapitant | chemical drug | Inhibited | 9.86E-16 |
| aprepitant | chemical drug | Inhibited | 9.86E-16 |
| rhesus theta-defensin 3 | chemical - endogenous mammalian | Inhibited | 1.01E-15 |
| rhesus theta-defensin 2 | chemical - endogenous mammalian | Inhibited | 1.01E-15 |
| HOXA2 | transcription regulator | | 1.02E-15 |
| rhesus theta-defensin 1 | chemical - endogenous mammalian | Inhibited | 1.05E-15 |
| TFF3 | other | Activated | 1.06E-15 |
| LXR ligand-LXR-Retinoic acid-RXRα | complex | Activated | 1.09E-15 |
| AZ-960 | chemical reagent | Inhibited | 1.1E-15 |
| ARIH2 | enzyme | Inhibited | 1.1E-15 |
| L-triiodothyronine | chemical - endogenous mammalian | | 1.15E-15 |
| Ciap | group | | 1.24E-15 |
| APR-246 | chemical drug | | 1.24E-15 |
| MEF2C | transcription regulator | Activated | 1.26E-15 |
| MAP3K3 | kinase | Activated | 1.28E-15 |
| ADAM17 | peptidase | Activated | 1.28E-15 |
| nefiracetam | chemical drug | Activated | 1.56E-15 |
| ERBB2 | kinase | Activated | 1.7E-15 |
| OXSR1 | kinase | Activated | 1.73E-15 |
| TH17 Cytokine | group | Activated | 1.8E-15 |

| | | | |
|---|---|---|---|
| MAC | complex | Activated | 2E-15 |
| PRKD1 | kinase | Activated | 2.02E-15 |
| APH1B | peptidase | | 2.02E-15 |
| ING1 | transcription regulator | | 2.15E-15 |
| PTPRO | phosphatase | Inhibited | 2.26E-15 |
| DNM2 | enzyme | Activated | 2.61E-15 |
| ETS1 | transcription regulator | Inhibited | 2.64E-15 |
| SDC2 | other | Activated | 2.75E-15 |
| tretinoin | chemical - endogenous mammalian | Activated | 2.84E-15 |
| BET | group | Activated | 3.09E-15 |
| HDGF | growth factor | Activated | 3.11E-15 |
| bile acid | chemical - endogenous mammalian | Activated | 3.12E-15 |
| IL11RA | transmembrane receptor | Activated | 3.12E-15 |
| TAC1 | other | Activated | 3.25E-15 |
| 4-hydroxymercuribenzoate | chemical - protease inhibitor | Inhibited | 3.29E-15 |
| CARD9 | other | Activated | 3.53E-15 |
| HAS1 | enzyme | Activated | 3.71E-15 |
| LEMD3 | other | Inhibited | 3.93E-15 |
| PTPRJ | phosphatase | Inhibited | 4.16E-15 |
| platelet activating factor | chemical - endogenous mammalian | Activated | 4.19E-15 |
| CCND1 | transcription regulator | | 4.22E-15 |
| bimatoprost | chemical drug | | 4.38E-15 |
| PTPN22 | phosphatase | Inhibited | 4.53E-15 |
| W7 | chemical reagent | Inhibited | 4.6E-15 |
| IL12 (complex) | complex | Activated | 4.66E-15 |
| PKR dimer | complex | Activated | 4.97E-15 |
| vemurafenib | chemical drug | Inhibited | 5.2E-15 |
| tioconazole | chemical drug | Inhibited | 5.31E-15 |
| tyrphostin AG 1478 | chemical - kinase inhibitor | Inhibited | 5.92E-15 |
| HMOX1 | enzyme | Inhibited | 5.97E-15 |
| SMAD2 | transcription regulator | Activated | 6.06E-15 |
| sulconazole | chemical drug | Inhibited | 6.32E-15 |
| thalidomide | chemical drug | Inhibited | 7.01E-15 |
| formoterol | chemical drug | Inhibited | 7.22E-15 |
| NFYA | transcription regulator | | 7.86E-15 |
| decitabine | chemical drug | Activated | 8.27E-15 |
| NAALADL2 | other | | 8.39E-15 |
| calcitriol | chemical drug | | 8.41E-15 |
| isoniazid | chemical drug | Inhibited | 9.25E-15 |
| PIAS3 | transcription regulator | | 9.29E-15 |
| miR-146a-5p (and other miRNAs w/seed GAGAACU) | mature microRNA | Inhibited | 1.04E-14 |
| HAS2 | enzyme | Activated | 1.12E-14 |

| | | | |
|---|---|---|---|
| CALR | transcription regulator | Activated | 1.32E-14 |
| modafinil | chemical drug | | 1.33E-14 |
| Creb | group | | 1.38E-14 |
| alitretinoin | chemical drug | Activated | 1.44E-14 |
| SFRP | group | Inhibited | 1.54E-14 |
| ammonia | chemical - endogenous mammalian | Activated | 1.56E-14 |
| carbon tetrachloride | chemical toxicant | Activated | 1.69E-14 |
| NEV-801 | chemical drug | Activated | 1.72E-14 |
| imiquimod | chemical drug | Activated | 1.77E-14 |
| SRPK1 | kinase | Activated | 1.82E-14 |
| strontium | chemical - endogenous mammalian | | 1.86E-14 |
| BCAR4 | other | Activated | 1.9E-14 |
| DGCR8 | enzyme | Activated | 1.97E-14 |
| CSF1 | cytokine | Activated | 2.08E-14 |
| bardoxolone methyl | chemical drug | Inhibited | 2.13E-14 |
| mir-10 | microRNA | | 2.15E-14 |
| HES1 | transcription regulator | Activated | 2.23E-14 |
| Glycine Receptor | complex | Activated | 2.24E-14 |
| rosmarinic acid | chemical - endogenous non-mammalian | Inhibited | 2.25E-14 |
| CDC25A | phosphatase | | 2.38E-14 |
| L 778123 | chemical drug | Inhibited | 2.38E-14 |
| dihydrotestosterone | chemical - endogenous mammalian | Activated | 2.41E-14 |
| GGTI-2154 | chemical reagent | Inhibited | 2.54E-14 |
| MITF-p300/CBP | complex | Activated | 2.7E-14 |
| pregnenolone sulfate | chemical - endogenous mammalian | Activated | 2.7E-14 |
| NMDA Receptor | complex | Activated | 2.73E-14 |
| CXCL12 | cytokine | Activated | 2.79E-14 |
| PTPN18 | phosphatase | Inhibited | 2.94E-14 |
| FASLG | cytokine | Activated | 3.06E-14 |
| immune complex | complex | Activated | 3.19E-14 |
| CD74-NRG1 | fusion gene/product | Activated | 3.22E-14 |
| PD 168393 | chemical - kinase inhibitor | Inhibited | 3.26E-14 |
| nadolol | chemical drug | | 3.34E-14 |
| bisindolylmaleimide iv | chemical - kinase inhibitor | Inhibited | 3.39E-14 |
| CAMK2A | kinase | Activated | 3.62E-14 |
| ACP3 | phosphatase | Inhibited | 3.67E-14 |
| PDLIM5 | other | Inhibited | 3.69E-14 |
| etiracetam | chemical drug | Inhibited | 3.76E-14 |
| dopamine | chemical - endogenous mammalian | Activated | 3.87E-14 |
| DAPK1 | kinase | Activated | 3.92E-14 |
| ASGR2 | transmembrane receptor | Activated | 4.01E-14 |
| GPI | enzyme | Activated | 4.01E-14 |

| | | | |
|---|---|---|---|
| Ciap | group | | 4.05E-14 |
| indolocarbazole R-3 | chemical reagent | Activated | 4.05E-14 |
| acitretin | chemical drug | | 4.09E-14 |
| plerixafor | chemical drug | Inhibited | 4.14E-14 |
| MAP3K14 | kinase | Activated | 4.16E-14 |
| CSF2 | cytokine | Activated | 4.23E-14 |
| PSMD10 | transcription regulator | | 4.33E-14 |
| BLNK | other | Activated | 4.34E-14 |
| sulfaphenazole | chemical drug | | 4.38E-14 |
| taxifolin | chemical - endogenous non-mammalian | Inhibited | 4.46E-14 |
| SYNE2 | other | Inhibited | 4.8E-14 |
| bisindolylmaleimide I | chemical - kinase inhibitor | Inhibited | 4.89E-14 |
| NRG3 | growth factor | Activated | 4.95E-14 |
| TNS3 | phosphatase | Activated | 5.02E-14 |
| Crhr | group | Activated | 5.03E-14 |
| CDKN2B | transcription regulator | | 5.11E-14 |
| pentoxifylline | chemical drug | Inhibited | 5.15E-14 |
| escitalopram | chemical drug | Inhibited | 5.27E-14 |
| nitrite | chemical - endogenous mammalian | Inhibited | 5.34E-14 |
| MUC13 | other | Activated | 5.67E-14 |
| SU6656 | chemical toxicant | Inhibited | 5.74E-14 |
| TOP1 | enzyme | Inhibited | 5.75E-14 |
| azelastine | chemical drug | Inhibited | 5.75E-14 |
| FGFR2 | kinase | Activated | 5.84E-14 |
| ITGB1 | transmembrane receptor | Activated | 5.91E-14 |
| EFNA1 | other | Activated | 6.06E-14 |
| uric acid | chemical - endogenous mammalian | Activated | 6.07E-14 |
| Stat5 dimer | complex | | 6.33E-14 |
| ethylene glycol tetraacetic acid | chemical reagent | Inhibited | 7.38E-14 |
| CBX4 | transcription regulator | Activated | 7.38E-14 |
| miR-1258 (miRNAs w/seed GUUAGGA) | mature microRNA | Inhibited | 7.6E-14 |
| Muc4 | other | Activated | 7.64E-14 |
| SMARCA2 | transcription regulator | | 7.75E-14 |
| P2RY2 | G-protein coupled receptor | Activated | 7.82E-14 |
| AHRR | transcription regulator | Inhibited | 8.5E-14 |
| EDN1 | cytokine | Activated | 8.79E-14 |
| SMARCB1 | transcription regulator | | 8.93E-14 |
| PPP2R1A | phosphatase | Inhibited | 9.28E-14 |
| SRC (family) | group | Activated | 9.35E-14 |
| RARG | ligand-dependent nuclear receptor | Inhibited | 1.02E-13 |
| BEX2 | other | Activated | 1.09E-13 |
| andrographolide | chemical drug | Activated | 1.09E-13 |

| | | | |
|---|---|---|---|
| ERBB2 | kinase | Activated | 1.11E-13 |
| Diap | group | Activated | 1.11E-13 |
| THRSP | other | Inhibited | 1.19E-13 |
| LAMTOR2 | other | Activated | 1.2E-13 |
| ID2 | transcription regulator | | 1.27E-13 |
| MAP3K2 | kinase | Activated | 1.31E-13 |
| epinephrine | chemical - endogenous mammalian | | 1.34E-13 |
| isoflurophate | chemical drug | Inhibited | 1.34E-13 |
| (2-(trimethylammonium)ethyl)methanethiosulfonate | chemical reagent | | 1.35E-13 |
| NDUFA13 | enzyme | Inhibited | 1.44E-13 |
| STAT5a/b | group | Inhibited | 1.66E-13 |
| ATP-gamma-S | chemical reagent | Activated | 1.71E-13 |
| CUL1 | enzyme | | 1.95E-13 |
| IL17RA | transmembrane receptor | Activated | 2.03E-13 |
| butaprost | chemical drug | | 2.05E-13 |
| dinoprost | chemical - endogenous mammalian | Activated | 2.15E-13 |
| corticosterone | chemical - endogenous mammalian | | 2.21E-13 |
| ETV4 | transcription regulator | | 2.21E-13 |
| phendimetrazine | chemical drug | Activated | 2.31E-13 |
| calyculin A | chemical toxicant | Activated | 2.34E-13 |
| sulfamethoxazole | chemical drug | Inhibited | 2.36E-13 |
| ALOX12 | enzyme | Activated | 2.45E-13 |
| PELI1 | enzyme | Activated | 2.45E-13 |
| SOX2 | transcription regulator | Activated | 2.46E-13 |
| TNFSF15 | cytokine | Activated | 2.53E-13 |
| UTP | chemical - endogenous mammalian | Activated | 2.57E-13 |
| niclosamide | chemical drug | | 2.67E-13 |
| Rp-cAMPS | chemical - kinase inhibitor | Inhibited | 2.75E-13 |
| BMP7 | growth factor | | 2.89E-13 |
| SPARC | other | Inhibited | 3.2E-13 |
| DRD1/5 | group | Activated | 3.28E-13 |
| CDK4/6 | group | Activated | 3.29E-13 |
| PLX7486 | chemical drug | Inhibited | 3.32E-13 |
| PEBP1 | other | | 3.36E-13 |
| aroclor 1254 | chemical toxicant | | 3.5E-13 |
| chloroform | chemical toxicant | Activated | 3.51E-13 |
| helenalin | chemical - endogenous non-mammalian | Inhibited | 3.95E-13 |
| P2RY4 | G-protein coupled receptor | Activated | 4.27E-13 |
| Crumbs complex | complex | | 4.42E-13 |
| EIF3 | complex | | 4.43E-13 |
| T 0070907 | chemical reagent | | 4.48E-13 |

| | | | |
|---|---|---|---|
| PRKCI | kinase | Activated | 4.5E-13 |
| adaphostin | chemical drug | Activated | 4.66E-13 |
| CASR | G-protein coupled receptor | | 4.67E-13 |
| CBX5 | transcription regulator | | 4.84E-13 |
| Ppp2c | group | Inhibited | 4.85E-13 |
| sesamin | chemical - endogenous non-mammalian | Inhibited | 4.99E-13 |
| PPBP | cytokine | Activated | 5.13E-13 |
| TSLP | cytokine | Activated | 5.27E-13 |
| Ppp1cc | phosphatase | Inhibited | 5.34E-13 |
| SP1 | transcription regulator | | 5.36E-13 |
| TMF1 | transcription regulator | | 5.98E-13 |
| DNA-methyltransferase | group | Activated | 6.33E-13 |
| Focal adhesion kinase | group | | 6.5E-13 |
| ODC1 | enzyme | Activated | 6.53E-13 |
| lycopene | chemical drug | Inhibited | 6.65E-13 |
| CR1 | transmembrane receptor | | 6.99E-13 |
| LRPAP1 | other | Activated | 7.15E-13 |
| MYB | transcription regulator | Activated | 7.31E-13 |
| P2RY11 | G-protein coupled receptor | Activated | 7.48E-13 |
| EPHA2 | kinase | | 7.74E-13 |
| CAMKK2 | kinase | Activated | 7.77E-13 |
| semaxinib | chemical drug | Inhibited | 8.11E-13 |
| GSK2816126 | chemical drug | | 8.24E-13 |
| FOXL2 | transcription regulator | Activated | 8.31E-13 |
| desloratadine | chemical drug | Inhibited | 8.58E-13 |
| MYF6 | transcription regulator | | 8.9E-13 |
| capmatinib | chemical drug | Inhibited | 9.04E-13 |
| AICAR | chemical - endogenous mammalian | | 9.91E-13 |
| arginine | chemical - endogenous mammalian | | 1.11E-12 |
| Wnt | group | Activated | 1.12E-12 |
| RCHY1 | enzyme | | 1.12E-12 |
| CRK | other | Activated | 1.14E-12 |
| IL-1R | group | Activated | 1.3E-12 |
| cytochalasin B | chemical toxicant | Inhibited | 1.3E-12 |
| SOX7 | transcription regulator | Activated | 1.35E-12 |
| DAB2 | other | | 1.38E-12 |
| PF4 | cytokine | Activated | 1.42E-12 |
| ELK1 | transcription regulator | Activated | 1.46E-12 |
| palomid 529 | chemical drug | | 1.52E-12 |
| Cyclin D | group | Activated | 1.62E-12 |
| JNK-IN-8 | chemical - kinase inhibitor | Inhibited | 1.69E-12 |
| leukadherin 1 | chemical reagent | Inhibited | 1.87E-12 |
| MARK2 | kinase | Inhibited | 1.9E-12 |
| COP I | complex | | 1.91E-12 |
| MYOD1 | transcription regulator | | 1.98E-12 |

| | | | |
|---|---|---|---|
| KT5720 | chemical - kinase inhibitor | Inhibited | 2.04E-12 |
| GW3965 | chemical reagent | Inhibited | 2.1E-12 |
| SELPLG | other | | 2.12E-12 |
| aluminum | chemical drug | Inhibited | 2.16E-12 |
| chlorpropamide | chemical drug | | 2.25E-12 |
| acetylcholine | chemical - endogenous mammalian | Activated | 2.26E-12 |
| IL21Rα-IL2Rγ | complex | Activated | 2.29E-12 |
| GNAS | enzyme | Activated | 2.34E-12 |
| RETNLB | other | Activated | 2.51E-12 |
| NTN4 | other | | 2.54E-12 |
| ponatinib | chemical drug | Inhibited | 2.61E-12 |
| HMOX2 | enzyme | Inhibited | 2.63E-12 |
| FOXF1 | transcription regulator | | 2.65E-12 |
| MBL2 | other | | 2.75E-12 |
| FOXF2 | transcription regulator | | 2.81E-12 |
| Pmaip1 | other | Activated | 3.24E-12 |
| bupropion | chemical drug | Activated | 3.41E-12 |
| Rb | group | | 3.82E-12 |
| anacardic acid | chemical - endogenous non-mammalian | | 3.88E-12 |
| mir-33 | microRNA | Inhibited | 3.9E-12 |
| HOTAIR | other | | 3.93E-12 |
| IGFBP7 | transporter | | 3.93E-12 |
| SOD3 | enzyme | Inhibited | 4.17E-12 |
| TBX2 | transcription regulator | Activated | 4.21E-12 |
| OSU-T315 | chemical - kinase inhibitor | Inhibited | 4.21E-12 |
| methylergonovine | chemical drug | Inhibited | 4.23E-12 |
| chrysin | chemical - endogenous non-mammalian | | 4.24E-12 |
| CYP27B1 | enzyme | Activated | 4.24E-12 |
| E2f | group | Activated | 4.3E-12 |
| ABCA1 | transporter | Activated | 4.71E-12 |
| Collagen Alpha1 | group | Activated | 4.73E-12 |
| (E)-2,3',4,5'-tetramethoxystilbene | chemical drug | Inhibited | 4.8E-12 |
| hexamethonium | chemical drug | Inhibited | 4.94E-12 |
| BMS-387032 | chemical drug | Inhibited | 5.05E-12 |
| CDK4/6 | group | Activated | 5.14E-12 |
| IAPP | other | | 5.21E-12 |
| LIMK1 | kinase | Activated | 5.22E-12 |
| RASA1 | transporter | | 5.44E-12 |
| morphiceptin | chemical reagent | Activated | 5.68E-12 |
| PRKD2 | kinase | Activated | 5.83E-12 |
| GATA2 | transcription regulator | Activated | 6.15E-12 |
| caspase | group | Activated | 6.53E-12 |
| dihydroartemisinin | chemical drug | Inhibited | 6.97E-12 |
| l-asparaginase | biologic drug | | 7.36E-12 |

| | | | |
|---|---|---|---|
| VTN | other | Activated | 7.53E-12 |
| VGF | growth factor | Activated | 7.6E-12 |
| WT1 | transcription regulator | | 7.69E-12 |
| decursin | chemical - endogenous non-mammalian | Inhibited | 8.24E-12 |
| CGP 74514A | chemical - kinase inhibitor | | 8.83E-12 |
| MDM4 | transcription regulator | | 9.07E-12 |
| fluoxetine | chemical drug | | 9.07E-12 |
| KLRC4-KLRK1/KLRK1 | transmembrane receptor | | 1.09E-11 |
| farnesyl transferase | complex | Activated | 1.16E-11 |
| cholecalciferol | chemical - endogenous mammalian | Activated | 1.23E-11 |
| PAK6 | kinase | | 1.28E-11 |
| INK4 | group | Activated | 1.28E-11 |
| Pld | group | Activated | 1.29E-11 |
| latrunculin A | chemical toxicant | Activated | 1.42E-11 |
| desmopressin | biologic drug | Activated | 1.5E-11 |
| TOPORS | enzyme | Inhibited | 1.5E-11 |
| beta-escin | chemical drug | | 1.52E-11 |
| MALT1 | peptidase | Activated | 1.58E-11 |
| RBM8A | other | Activated | 1.72E-11 |
| CXCL5 | cytokine | Activated | 1.76E-11 |
| ajoene | chemical - endogenous non-mammalian | Activated | 1.98E-11 |
| DUSP2 | phosphatase | | 2.13E-11 |
| MIRLET7 | group | Inhibited | 2.58E-11 |
| harmol | chemical - endogenous non-mammalian | Activated | 2.69E-11 |
| PBK | kinase | Activated | 2.82E-11 |
| Eotaxin | group | Activated | 2.94E-11 |
| CDKN1C | other | Inhibited | 3.18E-11 |
| LILRB4 | other | Inhibited | 3.38E-11 |
| LDL-cholesterol | complex | Activated | 3.56E-11 |
| EPS8 | peptidase | | 3.74E-11 |
| HOXA7 | transcription regulator | | 4.29E-11 |
| Salmonella enterica serotype abortus equi lipopolysaccharide | chemical toxicant | Activated | 4.39E-11 |
| CDKN2C | transcription regulator | Inhibited | 4.41E-11 |
| SPDEF | transcription regulator | Inhibited | 5.33E-11 |
| cephaloridine | chemical drug | Activated | 5.65E-11 |
| STAT3/5 | group | | 5.82E-11 |
| ADIPOR1 | transmembrane receptor | | 5.86E-11 |
| CBFA2T3 | transcription regulator | Inhibited | 6.17E-11 |
| ETV1 | transcription regulator | | 6.23E-11 |
| ZNF217 | transcription regulator | | 6.62E-11 |
| MITF | transcription regulator | Activated | 6.8E-11 |
| ginkgolide B | chemical drug | | 7.34E-11 |

| | | | |
|---|---|---|---|
| CAMK2D | kinase | Activated | 8.38E-11 |
| 6-cyano-7-nitroquinoxaline-2,3-dione | chemical reagent | | 8.53E-11 |
| lysophosphatidylinositol | chemical - endogenous mammalian | Activated | 8.78E-11 |
| BIRC2 | enzyme | | 9E-11 |
| SORBS3 | other | | 9.27E-11 |
| BAY 61-3606 | chemical - kinase inhibitor | Inhibited | 9.37E-11 |
| silicon phthalocyanine | chemical drug | Inhibited | 9.52E-11 |
| GADD45G | other | Activated | 1.03E-10 |
| porphyrin | chemical - other | | 1.08E-10 |
| fumagillin | chemical - endogenous non-mammalian | | 1.19E-10 |
| dexmedetomidine | chemical drug | | 1.31E-10 |
| DOV-102,677 | chemical drug | | 1.44E-10 |
| D-methylphenidate | chemical drug | | 1.44E-10 |
| SH2B2 | other | | 1.46E-10 |
| FAIM | other | | 1.49E-10 |
| IH636 grape seed proanthocyanidin extract | chemical drug | | 1.61E-10 |
| doramapimod | chemical - kinase inhibitor | Inhibited | 1.62E-10 |
| neomycin | chemical drug | Activated | 1.65E-10 |
| NOTCH4 | transcription regulator | Activated | 1.66E-10 |
| mir-29 | microRNA | | 1.66E-10 |
| XCT790 | chemical reagent | Inhibited | 1.68E-10 |
| ditiocarb | chemical drug | | 1.75E-10 |
| l-asparaginase | biologic drug | Inhibited | 1.89E-10 |
| SB203580 | chemical - kinase inhibitor | Inhibited | 2.23E-10 |
| salsalate | chemical drug | Inhibited | 2.24E-10 |
| NFX1 | transcription regulator | | 2.25E-10 |
| SNAI1 | transcription regulator | Activated | 2.38E-10 |
| HNRNPD | transcription regulator | | 2.69E-10 |
| PINX1 | other | | 3.05E-10 |
| hexachlorobenzene | chemical toxicant | Activated | 3.08E-10 |
| 2-(4-amino-1-isopropyl-1H-pyrazolo[3,4-d]pyrimidin-3-yl)-1H-indol-5-ol | chemical reagent | Inhibited | 3.28E-10 |
| ridaforolimus | chemical drug | | 3.66E-10 |
| pexmetinib | chemical drug | Inhibited | 4.1E-10 |
| SPI1 | transcription regulator | | 4.12E-10 |
| PDGF-DD | complex | | 4.2E-10 |
| isorhamnetin | chemical - endogenous non-mammalian | Inhibited | 4.27E-10 |
| FLI1 | transcription regulator | | 4.48E-10 |
| CCM2 | other | Inhibited | 4.58E-10 |
| TERT | enzyme | | 4.62E-10 |
| IGFBP6 | other | Activated | 4.65E-10 |

| | | | |
|---|---|---|---|
| pyrazole | chemical - endogenous non-mammalian | Activated | 4.87E-10 |
| MGAT5 | enzyme | | 5.14E-10 |
| PD 180970 | chemical - kinase inhibitor | Inhibited | 5.39E-10 |
| CDC34 | enzyme | Inhibited | 5.52E-10 |
| AGO2 | translation regulator | Activated | 5.91E-10 |
| AGN194204 | chemical drug | | 6.04E-10 |
| RGS5 | enzyme | | 6.31E-10 |
| FBLN5 | other | Activated | 7.03E-10 |
| EIF4EBP2 | translation regulator | | 7.14E-10 |
| AGRP | other | Activated | 7.24E-10 |
| lansoprazole | chemical drug | Inhibited | 7.61E-10 |
| mimosine | chemical drug | Inhibited | 7.65E-10 |
| DNMT3A | enzyme | | 7.65E-10 |
| CDKN3 | phosphatase | Inhibited | 8.05E-10 |
| SRF-ELK1 | complex | Activated | 8.07E-10 |
| DOCK4 | other | | 8.62E-10 |
| terbutaline | chemical drug | Activated | 9.07E-10 |
| JQ1 | chemical reagent | Inhibited | 9.51E-10 |
| lisofylline | chemical drug | Inhibited | 9.67E-10 |
| PTPN14 | phosphatase | Inhibited | 1.02E-09 |
| FRMD6 | other | Inhibited | 1.08E-09 |
| astemizole | chemical drug | Inhibited | 1.28E-09 |
| TAB2/TAB3 | group | Activated | 1.34E-09 |
| ARHGEF11 | other | Activated | 1.39E-09 |
| dimethyl sulfone | chemical drug | | 1.46E-09 |
| pyrilamine | chemical drug | | 1.57E-09 |
| UBE2L3-KRAS | fusion gene/product | | 1.63E-09 |
| CIRBP | translation regulator | Activated | 1.74E-09 |
| midazolam | chemical drug | Inhibited | 1.78E-09 |
| CBX7 | other | | 1.95E-09 |
| TFPI2 | other | Inhibited | 2.17E-09 |
| PSAP | enzyme | | 2.35E-09 |
| dATP | chemical - endogenous mammalian | Inhibited | 2.38E-09 |
| ULBP1 | transmembrane receptor | | 2.53E-09 |
| azathioprine | chemical drug | | 2.69E-09 |
| harmaline | chemical drug | Activated | 2.73E-09 |
| LAMA3 | other | Activated | 3.14E-09 |
| 20s proteasome | complex | | 3.43E-09 |
| fenoterol | chemical drug | Activated | 3.51E-09 |
| HPRT1 | enzyme | | 3.64E-09 |
| HINT1 | enzyme | Inhibited | 3.72E-09 |
| bergamottin | chemical - endogenous non-mammalian | Inhibited | 4.14E-09 |
| DNA-methyltransferase | group | Inhibited | 4.26E-09 |
| flutamide | chemical drug | | 4.33E-09 |
| ETV6 | transcription regulator | | 4.35E-09 |

| | | | |
|---|---|---|---|
| HDAC2 | transcription regulator | | 4.76E-09 |
| NONO | transcription regulator | Inhibited | 4.87E-09 |
| TRIP12 | enzyme | Activated | 5.11E-09 |
| Irgm1 | other | Inhibited | 5.17E-09 |
| FOXF2 | transcription regulator | | 5.22E-09 |
| mevastatin | chemical drug | | 5.59E-09 |
| Par | group | Activated | 5.7E-09 |
| TBX2 | transcription regulator | Activated | 5.7E-09 |
| mir-132 | microRNA | Activated | 5.7E-09 |
| NEO1 | transcription regulator | | 5.83E-09 |
| DNMT3L | transcription regulator | Inhibited | 6.53E-09 |
| acyline | biologic drug | Activated | 7.14E-09 |
| REL | transcription regulator | Activated | 7.28E-09 |
| Alpha 1 antitrypsin | group | Inhibited | 9.8E-09 |
| GNAT1 | enzyme | Inhibited | 1.09E-08 |
| GLPG0187 | chemical drug | Activated | 1.16E-08 |
| saracatinib | chemical drug | Inhibited | 1.17E-08 |
| tetomilast | chemical drug | | 1.18E-08 |
| arofylline | chemical drug | | 1.18E-08 |
| L 869298 | chemical drug | | 1.18E-08 |
| CAND1 | transcription regulator | Activated | 1.19E-08 |
| TRERF1 | transcription regulator | | 1.31E-08 |
| L-826,141 | chemical drug | | 1.37E-08 |
| FOXF1 | transcription regulator | Activated | 1.47E-08 |
| flavone | chemical - endogenous non-mammalian | | 1.53E-08 |
| Fe3+ | chemical - endogenous mammalian | Activated | 1.82E-08 |
| CCND1 | transcription regulator | Activated | 1.95E-08 |
| TCF7L2 | transcription regulator | | 1.99E-08 |
| NCX-4040 | chemical drug | | 2.03E-08 |
| PSMB11 | peptidase | Inhibited | 2.19E-08 |
| CUL4A | other | | 2.23E-08 |
| Cyclin D1/cdk4 | complex | | 2.25E-08 |
| Rb | group | Inhibited | 2.27E-08 |
| SELP | transmembrane receptor | Activated | 2.67E-08 |
| NUPR1 | transcription regulator | | 0.000000027 |
| CDC7 | kinase | Inhibited | 2.85E-08 |
| E2F6 | transcription regulator | Inhibited | 3.11E-08 |
| calcitriol | chemical drug | Inhibited | 3.73E-08 |
| CAK | complex | | 4.12E-08 |
| (R)-limonene | chemical toxicant | Activated | 4.97E-08 |
| ZIC2 | transcription regulator | Inhibited | 5.28E-08 |
| norethynodrel | chemical toxicant | | 6.55E-08 |
| UBTD1 | other | Inhibited | 6.77E-08 |
| SWI-SNF | complex | Activated | 0.000000068 |
| USO1 | other | | 6.93E-08 |
| CD59 | other | | 7.08E-08 |

| | | | |
|---|---|---|---|
| ITPKB | kinase | Inhibited | 7.16E-08 |
| Mcpt4 | peptidase | Inhibited | 7.28E-08 |
| SMARCE1 | transcription regulator | | 7.43E-08 |
| cannabinol | chemical - endogenous mammalian | Inhibited | 7.48E-08 |
| ergocalciferol | chemical - endogenous mammalian | | 8.26E-08 |
| Brd4 | kinase | Activated | 9.47E-08 |
| endothelin receptor | group | | 9.63E-08 |
| LATS1 | kinase | Inhibited | 9.78E-08 |
| decitabine | chemical drug | Activated | 0.0000001 |
| 4-hydroxymercuribenzoate | chemical - protease inhibitor | Inhibited | 0.000000104 |
| MITF | transcription regulator | Activated | 0.000000109 |
| Salmonella enterica serotype abortus equi lipopolysaccharide | chemical toxicant | Activated | 0.000000123 |
| BOP1 | other | | 0.000000124 |
| inecalcitol | chemical drug | | 0.00000013 |
| DNMT3A | enzyme | Inhibited | 0.000000138 |
| TAF4 | transcription regulator | Activated | 0.000000168 |
| Vegf | group | Activated | 0.000000181 |
| TP63 | transcription regulator | Activated | 0.000000218 |
| PANX3 | other | Inhibited | 0.000000233 |
| Gamma tubulin | group | Inhibited | 0.000000234 |
| TFDP2 | transcription regulator | Activated | 0.000000234 |
| repotrectinib | chemical drug | | 0.000000239 |
| UHRF1 | transcription regulator | | 0.000000258 |
| MAP2 | other | Activated | 0.000000259 |
| ITI-078 | chemical reagent | Activated | 0.000000269 |
| tegaserod | chemical drug | Activated | 0.000000277 |
| DNASE2 | enzyme | Inhibited | 0.000000299 |
| MEF2 | group | | 0.000000302 |
| bivalirudin | biologic drug | Inhibited | 0.000000368 |
| INHA | growth factor | | 0.000000373 |
| Integrin alpha 6 beta 1 | complex | | 0.000000373 |
| ASCL1 | transcription regulator | | 0.000000374 |
| epothilone B | chemical drug | | 0.0000004 |
| EHF | transcription regulator | Activated | 0.000000433 |
| calcimycin/tetradecanoylphorbol acetate | chemical reagent | Activated | 0.000000454 |
| diosmin | chemical drug | | 0.000000463 |
| TGFBR2 | kinase | | 0.000000475 |
| ethyl ether | chemical reagent | Activated | 0.000000495 |
| WNK1 | kinase | Activated | 0.000000497 |
| NEUROG2 | transcription regulator | Inhibited | 0.000000522 |
| PF-04957325 | chemical reagent | Activated | 0.000000534 |
| FAS | transmembrane receptor | | 0.000000537 |

| | | | |
|---|---|---|---|
| STING1 | other | Activated | 0.00000061 |
| RACGAP1 | transporter | | 0.000000623 |
| RNF123 | other | | 0.00000068 |
| VCAN | other | | 0.000000734 |
| MIR99A-LET7C-MIR125B2 | group | Inhibited | 0.000000766 |
| MIR100-LET7A2-MIR125B1 | group | Inhibited | 0.000000766 |
| THZ1 | chemical - kinase inhibitor | | 0.00000079 |
| EIF3H | other | Activated | 0.000000804 |
| CTSD | peptidase | | 0.00000082 |
| TEAD2 | transcription regulator | | 0.000000905 |
| 4-tert-octylphenol | chemical toxicant | Activated | 0.000000907 |
| MRTFB | transcription regulator | Activated | 0.00000094 |
| bis(4-hydroxycinnamoyl)methane | chemical - endogenous non-mammalian | | 0.00000105 |
| VGLL4 | other | Activated | 0.00000107 |
| CREB1 | transcription regulator | Activated | 0.00000108 |
| estrogen | chemical drug | Activated | 0.00000112 |
| RASA2 | other | Inhibited | 0.00000114 |
| NKX2-5 | transcription regulator | | 0.00000116 |
| dihydrotestosterone | chemical - endogenous mammalian | Activated | 0.00000136 |
| SOX2 | transcription regulator | | 0.00000139 |
| PTGES3 | enzyme | | 0.00000145 |
| Npm | group | | 0.00000169 |
| NFkB1-CRel | complex | Activated | 0.00000172 |
| FBXL12 | other | Activated | 0.00000174 |
| fulvestrant | chemical drug | Inhibited | 0.00000192 |
| SCML2 | transcription regulator | Activated | 0.00000197 |
| B4GALNT1 | enzyme | | 0.00000205 |
| NFIC | transcription regulator | Activated | 0.00000211 |
| ifosfamide | chemical drug | | 0.00000213 |
| lanthanum chloride | chemical reagent | Activated | 0.0000022 |
| 17-alpha-ethinylestradiol | chemical drug | | 0.0000023 |
| deferoxamine | chemical drug | Activated | 0.0000023 |
| HOXA4 | transcription regulator | Inhibited | 0.0000023 |
| CYP27B1 | enzyme | | 0.00000242 |
| cation | chemical - other | Activated | 0.00000247 |
| E. coli B4 lipopolysaccharide | chemical toxicant | Activated | 0.00000248 |
| zearalenone | chemical toxicant | | 0.0000025 |
| GATA2 | transcription regulator | | 0.00000274 |
| MGMT | enzyme | | 0.00000275 |
| galeterone | chemical drug | | 0.00000298 |
| abiraterone acetate | chemical drug | | 0.00000298 |
| Integrin alpha V beta 3 | complex | | 0.000003 |
| LIMD1 | transcription regulator | Activated | 0.00000304 |
| YAP1 | transcription regulator | Activated | 0.00000309 |

| | | | |
|---|---|---|---|
| CHUK | kinase | Activated | 0.00000329 |
| Ephb dimer | complex | | 0.00000329 |
| FGF3 | growth factor | Activated | 0.00000332 |
| E2F2 | transcription regulator | | 0.0000036 |
| E2F3 | transcription regulator | | 0.00000382 |
| ABCB4 | transporter | Inhibited | 0.00000395 |
| PURA | transcription regulator | Activated | 0.00000416 |
| PRDM1 | transcription regulator | | 0.00000432 |
| TRIM65 | other | Inhibited | 0.00000451 |
| TBX2 | transcription regulator | Activated | 0.00000467 |
| mir-194 | microRNA | Inhibited | 0.00000468 |
| CDKN1A | kinase | Inhibited | 0.00000498 |
| THAP12 | transcription regulator | Activated | 0.0000051 |
| SMARCB1 | transcription regulator | | 0.00000531 |
| let-7 | microRNA | Inhibited | 0.00000551 |
| WNT5A | cytokine | | 0.00000574 |
| BI 894999 | chemical drug | Inhibited | 0.00000574 |
| cinacalcet | chemical drug | Activated | 0.00000576 |
| MAD1L1 | other | | 0.00000576 |
| BRD4 | kinase | Activated | 0.00000606 |
| AIMP2 | other | Activated | 0.00000646 |
| TEAD | group | Activated | 0.00000663 |
| LIF | cytokine | | 0.00000681 |
| CASR | G-protein coupled receptor | Activated | 0.0000074 |
| DDB1 | other | | 0.00000744 |
| CDKN2A | transcription regulator | Inhibited | 0.00000749 |
| 6-aminopyrazolopyrimidine derivative compound II | chemical - kinase inhibitor | Inhibited | 0.00000793 |
| PTGER4 | G-protein coupled receptor | | 0.00000812 |
| GATA6 | transcription regulator | | 0.00000817 |
| 3-hydroxykynurenine | chemical - endogenous mammalian | | 0.00000914 |
| NFAT5 | transcription regulator | Activated | 0.00000925 |
| CR1L | other | Inhibited | 0.00000945 |
| I kappa b kinase | complex | Inhibited | 0.00000985 |
| Cmtm2a | transcription regulator | | 0.000011 |
| DNMT3B | enzyme | Inhibited | 0.0000114 |
| fluticasone propionate | chemical drug | | 0.0000126 |
| ACTB | other | Activated | 0.0000126 |
| ethyl ether | chemical reagent | Activated | 0.0000127 |
| [Ac-His1,D-Phe2,Lys15,Arg16,Leu27]VIP-(3-7)-GRF-(8-27) | chemical reagent | Inhibited | 0.0000149 |
| MEF2D | transcription regulator | | 0.0000151 |
| NOTCH4 | transcription regulator | Activated | 0.0000152 |
| CCNE1 | transcription regulator | | 0.0000154 |
| SMARCA4 | transcription regulator | Activated | 0.0000165 |

| | | | |
|---|---|---|---|
| L-carnitine | chemical - endogenous mammalian | Inhibited | 0.0000173 |
| NC9 | chemical reagent | Inhibited | 0.0000175 |
| Pam3-Cys-Ser-Lys4 | chemical reagent | Activated | 0.0000179 |
| HTR2A | G-protein coupled receptor | Activated | 0.0000186 |
| TREM1 | transmembrane receptor | | 0.0000194 |
| ASH2L | transcription regulator | | 0.0000194 |
| cdc25C phosphatase (211-221) | biologic drug | | 0.0000202 |
| 4-tert-octylphenol | chemical toxicant | Activated | 0.0000204 |
| Ifnar | group | Activated | 0.0000206 |
| F2R | G-protein coupled receptor | | 0.0000206 |
| HLA-E | transmembrane receptor | | 0.0000212 |
| DEGS1 | enzyme | | 0.0000221 |
| CREB1 | transcription regulator | Activated | 0.0000233 |
| thioacetamide | chemical toxicant | Activated | 0.0000234 |
| PCSK5 | peptidase | Activated | 0.0000237 |
| miR-34a-5p (and other miRNAs w/seed GGCAGUG) | mature microRNA | Inhibited | 0.0000244 |
| SPTAN1 | other | | 0.0000251 |
| PLK2 | kinase | | 0.000028 |
| NR4A1 | ligand-dependent nuclear receptor | | 0.0000299 |
| cholecalciferol | chemical - endogenous mammalian | | 0.0000311 |
| NUMB/NUMBL | group | | 0.0000314 |
| GATA4 | transcription regulator | | 0.0000316 |
| STAT5B | transcription regulator | | 0.0000317 |
| CBFA2T3 | transcription regulator | | 0.0000325 |
| lysophosphatidylinositol | chemical - endogenous mammalian | Activated | 0.0000397 |
| IHH | enzyme | | 0.0000399 |
| dimethylnitrosamine | chemical toxicant | Activated | 0.0000408 |
| mir-223 | microRNA | | 0.0000469 |
| FADD | other | | 0.0000475 |
| 1810019D21Rik | other | | 0.0000491 |
| BNIP3L | other | | 0.0000491 |
| GATA3 | transcription regulator | | 0.0000494 |
| terfenadine | chemical drug | | 0.0000541 |
| PP2/AG1879 tyrosine kinase inhibitor | chemical - kinase inhibitor | Inhibited | 0.0000543 |
| PTH | other | Activated | 0.0000591 |
| azelastine | chemical drug | | 0.0000677 |
| KDM3A | transcription regulator | Activated | 0.0000763 |
| ACKR2 | G-protein coupled receptor | Inhibited | 0.0000763 |
| HTR2C | G-protein coupled receptor | | 0.0000801 |
| GCSAM | other | | 0.000084 |

| | | | |
|---|---|---|---|
| Mt3 | other | Activated | 0.000084 |
| GATA1 | transcription regulator | | 0.0000842 |
| NSOP00313 | chemical reagent | | 0.0000871 |
| tributyltin | chemical reagent | | 0.0000908 |
| WBP2 | transcription regulator | | 0.000102 |
| FOSB | transcription regulator | | 0.000104 |
| conteltinib | chemical drug | | 0.000108 |
| Irgm1 | other | Inhibited | 0.000112 |
| EP400 | other | Activated | 0.000118 |
| IKZF2 | transcription regulator | Inhibited | 0.000118 |
| TRIM24 | transcription regulator | Inhibited | 0.000118 |
| UTS2 | other | | 0.000118 |
| tryptase | group | | 0.000121 |
| PLK4 | kinase | Activated | 0.000121 |
| UBE2L6 | enzyme | | 0.000121 |
| vancomycin | biologic drug | Activated | 0.000131 |
| HDAC11 | transcription regulator | Inhibited | 0.000132 |
| CYP1B1 | enzyme | | 0.000134 |
| cerivastatin | chemical drug | | 0.000134 |
| laminaran | chemical drug | Activated | 0.000142 |
| dimethyl sulfoxide | chemical drug | Activated | 0.000146 |
| TAF4B | transcription regulator | Activated | 0.000162 |
| FGFR2 | kinase | | 0.000163 |
| halofuginone | chemical drug | Inhibited | 0.000179 |
| Type I BMP receptor | group | Inhibited | 0.000187 |
| KDM1A | enzyme | | 0.00019 |
| RBL1 | transcription regulator | Inhibited | 0.000191 |
| PD173074 | chemical reagent | Activated | 0.000202 |
| mir-19 | microRNA | Inhibited | 0.000218 |
| B4GALNT2 | enzyme | | 0.000229 |
| C5 | other | | 0.000244 |
| JQ1 | chemical reagent | Inhibited | 0.000253 |
| histone deacetylase | complex | | 0.000275 |
| PCBP2 | other | Inhibited | 0.000276 |
| sodium dodecyl sulfate | chemical drug | | 0.000276 |
| buprenorphine | chemical drug | | 0.0003 |
| TNFAIP6 | other | | 0.000304 |
| isotretinoin | biologic drug | | 0.000313 |
| IRAK4 | kinase | Activated | 0.000316 |
| GLIPR2 | other | | 0.000337 |
| ADAMTS18 | peptidase | Activated | 0.000342 |
| KRT17 | other | | 0.000342 |
| CRKL | kinase | Activated | 0.000352 |
| ZFP36 | transcription regulator | Inhibited | 0.000354 |
| SMPD1 | enzyme | | 0.000364 |
| CBX7 | other | | 0.000364 |
| Alpha Actinin | group | | 0.000364 |
| SAV1 | other | | 0.000364 |

| | | | |
|---|---|---|---|
| hydroxyurea | chemical drug | Activated | 0.000372 |
| TAF4 | transcription regulator | | 0.000372 |
| SEMA7A | transmembrane receptor | | 0.000396 |
| 8,9-epoxyeicosatrienoic acid | chemical - endogenous mammalian | | 0.000406 |
| deferasirox | chemical drug | | 0.000421 |
| ritonavir | chemical drug | | 0.000421 |
| CIP2A | other | | 0.000425 |
| lipid A | chemical toxicant | Activated | 0.000449 |
| EFNA3 | kinase | | 0.000449 |
| Rock | group | | 0.000449 |
| ADAM12 | peptidase | | 0.00045 |
| belizatinib | chemical drug | Inhibited | 0.00045 |
| JAG1 | growth factor | Activated | 0.00049 |
| carboplatin | chemical drug | Activated | 0.00049 |
| bromodeoxyuridine | chemical drug | Activated | 0.00049 |
| Go6983 | chemical - kinase inhibitor | | 0.000493 |
| TNK1 | kinase | Activated | 0.000493 |
| oxytetracycline | chemical drug | | 0.000493 |
| imipramine blue | chemical drug | | 0.000514 |
| ENG | transmembrane receptor | | 0.000514 |
| GDF9 | growth factor | | 0.000514 |
| EFNA5 | kinase | | 0.00053 |
| ARHGAP21 | other | Activated | 0.00053 |
| TCF12 | transcription regulator | | 0.000549 |
| MIR143-145a | group | | 0.000565 |
| DSCAML1 | other | Activated | 0.000614 |
| SOX7 | transcription regulator | | 0.000622 |
| GMP | chemical - endogenous mammalian | | 0.000622 |
| Ro 25-6760 | chemical toxicant | | 0.000623 |
| AMH | growth factor | | 0.000643 |
| LECT2 | other | Activated | 0.000652 |
| 6-mercaptopurine | chemical drug | Inhibited | 0.000652 |
| TET2 | enzyme | | 0.000658 |
| FZD9 | G-protein coupled receptor | Activated | 0.000673 |
| latrunculin B | chemical reagent | Inhibited | 0.000673 |
| 10E,12Z-octadecadienoic acid | chemical - endogenous mammalian | | 0.000724 |
| SERPINB1 | other | | 0.000729 |
| PLK2 | kinase | Activated | 0.000805 |
| immethridine | chemical reagent | Inhibited | 0.000805 |
| 8-epi-prostaglandin F2alpha | chemical - endogenous mammalian | | 0.000814 |
| ferric chloride | chemical toxicant | | 0.000814 |
| AGA | enzyme | | 0.000814 |
| E2F2 | transcription regulator | Activated | 0.000848 |
| stallimycin | biologic drug | Activated | 0.000848 |

| | | | |
|---|---|---|---|
| NUMB | other | | 0.000899 |
| ITGA1 | other | | 0.000899 |
| trypsin | group | | 0.000989 |
| ryanodine | chemical - endogenous non-mammalian | | 0.000989 |
| palbociclib | chemical drug | | 0.000989 |
| PLK4 | kinase | Activated | 0.000997 |
| SFN | other | | 0.000997 |
| semaxinib | chemical drug | | 0.000997 |
| signal peptidase | complex | | 0.00106 |
| 3-o-acetyl-beta-boswellic acid | chemical reagent | | 0.00106 |
| TREX1 | enzyme | Inhibited | 0.00118 |
| creatine kinase | group | | 0.00118 |
| CDH11 | other | Activated | 0.00122 |
| ROCK1 | kinase | Activated | 0.00122 |
| SAA1 | transporter | | 0.00125 |
| vatalanib | chemical drug | Inhibited | 0.00143 |
| trametinib | chemical drug | Activated | 0.00143 |
| PDGFRB | kinase | | 0.00143 |
| MED16 | transcription regulator | | 0.00143 |
| Cdkn1c | other | Inhibited | 0.00143 |
| oxymorphone | chemical drug | | 0.00143 |
| EHMT1 | transcription regulator | Activated | 0.00144 |
| CEP-37440 | chemical drug | | 0.00147 |
| CD5L | transmembrane receptor | | 0.00147 |
| ramipril | chemical drug | Inhibited | 0.00148 |
| CARD9 | other | | 0.00148 |
| NTRK2 | kinase | Activated | 0.00149 |
| tetralinoleoyl cardiolipin | chemical - endogenous mammalian | Inhibited | 0.00149 |
| ADH5 | enzyme | Inhibited | 0.00149 |
| Mff | other | Inhibited | 0.00149 |
| VGLL3 | other | | 0.00151 |
| emricasan | chemical drug | | 0.00157 |
| XCT790 | chemical reagent | | 0.00157 |
| 19,20-epoxydocosapentaenoic acid | chemical - endogenous mammalian | | 0.00157 |
| ACTA2 | other | | 0.00157 |
| WAC | other | | 0.00157 |
| CLEC12A | other | | 0.00157 |
| E2F7 | transcription regulator | | 0.00157 |
| miR-27a-3p (and other miRNAs w/seed UCACAGU) | mature microRNA | | 0.00171 |
| mir-145 | microRNA | | 0.00171 |
| IHH | enzyme | | 0.00171 |

| | | | |
|---|---|---|---|
| NEUROG1 | transcription regulator | Inhibited | 0.00176 |
| PPP3R1 | phosphatase | | 0.00176 |
| dihydromorphine | chemical drug | | 0.00176 |
| ETV4 | transcription regulator | | 0.00179 |
| butaprost | chemical drug | | 0.00191 |
| ST8SIA1 | enzyme | | 0.00191 |
| EFNA2 | kinase | | 0.00194 |
| COX10 | enzyme | Inhibited | 0.00194 |
| 4-(heptyloxy)phenol | chemical reagent | | 0.00197 |
| 4-(octyloxy)phenol | chemical reagent | | 0.00197 |
| ACTC1 | enzyme | | 0.00197 |
| AGN 190299 | chemical drug | | 0.00197 |
| tyrphostin AG 1295 | chemical - kinase inhibitor | | 0.00197 |
| ELF2 | transcription regulator | | 0.00197 |
| LAMA1 | other | | 0.00197 |
| malolactomycin D | chemical - endogenous non-mammalian | | 0.00197 |
| CTS-1027 | chemical drug | | 0.00197 |
| [D-Ala2,N-Me-Phe4,Gly5-ol]-Enkephalin | chemical reagent | | 0.002 |
| Collagen Alpha1 | group | | 0.002 |
| verteporfin | chemical drug | Inhibited | 0.002 |
| NKX2-5 | transcription regulator | | 0.00213 |
| moclobemide | chemical drug | | 0.00219 |
| EFNA4 | kinase | | 0.00229 |
| IDH1 | enzyme | | 0.00238 |
| PARPBP | other | | 0.00238 |
| DYRK1A | kinase | Inhibited | 0.00238 |
| azoxymethane | chemical toxicant | | 0.00253 |
| TAK-901 | chemical drug | | 0.00253 |
| laquinimod | chemical drug | Activated | 0.00263 |
| saracatinib | chemical drug | | 0.00266 |
| MIR320 | group | | 0.00266 |
| SERPINH1 | other | | 0.00266 |
| miR-148a-3p (and other miRNAs w/seed CAGUGCA) | mature microRNA | Inhibited | 0.0027 |
| NMU | other | | 0.0027 |
| GRK2 | kinase | | 0.0027 |
| HOXA4 | transcription regulator | | 0.0027 |
| RYBP | transcription regulator | | 0.0027 |
| TFPI2 | other | Activated | 0.00294 |
| Mitochondrial complex 1 | complex | Inhibited | 0.00294 |
| tioconazole | chemical drug | | 0.00333 |
| etiracetam | chemical drug | | 0.00333 |
| laquinimod | chemical drug | Activated | 0.00352 |
| TNFRSF11B | transmembrane receptor | | 0.00355 |
| triclosan | chemical drug | | 0.00355 |

| | | | |
|---|---|---|---|
| NUP98-KDM5A | fusion gene/product | Inhibited | 0.00355 |
| NUP98-NSD1 | fusion gene/product | Inhibited | 0.00355 |
| RUNX1-RUNX1T1 | fusion gene/product | | 0.00355 |
| CDK9 | kinase | | 0.00357 |
| ISGF3 | complex | | 0.00412 |
| HDAC11 | transcription regulator | | 0.00412 |
| RNF40 | enzyme | | 0.00412 |
| 15-epi-lipoxin A4 | chemical - endogenous mammalian | | 0.00412 |
| Cd64 | group | | 0.00412 |
| KNG1 | other | | 0.00431 |
| PCYT1A | enzyme | | 0.00431 |
| benz[a]anthracene | chemical toxicant | | 0.00435 |
| 101.10 peptide | chemical reagent | Inhibited | 0.00457 |
| morin | chemical - endogenous non-mammalian | | 0.00457 |
| creatine | chemical - endogenous mammalian | | 0.00457 |
| Gm12602 | other | Inhibited | 0.00457 |
| ATG13 | other | | 0.00457 |
| meperidine | chemical drug | | 0.00457 |
| desmopressin | biologic drug | Activated | 0.00534 |
| CYP1A1 | enzyme | | 0.00559 |
| BCO1 | enzyme | | 0.00572 |
| CFH | other | | 0.00572 |
| 18-alpha-glycyrrhetinic acid | chemical - endogenous non-mammalian | | 0.00572 |
| bis(4-hydroxycinnamoyl)methane | chemical - endogenous non-mammalian | | 0.00572 |
| 15-LOX | group | | 0.00572 |
| AMOTL1 | other | | 0.00572 |
| AMOT | other | | 0.00572 |
| ZHX2 | transcription regulator | | 0.00572 |
| SLC3A2 | transporter | | 0.00572 |
| vinyl carbamate | chemical toxicant | | 0.00572 |
| adapalene | chemical drug | | 0.00572 |
| ENPP7 | enzyme | | 0.00572 |
| Cacnb1 | ion channel | | 0.00572 |
| LPXN | transcription regulator | | 0.00572 |
| CXCL11 | cytokine | | 0.00572 |
| etodolac | chemical drug | | 0.00572 |
| arzoxifene | chemical drug | | 0.00572 |
| PEPCK | group | | 0.00572 |
| HMG20B | transcription regulator | | 0.00572 |
| TRIM41 | other | | 0.00572 |
| Diap | group | | 0.00572 |
| CACNA2D1 | ion channel | | 0.00572 |
| ST6GALNAC2 | enzyme | | 0.00572 |

| | | | |
|---|---|---|---|
| RBBP7 | transcription regulator | | 0.00572 |
| 6-amino-4-(4-phenoxyphenylethylamino)quinazoline | chemical reagent | | 0.00598 |
| NEO1 | transcription regulator | | 0.00598 |
| BQ 123 | biologic drug | | 0.00598 |
| NEUROD2 | transcription regulator | | 0.00598 |
| TCF21 | transcription regulator | | 0.00598 |
| STAP2 | other | | 0.00716 |
| BI 811283 | chemical drug | | 0.00784 |
| GREM1 | other | | 0.00826 |
| CDH2 | other | | 0.00826 |
| p38 MAP kinase inhibitor | chemical drug | | 0.0111 |
| CHI3L1 | enzyme | | 0.0111 |
| D-2-amino-5-phosphonovaleric acid | chemical reagent | | 0.0111 |
| polymethyl methacrylate | chemical reagent | | 0.0111 |
| LSP1 | other | | 0.0111 |
| SCIN | other | | 0.0111 |
| ERO1A | enzyme | | 0.0111 |
| HSD17B1 | enzyme | | 0.0111 |
| OLFM2 | other | | 0.0111 |
| Cmtm2a | transcription regulator | | 0.0111 |
| IL17B | cytokine | | 0.0111 |
| ADAMTS1 | peptidase | | 0.0111 |
| S100A11 | other | | 0.0111 |
| Bc1 | translation regulator | | 0.0111 |
| ULK1 | kinase | | 0.0111 |
| Hyaluronidase | biologic drug | | 0.0111 |
| miR-515-5p (and other miRNAs w/seed UCUCCAA) | mature microRNA | | 0.0111 |
| PTK6 | kinase | | 0.0111 |
| AMOTL2 | other | | 0.0111 |
| emetine | chemical toxicant | | 0.0111 |
| ARID2 | transcription regulator | | 0.0111 |
| SLCO1C1 | transporter | | 0.0111 |
| MS645 | chemical reagent | | 0.0111 |
| leucine-2-alanine enkephalin | chemical drug | | 0.0111 |
| GR-MD-02 | chemical drug | | 0.0111 |
| SPINT2 | other | | 0.0142 |
| EBF2 | transcription regulator | Activated | 0.0149 |
| quinethazone | chemical drug | | 0.0178 |
| GPR17 | G-protein coupled receptor | | 0.018 |
| Mamld1 | other | | 0.018 |
| tranexamic acid | chemical drug | | 0.018 |
